# Ontogeny of the spinal cord dorsal horn

**DOI:** 10.1101/2025.03.14.643370

**Authors:** Robert Brian Roome, Archana Yadav, Lydia Flores, Amrit Puarr, Diana Nardini, Alexander Richardson, Ronald R. Waclaw, Ruth M. Arkell, Vilas Menon, Jane E. Johnson, Ariel J. Levine

## Abstract

The dorsal horn of the mammalian spinal cord is an exquisite example of form serving function. It is comprised of diverse neuronal populations stacked into laminae, each of which receives different circuit connections and plays specialized roles in behavior. An outstanding question is how this organization emerges during development from an apparently homogeneous pool of neural progenitors. Here, we found that dorsal neurons are diversified by time, with families of related cell types born as temporal cohorts, and by a spatial-molecular gradient that specifies the full array of individual cell types. Excitatory dorsal neurons then settle in a chronotopic arrangement that transforms their progressive birthdates into anatomical order. This establishes the dorsal horn laminae, as these neurons are also required for spatial organization of inhibitory neurons and sensory axons. This work reveals essential ontogenetic principles that shape dorsal progenitors into the diverse cell types and architecture that subserve sensorimotor behavior.

**Highlights:**

- Temporal cohorts of late-born dorsal neurons give rise to neuronal families
- Sequentially-born excitatory neuron families form adjacent laminae
- Laminar structure specifically requires excitatory (but not inhibitory or sensory) neurons
- Graded expression of Zic transcription factors directs fine neuronal identity

## Introduction

The dorsal horn of the spinal cord is crucial for some of the most important neural tasks of vertebrate animals, from processing pain and touch signals from the body to implementing sensorimotor control. It relies on a highly structured organization in which dozens of molecularly discrete excitatory and inhibitory neuronal cell types are arrayed within anatomical layers, or laminae^1–4^. Each cell type has a characteristic transcriptional profile and “families” of closely related cell types are highly conserved across mammalian species^5–7^. Each cell type also has a characteristic spatial distribution that is a strong predictor of its synaptic inputs and function, as the sensory and descending pathways that target the dorsal horn each terminate in stereotyped laminar zones^1,7–11^. For example, neurons in lamina I and II process pain, itch, and temperature^8,12–17^, those in lamina III and IV process light touch^8,10,18–21^, and those in lamina V and VI process the sense of body-position and contribute to motor control^9,22–31^. Overall, the arrangement of diverse cell types in laminae is a hallmark of the dorsal horn and the framework that supports normal behavioral function.

Despite our detailed knowledge of these cell types and structure, there is an enduring mystery in our understanding of the dorsal horn of the spinal cord: we still do not know how all this diversity and anatomical organization comes to be. This is in sharp contrast with development of the ventral horn – a textbook example of developmental logic^32,33^. The ventral horn is mainly formed during an early phase of spinal neurogenesis, when secreted ligands spatially pattern the progenitors into eleven discrete lineage domains that each give rise to a “cardinal class” of interneurons^34–43^. Then, as neurogenesis proceeds, something surprising happens among a small group of late dorsal progenitors: they lose their early lineage restriction, expand dramatically, and form a large common pool of pdL progenitors (<u>p</u>rogenitors of <u>d</u>orsal late) that generate the inhibitory (dILA; <u>d</u>orsal interneuron late) and excitatory (dILB) neurons that comprise the dorsal spinal cord^35,36,44–47^. How does this seemingly homogeneous population of progenitors generate the highly diverse and highly structured cellular organization of the dorsal horn?

Here, we examined mouse dorsal horn development from neurogenesis until the establishment of laminar structure using neuronal birthdating, transcriptional profiling, spatial transcriptomics, and mouse mutant analysis. We found that the families of the mature dorsal horn neuron types emerged as progressive waves of dIL neurons during late spinal neurogenesis. The excitatory families then settled into a chronotopic order, with progressively born families settling into adjacent lamina. To test the hypothesis that dorsal excitatory neurons establish the laminar structure of the dorsal horn, we genetically eliminated dorsal excitatory, dorsal inhibitory, or sensory neurons and found that dorsal excitatory neurons specifically were crucial for the proper anatomical organization of the dorsal horn. We also discovered a spatial gradient of Zic transcription factors in dorsal progenitors and post-mitotic dIL neurons that sub-divided each family into more refined cell types, generating an impressive array of approximately 150 molecularly discrete types. Together, this work uncovered the underlying developmental mechanisms that transform neural progenitors into the dorsal horn of the spinal cord and thereby establish the neural substrates for sensorimotor control.

## Results

### A transcriptional atlas of the late embryonic dorsal horn

Our understanding of cell type organization and development has exploded recently due to advances in single cell profiling that enable powerful new descriptive studies. We and others have applied these methods to study spinal cord biology in many different contexts, but there is a persistent gap in our knowledge: the crucial period in which dorsal spinal neurons differentiate from relatively generic immature neurons into the diverse cell types of the dorsal horn. Describing this period in detail should therefore provide insights into the processes of neuronal diversification, how cell types relate to each other, and laminar development.

To examine the full complement of dorsal horn neurons, we needed to define the earliest period after which all neurons have been born; accordingly, we charted a detailed timecourse of spinal neurogenesis. EdU, which is incorporated into the DNA of dividing cells, was injected into female mice with timed pregnancies at half-day timepoints from <u>e</u>mbryonic day (E) E10.5 through E13.5, covering the general period of spinal cord neurogenesis. At E16.5, when the dorsal horn’s laminar structure is evident^48^, we analyzed the distribution of EdU+ neurons, including both dorsal inhibitory neurons (marked by Pax2) and dorsal excitatory neurons (marked by Lmx1b) (Figure 1A). Nearly all neurons that settled in the dorsal horn were born between E11.0 and E13.0 (Figure 1B), and neurogenesis was essentially complete by E13.5, consistent with previous work^36,37,49–51^. Based on these results, we generated a single-cell RNA-sequencing dataset of spinal cords at E14.5 and E16.5 (Figure S1A). We profiled 195,146 spinal cord cells (Figure S1, Tables S1 and S2) and obtained 94,048 neurons (Figure 1C). Hierarchical clustering was used to reveal multiple levels of spinal neuron organization. At the highest level, neurons were divided into three major groups that each accounted for approximately one third of the dataset.

**Figure 1:**
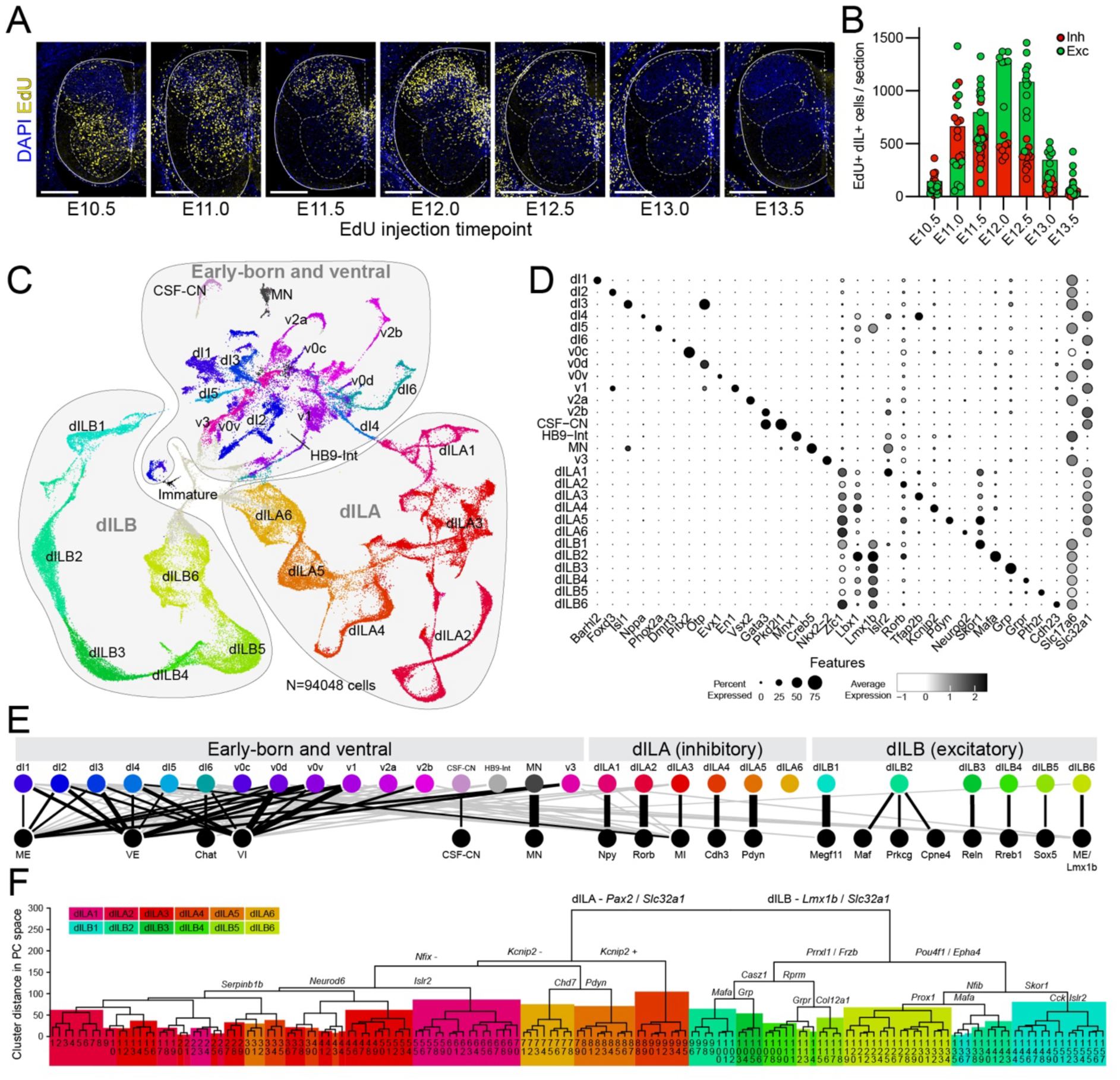
A single-cell transcriptomic atlas of the mouse spinal cord post-neurogenesis. (A) Neuronal birthdate labeling in spinal cord sections from E16.5 embryos that received an EdU pulse (yellow) to mark dividing cells at the indicated timepoints between E10.5 and E13.5, with DAPI staining (blue). (B) Neuron counts per section of inhibitory (Pax2+, red) or excitatory (Lmx1b+, green) neurons that were EdU positive following EdU pulses at the indicated timepoints. (C) UMAP plot of single-cell RNA-sequencing data taken from E14.5 and E16.5 wild-type mouse embryos showing all neurons, and annotated by Group (1 = Early-born and ventral, 2 = dILA, 3 = dILB) and by neuronal classes – including six dILA and six dILB families, each shown in a distinct color. (D) Dot plot showing gene expression in each neuron class. Dot size indicates the percent of each cluster expressing each gene and dot shade indicates the average expression level of each gene in cells of each cluster. (E) Diagram illustrating the relationships between embryonic neuron classes and postnatal/adult neuron classes from Russ et al., (2021)^7^. Line thickness is inversely proportional to centroid-centroid distance in principal component (PC) space, with the strongest relationships emphasized with black (versus gray) lines. (F) A dendrogram of centroid-centroid distance in PC space for refined neuron types among all dIL neurons. Each terminal branch number corresponds to named cell-types in Figure S3. Selected branch points are annotated with top DEGs. Where branches encompass large parts of dIL families they are shaded with colors representing each dIL family (see legend). Numbers: (A,B) EdU birthdating data was derived from 79 E16.5 mouse embryos given EdU on E10.5 (n=13 from four litters), E11.0 (n=10 from three litters), E11.5 (n=15 from four litters), E12.0 (n=7 from three litters), E12.5 (n=12 from four litters), E13.0 (n=10 from three litters) or E13.5 (n=12 from four litters). (C-F) Single-cell RNA-sequencing data was derived from fifteen litters of mouse embryos (n=3 E14.5, n=12 E16.5), and of which data from N=94048 neurons were extracted. Scale bars: (A) 250 μm Abbreviations: DEG – differentially-expressed gene, Exc – excitatory, Inh – inhibitory, UMAP – uniform manifold approximation and projection.

<u>Group 1</u> was comprised of the cardinal spinal cord neuron types including dI1, dI2, dI3, dI4, dI5, dI6, v0c, v0d, v0v, v1, v2a, v2b, and v3 interneurons, motoneurons, as well as Hb9 interneurons and cerebrospinal fluid contacting neurons. Semi-supervised clustering yielded the full set of expected spinal lineage classes, which we identified by expression of classic markers (Figure 1C,D, Tables S3 and S4) and by correlation with a well-annotated atlas of the period of spinal neurogenesis (E9.5 – E13.5) by Delile and colleagues (see Methods)^42,52,53^.

<u>Groups 2 and 3</u> represented the putative dILA neurons (Group 2) and dILB neurons (Group 3), based on the expression of general embryonic dorsal markers (shared with putative dI4 and dI5 classes), modest residual expression of early post-mitotic markers (suggesting recent neurogenesis), later-born temporal transcription factors, as well as the expression of known markers of mature dorsal neurons, the region where dILA/B neurons reside^7,35,36,52^ (Figure 1C,D). Given that late-born dILA and dILB neurons share some gene expression with their early-born dI4 and dI5 neurons counterparts, we sought distinguishing characteristics between them to justify a point of demarcation in our dataset. Exploratory analysis revealed that the transcription factor Zic1 was expressed in putative dILA and dILB neurons, but not dI4 and dI5 neurons. Zic1 is expressed in the dorsal progenitor zone throughout neurogenesis and also appears in post- mitotic neurons around E11.5, which aligns with the emergence of the first dIL neurons^54,55^.

Immunofluorescence confirmed this pattern and showed that Zic1 protein expression in neurons was coincident with the dramatic dorso-ventral expansion of the Lbx1-producing progenitors at the onset of dIL neuron production (Figure S2). In transcriptomic data, Zic1 expression was highly enriched in Group 2 and Group 3 relative to dI4 and dI5 clusters and similarly divided early and late “dI4” and “dI5” neurons in a single cell atlas from the period of active spinal neurogenesis^52^ (Figure S2). Accordingly, we used Zic1 expression to delineate the boundaries between dI4-dILA and dI5-dILB (see Methods).

The internal organization of dILA and dILB neurons has not been analyzed in depth before, limiting our understanding of how dIL neurons diversify into the dozens of cell types in the mature dorsal horn. Sub-clustering of dILA and dILB yielded six classes within each Group that we designated dILA1, dILA2, dILA3, dILA4, dILA5, dILA6, dILB1, dILB2, dILB3, dILB4, dILB5, and dILB6 (Figure 1C,D and Figure S3,). Each of these classes were robust and well-separated, with high silhouette values, very low LISI scores, and high discrimination accuracy and precision with a random forest classifier (Figure S3-4, Tables S3 and S4). Comparison of these dILA and dILB classes with a harmonized transcriptomic atlas of postnatal spinal neurons^7^ revealed that each dILA or dILB class correlated strongly with a single “family” or a small group of related families (Figure 1E, Table S5), with the exception of dILA6 which did not have a clear postnatal counterpart. Thus, at E14.5 and E16.5, young dILA and dILB neurons were already well organized as families, the intermediate level of cell type hierarchy.

To examine the most refined level of cell type organization hierarchy, that of specific cell types, we sub-clustered each family and found approximately 150 molecularly distinguishable clusters (Figure 1F and Figure S3-5, Tables S6-S8). To determine whether refined dIL neuron types were segmentally restricted, we compared the cell type proportions in samples derived from cervical, thoracic, and lumbar segments. We found that refined dIL neuron-types were largely present at all segments, with a few modest differences (Figure S3).

This work provided the first comprehensive characterization of embryonic spinal neurons post-neurogenesis, allowing us to link the transcriptomic profiles of progenitor lineage domains with their differentiated progeny. In particular, we found that both the family and refined type organization of the mature dorsal horn are already present shortly after neurogenesis.

### Temporal diversification of embryonic dILA and dILB families

What processes diversify pdL progenitors into the dILA/B families? During the early wave of spinal neurogenesis, morphogen-directed spatial patterning is the dominant driver that creates the set of canonical lineage classes^56–59^. However, spatially restricted domains within the pdL progenitor pool have not been found, raising the possibility that a distinct logic governs the development of the late-born dorsal neurons. Based on analogy with other laminated neural structures including the vertebrate retina, cortical plate, and tectum, we hypothesized that temporal patterning may govern the cell type diversification and anatomical layering of the dorsal horn^60–64^.

We first noted that dILA and dILB families were organized as streams in a UMAP plot, with one end closest to early-born Zfhx3+ dI4 and dI5 counterparts^50^ and the other end closest to Robo3+ immature/migrating neurons^65–69^ and Sox2+ progenitors^70^ (Figure 2A), suggestive of continuous temporal variation. A complementary dimensionality reduction and visualization approach, PHATE, revealed a similar pattern (Figure 2B)^71^. Such inter-family relationships were further supported by the observation that the rare mistakes made by a classifier trained on family identity were between families that were adjacent in UMAP and PHATE plots (Figure S5). These exploratory findings prompted us to test whether families are organized temporally, which we did using three independent approaches.

**Figure 2:**
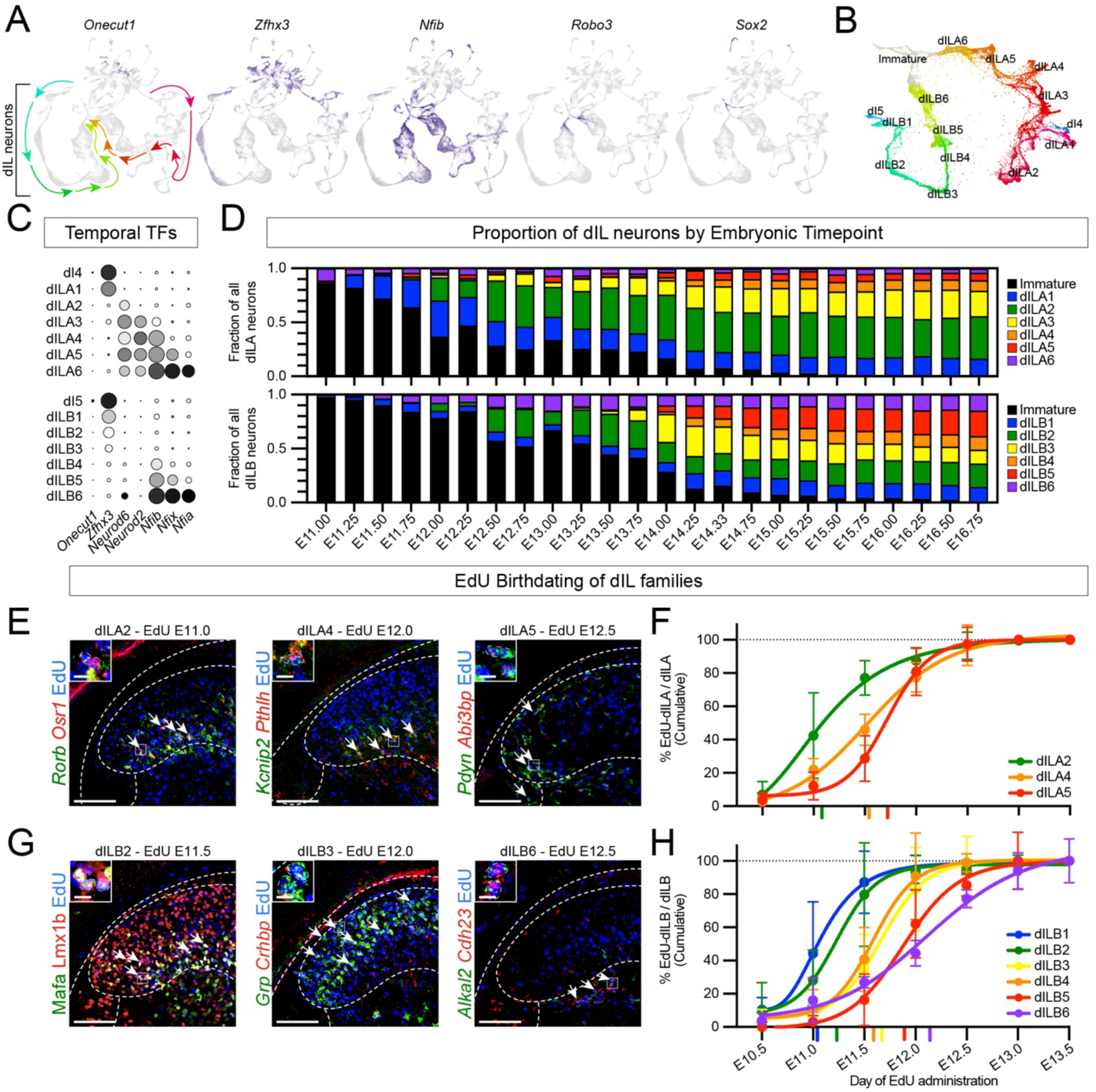
dIL families represent temporal cohorts. (A) UMAP feature plots of gene expression of each indicated gene in single-cell RNA-sequencing data, including temporal transcription factors (*Onecut1, Z+x3, Nfib*), a marker of migrating and commissural neurons (*Robo3*) and a marker of neural progenitors (*Sox2*). (B) PHATE plot of dIL neurons, together with their early-born dI4/5 counterparts and immature neurons. (C) Dot plot of temporal transcription factors among dIL families, with dot size representing the percent of cells in each type expressing the gene and shade representing average expression level. (D) Composition over time of dILA families (top) and dILB families (bottom), together with immature dILA/B neurons (black) in single-nucleus RNA-sequencing data from precisely-timed mouse embryo dissections^72^. (E-H) EdU birthdating of dILA and dILB families. (E,G) Examples of dILA (E) and dILB (G) family neurons (indicated by marker RNA or protein in green and red) co-labeled with EdU (blue) injected at the indicated timepoints. White arrows indicate examples of dIL family subtypes co-labeled with EdU. Dashed white lines indicate approximate anatomical boundaries of the spinal cord, including the pial outer boundary, the gray/white boundary, and the superficial dorsal horn. Boxes in dashed-white lines indicate the field of view magnified in the inset. (F,H) Cumulative percent of each dILA (F) or dILB (H) family co-labeling with EdU pulsed at the indicated embryonic time point (x-axis) showing mean +/- standard deviation for all litters at each timepoint, connected by sigmoidal fit lines for each family. The imputed 50% cumulative birthdating timepoint for each family is indicated by a colored vertical line beneath the x-axis. Numbers: (A-C) Single-cell RNA-sequencing data is the same as Figure 1C (see legend). (D) Enriched spinal cord neuron single-nucleus RNA-sequencing data (N=349,306 neurons) was retrieved from Qiu et al., (2024) and refined cell type labels were transferred from our data (Figure 1C- F); from this, 74,764 dILA and 71,834 dILB neurons were analyzed across 24 time points. (E-H) EdU birthdating data was derived from 34 E16.5 mouse litters from a separate cohort to Figure 1A,B, and data from individual embryos per litter were averaged. Pregnant mothers were given EdU on E10.5 (n=3-5 litters), E11.0 (n=4 litters), E11.5 (n=5 litters), E12.0 (n=5 litters), E12.5 (n=4-5 litters), E13.0 (n=5 litters) or E13.5 (n=2-5 litters). Scale bars: (E) 250 μm, 25 μm for insets. Abbreviations: TTF – temporal transcription factor

First, we assessed the expression of the temporal transcription factors Onecut, Zfhx, Nfi, and Neurod genes in each dILA and dILB family (Figure 2C). The Onecut genes, that often mark the earliest-born cell types, were not present in the dIL families, but we observed a smooth progression of Zfhx (mid early-born), Neurod (mid late-born), and Nfi (latest-born) genes from dILA1/B1 through dILA6/B6. Therefore, both the inhibitory and excitatory families of the dorsal horn vary by a combinatorial code of temporal transcription factors.

Second, we reasoned that molecularly distinct dILA and dILB families should appear in a progressive order if they are born successively. Recent work from Qiu et al. provided a large embryonic single cell atlas with high temporal resolution, with embryos staged and sorted based on refined anatomical criteria into quarter-day timepoints^72^. We extracted spinal cord neurons from this dataset, used label transfer to assign family identity, and then analyzed family composition over time. Amongst dILA families, the main order was dILA1 > dILA2 > dILA3 > dIL4 and dILA5, while dILA6 was detected throughout (Figure 2D). Amongst dILB families, the main order was dILB1 > dILB2 > dILB3 > dILB5 > dILB4, while dILB6 was detected throughout (Figure 2D). The apparent early appearance of dILA6 and dILB6 may be accurate or may reflect the technical caveat that late-born neurons in these data resemble immature neurons. Overall, these data further support the conclusion that dILA and dILB families represent temporal cohorts.

Third, we assessed dILA and dILB birthdates using timed EdU administration over half-day intervals combined with expression of family marker genes. This method measures birthdate more directly, though it provides slightly lower temporal resolution due to imprecision in timing of conception relative to injection time and experimental analysis^73,74^. Amongst dILA families, we performed targeted analysis of three families for which either a single marker or a combination of two markers was sufficient to identify constituent neurons unambiguously in tissue sections (Figure 2E). dILA2, dILA4, and dILA5 were born in overlapping waves of neurogenesis with dILA2 born at E11.0-11.5, and dILA4 and dILA5 born around E11.5-12.0 (Figure 2F and Figure S6), with dILA4 preceding dILA5 slightly. Amongst dILB families, EdU birthdating was performed for all six families, or subsets thereof (Figure 2G). We found that the main birthdates for each group were as follows: dILB1 at E11.0-11.5, dILB2 at E11.5, dILB3 and dILB4 at E11.5-12.0, dILB5 at E12.0-12.5, and dILB6 at E12.0-13.5 (Figure 2H, Figure S7,8). These data showed an orderly progression of family birthdate.

From this, we conclude that families of dorsal horn neurons are born progressively across the period of late neurogenesis, and that temporal diversification is a major organizing principle of the dorsal horn.

### A chronotopic map of dILB families establishes the laminar structure of the dorsal horn

A common theme of neural development, particularly in laminated structures, is chronotopy: an organizational principle in which progressively born neuronal subsets settle into ordered anatomical arrangements, providing the underlying structure for mature circuit function^60–64,75,76^. To explore whether successively born dILA or dILB neurons could account for the laminar structure of the dorsal horn, we examined the anatomical location of all dorsal inhibitory (Pax2+) and dorsal excitatory (Lmx1b+) neurons with respect to neuronal birthdate (EdU+). We found that dILA neurons born at particular times had stereotypical spatial distributions, but often spanned many laminae and had no spatially congruent pattern across timepoints (Figure 3A,B). In contrast, we observed that progressively born Lmx1b+ neurons settled into a coherent order: putative dI5 neurons, born at E10.5^35,36,77^, were found in lateral lamina V, followed by dILB neurons born at E11.0/E11.5 in lamina III/IV, those born at E12.0/E12.5 in lamina I/II, and those born at E13.0/E13.5 in part of lamina IIi/III and medial lamina V (Figure 3A,B).

**Figure 3:**
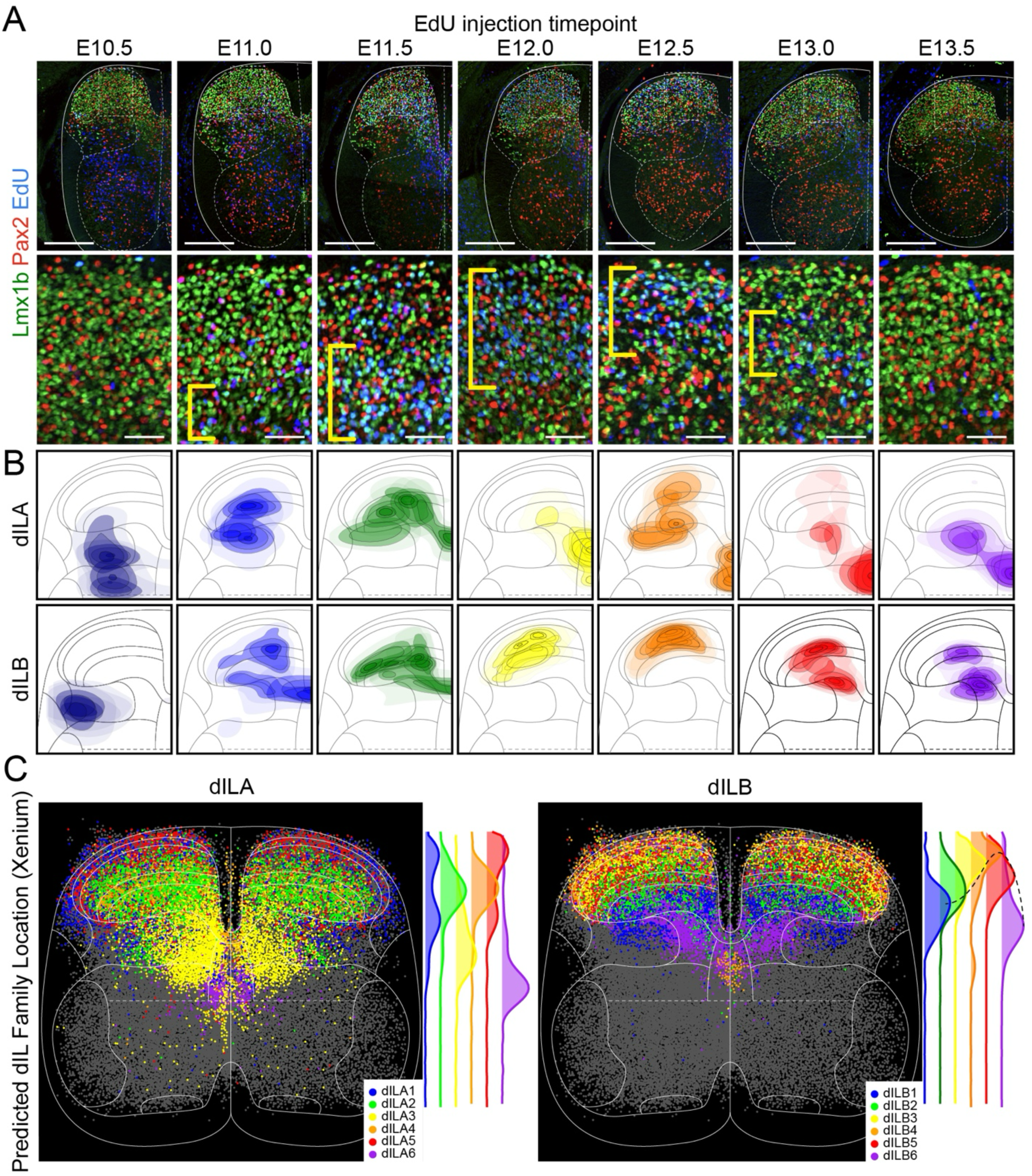
dILB neurons settle into a chronotopic map. (A) Images of embryonic spinal cord sections taken from separate E16.5 embryos that received EdU (blue) pulses at the indicated timepoints between E10.5 and E13.5, with immuno-labeling for the excitatory marker Lmx1b (green) and the inhibitory marker Pax2 (red). Hemisections (top row) and magnifications (bottom row) are shown, with the approximate region of Lmx1b+/EdU+ nuclei indicated with brackets (yellow). Solid and dashed white lines indicate approximate anatomical boundaries of the spinal cord. (B) Density plots of coordinates for birth-dated dILA neurons (Pax2+/EdU+) or dILB neurons (Lmx1b+/EdU+), for EdU pulse experiments at each of the indicated timepoints. Separate perimeters were generated for each replicate per timepoint and for five proportions of labeled cells (50%, 25%, 15%, 5%, 2.5%), and were colored in order of decreasing opacity. Replicates consist of all embryos of a single litter labeled with EdU at the indicated timepoint. (C) Scatter plots of neurons in data from Xenium spatial transcriptomics, annotated based on predicted dIL family identity. Adjacent histograms demonstrate the frequency of neurons belonging to dILA or dILB families by depth. Probe sets used were the mouse brain panel v1.1 (248 genes) plus 100 custom genes based on single-cell RNA-sequencing generated here (Table S9). Numbers: (A,B) n=79 embryos, 7-15 per timepoint. Note that data is reprised from Figure 1 (A,B), see legend for details. (C) For spatial transcriptomics, cell coordinates were derived from six whole sections and seven hemisections (N=9.5 sections) among n=3 replicate wild-type E16.5 spinal cords. From these sections 54,166 cells were identified as neurons and plotted, of which 27,714 were annotated as dIL neurons. Scale bars: (A) 250 μm (top row), 50 μm (bottom row).

We next assessed the spatial distribution of each dILA/B family to determine whether the particular families followed the pattern of the overall Pax2+ and Lmx1b+ neurons. Xenium spatial transcriptomics was used to profile gene expression in tissue sections from E16.5 mouse spinal cords (n=3) and predicted cell identities^78^ were mapped onto normalized cell coordinates from all replicates. As predicted by the EdU birthdating, we found that dILA families settled into stereotypical locations that lacked an overarching temporal-spatial order, while progressively born dILB families were found in progressively adjacent laminae (Figure 3C, S9). The highest density of the first-born dILB1 family was found in lamina IV-V, dILB2 was in lamina III-IV, dILB3 was in lamina II-III, and dILB4 was in lamina IIo. The progression then reversed direction, with dILB5 settling broadly throughout lamina II-III and dILB6 in medial lamina V as well as a streak along dorsal lamina V. The arrangement of dILA and dILB families corresponded to the progression of Pax2 and Lmx1b-birthdated neurons across neurogenesis.

The chronotopic development of dILB families raised the possibility that excitatory neuron organization is the critical determinant of laminar structure in the dorsal horn. To test this hypothesis, we used mouse genetics to selectively remove each of the major classes of neurons from the dorsal horn: dorsal inhibitory neurons, dorsal excitatory neurons, and sensory neurons whose cell bodies are in the dorsal root ganglion but whose axons innervate the dorsal horn.

We then compared the impact of each perturbation on the distribution of the remaining neural elements. To remove dorsal inhibitory neurons from the spinal cord, we used Ptf1a^Cre/Cre^ homozygous null embryos, in which the dI4/dILA neurons are trans-fated to become dI5/dILB neurons^46,79^. As expected, Pax2 expression was nearly absent from the dorsal horn, but the overall morphology of the tissue appeared normal (Figure 4A). In situ hybridization and immunofluoresence for markers of dILB and sensory neuron types revealed that their typical spatial distributions were preserved (Figure 4A,C and Figure S10). Spatial transcriptomics performed on Ptf1a^Cre/Cre^ tissue also revealed that predicted cell type locations for dILB families were normal (Figure 4E and Figure S11-12). Thus, inhibitory neurons were not required for the anatomical organization of the dorsal horn.

**Figure 4:**
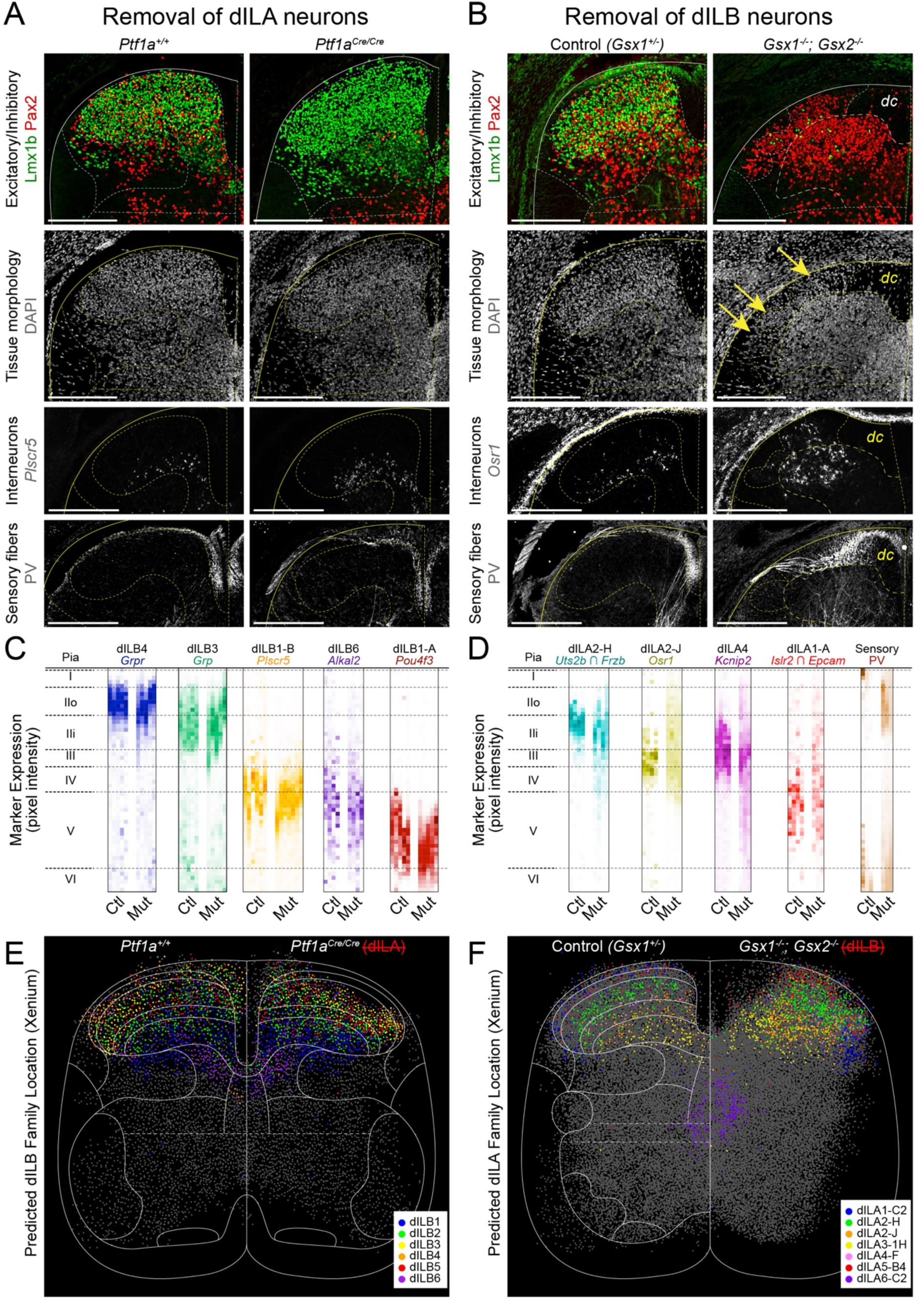
dILB neurons are required for laminar organization of the dorsal horn. (A,B) Images of E16.5 spinal cord dorsal horn hemi-sections from control and mutant embryos. (A) Sections from *Ptf1a^+/+^* and *Ptf1a^Cre/Cre^* embryos showing loss of Pax2-immunolabeled neurons (red), tissue morphology (DAPI, gray), an example marker of an excitatory interneuron (*Plscr5* RNA, gray), and proprioceptive sensory afferents (PV antibody, gray). (B) Sections from *Gsx1^+/-^*; *Gsx2^+/+^* and *Gsx1^-/-^*; *Gsx2^-/-^* embryos showing loss of Lmx1b-immunolabeled neurons (green), tissue morphology (DAPI, gray), an example marker of an inhibitory interneuron (*Osr1* RNA, gray), and proprioceptive sensory fibers (PV antibody, gray). (C,D) Spatial distribution of individual gene or protein markers or marker sets (as indicated at the top of each box), showing the normalized proportion of total signal per dorso-ventral bin (color intensity) where the pia is set as the dorsal boundary and approximate laminar positions are shown, for either (C) absence of dILA or (D) absence of dILB neurons. Biological replicates are grouped together as individual columns for either Ctl (control – *Ptf1a^+/+^* or *Gsx1^+/-^*; *Gsx2^+/+^*) or Mut (*Ptf1a^Cre/Cre^* or *Gsx1^-/-^*; *Gsx2^-/-^*) embryos. (E,F) Scatter plots of neurons in data from Xenium spatial transcriptomics, annotated based on predicted dIL family identity. (E) Brachial control *Ptf1a^+/+^* embryo sections (n=3, left side) were compared to a single *Ptf1a^Cre/Cre^* embryo section (n=1, right side). The control sections are also displayed in Figure S10. (F) Thoracic control *Gsx1^+/+^*; *Gsx2^+/+^* embryo sections (n=3, left side) were compared to *Gsx1^-/-^*; *Gsx2^-/-^* embryo sections (n=3, right side). Control neurons were downsampled to match the number of mutant cells displayed. Cells not belonging to the indicated dIL cell type are plotted in gray. Solid and dashed white and yellow lines indicate approximate anatomical boundaries of the spinal cord. Numbers: (A,D) Representative images were derived from n=4-6 *Ptf1a^+/+^*, n=3-7 *Ptf1a^Cre/Cre^*, n=3-4 *Gsx1^+/-^*; *Gsx2^+/+^* and n=3-4 *Gsx1^-/-^*; *Gsx2^-/-^* and histograms were derived from these same embryos (different numbers within each range were analyzed for each marker set depending on staining quality). (E,F) Spatial transcriptomics from E16.5 wild-type embryo brachial segments was previously described (Fig. S9). For E16.5 wild-type thoracic segments, cell coordinates were derived from five whole sections and one hemisection (N=5.5 sections), among n=3 replicate spinal cords. From these sections 27,758 cells were identified as neurons and plotted, of which 13,724 were annotated as dIL neurons. For E16.5 *Ptf1a*- null brachial segments, cell coordinates were derived from two whole sections (N=2 sections) among n=1 spinal cord. From these sections, 8,267 cells were identified as neurons, and of which 2,767 were dIL neurons. E16.5 brachial data was downsampled to 8,106 neurons of which 4,151 were dIL neurons to match. For E16.5 *Gsx1/2*-null thoracic segments, cell coordinates were derived from four whole sections and three hemisections (N=5.5 sections) among n=3 replicate spinal cords. From these sections, 24,737 cells were identified as neurons, and of which 12,217 were dIL neurons. E16.5 thoracic data was downsampled to 24,583 neurons with, also, 12,217 dIL neurons, to match. Scale bars: 250 μm Abbreviations: ctl – control, dc – dorsal columns, mut – mutant, PV - parvalbumin.

To remove dorsal excitatory neurons from the spinal cord, we used Gsx1^-/-^; Gsx2^-/-^ double mutant embryos, in which dI5/dILB neurons are trans-fated to become dI4/dILA neurons^45,80,81^. In Gsx double mutants, Lmx1b was nearly absent from the dorsal horn at E16.5, as expected, and the remaining tissue displayed abnormal morphology (Figure 4B).

There was a striking spatial disorganization of the remaining neural components of the dorsal horn. The restricted laminar distributions of dILA neuron types found in wild-type embryos were lost in double mutants, with some neuron types present in a mildly expanded range (Uts2b+/Frzb+ dILA2-H neurons) and others being scattered from the pia all the way to the base of the dorsal horn (Osr1+ dILA2-J neurons and Kcnip2+ dILA4 neurons) (Figure 4B,D,F and Figure S13-14). Despite this disorganization, dILA types destined for the same lamina were often found together in clumps, which may reflect cell-type intrinsic sorting mechanisms. Sensory fibers were similarly disorganized, with PV+ large proprioceptor fibers crossing through the superficial dorsal horn (Figure 4B,D). We also observed neurons scattered amongst the dense longitudinal sensory fibers just below the pia, reflecting an abnormal boundary between the gray and white matter (Figure 4B).

We also considered whether laminar organization could be imposed on dIL neurons by an extrinsic source. At E16.5, the only other neural element in the dorsal horn are sensory fibers, as descending axons from the brain have not yet penetrated the dorsal gray matter^82–90^. To ablate sensory neurons, we used Pax3^Cre^;Isl2^lsl-DTA^ embryos^91,92^, which express diphtheria-toxin in sensory neurons (and in some other neural crest-derivatives). As expected, this ablation led to loss of dorsal root ganglia and of sensory axons in the gray matter (Figure S15). In addition, Lissauer’s tract and the dorsal columns were reduced dramatically in size and the dorsal horns often fused at the midline. Despite these changes, dILA and dILB neuron markers were present and displayed laminar patterns in the same dorso-ventral order as control spinal cords (Figure S15) consistent with a prior report^93^, indicating that sensory neuron innervation was not required for lamination.

Together, these data suggest a hierarchy of structure in the dorsal horn, in which excitatory neurons form laminar structures organized by birthdate and that inhibitory and sensory axons are dependent on this process to achieve their normal laminar distributions.

### Spatial-molecular diversification of dIL neuron families generates refined cell types

The temporal diversification process that generated the six dILA and six dILB families is unlikely explain the impressive molecular diversity of the ∼150 refined dorsal spinal neuron types, so we next asked what other sources of variation may instruct dorsal neuron identity. To nominate candidate factors, we explored the most highly weighted genes for the top principal components explaining molecular variation within each dIL family (Table S10). This revealed that Zic transcription factors contributed highly to variation in nearly all families. In addition to Zic1, which we showed is expressed in post-mitotic dIL neurons (Figure S2), there are four other mammalian members of the Zic zinc finger transcription factor family, Zic2, Zic3, Zic4, and Zic5, all of which have been shown to be present in the embryonic spinal dorsal horn, particularly within the progenitor domain^54,55,94–96^. Zics can interact with the transcription factors that are downstream of Shh, TGF-beta, and Wnt signaling – three major pathways that pattern the embryonic spinal cord^97–102^. This led us to hypothesize that Zic factors may regulate heterogeneity amongst pdL progenitors and the identity of refined neuron types in the dorsal spinal cord.

We first assessed whether Zic gene expression in dILA and dILB post-mitotic neurons was related to the identity of refined neuron types. All five Zic genes were detected in UMAP feature plots amongst dILA and dILB neurons (Figure 5A) and we observed corresponding expression of all five Zic factors in the dorsal spinal cord of E16.5 embryonic tissue sections (Figure S16).

**Figure 5:**
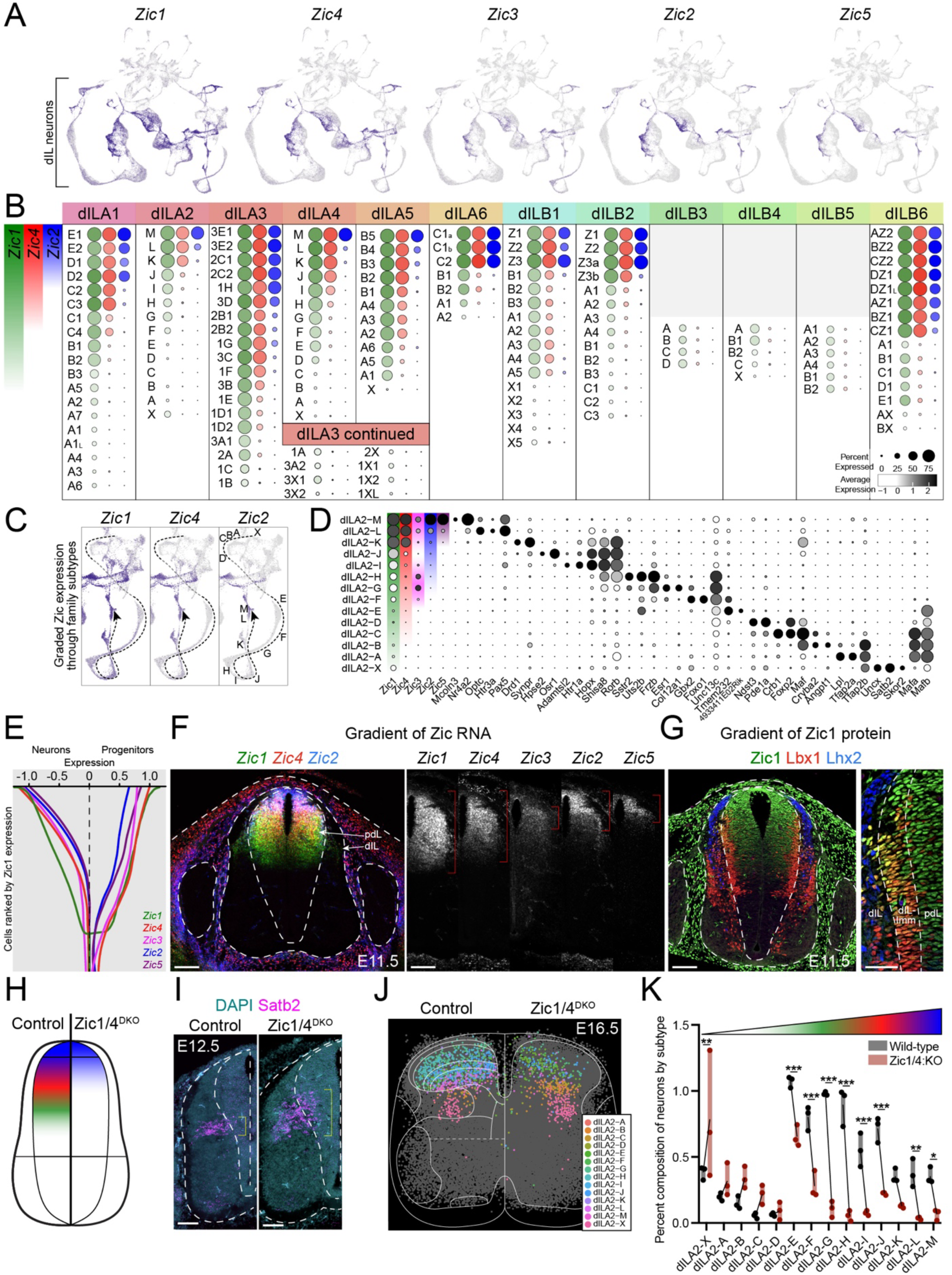
A progenitor gradient of Zic transcription factors patterns dIL neurons. (A) UMAP feature plots of *Zic1-5* expression in E14.5/E16.5 single cell data. (B) Dot plots of *Zic1* (green), *Zic4* (red), and *Zic2* (blue) RNA expression in refined cell types within each dIL family, with dot size representing the percent of cells in each type expressing the gene and shade representing average expression level. (C) Gradients of Zic gene expression align with UMAP structure, within the dILA2 refined types M-X. (D) Dot plot of dILA2 neuron subtypes, showing unique markers for each type as well as groups of gene expression which co-vary along the Zic axis. (E) dIL neurons (left) and progenitors (right) on the y-axis, ranked by *Zic1* expression (x-axis), showing expression of each Zic gene. (F) Images of E11.5 spinal cord sections with in situ hybridization for Zic genes (left, merged; right single genes). (G) Image of E11.5 spinal cord section with immuno-labeling for Zic1 (green), a marker of newborn immature and migrating dIL neurons (Lbx1, red), and with Lhx2 (dI1, blue) shown for orientation. The ventricular zone is shown as the most medial area bounded by dashed white lines. Right panel shows higher magnification inset of newborn dIL neurons as they exit the ventricular zone. (H) Genetic strategy for perturbing the Zic gradient through double knockout (DKO) alleles of *Zic1* and *Zic4*. (I) E12.5 spinal cords from control and *Zic1/4*^DKO^ embryos, showing in-situ hybridization for *Satb2* (magenta) and DAPI (cyan). (J) Scatter plots of normalized coordinates of neurons identified in Xenium spatial transcriptomics data, annotated to show dILA2 refined subtypes in control brachial sections (n=3, left) and in *Zic1/4*^DKO^ sections (n=3, right). Spatial transcriptomics from E16.5 wild-type embryo brachial segments was previously described (Fig. S9). For E16.5 *Zic1/4*^DKO^ brachial segments, cell coordinates were derived from three whole sections and one hemisection (N=3.5 sections) among n=3 replicate spinal cords. From these sections, 17,505 cells were identified as neurons, and of which 7,376 were dIL neurons. E16.5 brachial data was downsampled to 17,347 neurons of which 8,852 were dIL neurons to match. (K) Quantification of the data from (J) showing the relative composition of neuron subtypes as a percentage of all neurons, including replicates and lines connecting control and mutant means. Numbers: (A-D) N=94,048 neurons from single-cell RNA-sequencing data generated here (see Figure 1C), (E) Single-cell RNA-sequencing data from dIL progenitors (N=1,963 cells)^52^ and dIL neurons (N=61,444 cells, see Figure 1C), (F-G) n=3 E11.5 wild-type embryos, (I) n=3 E12.5 wild-type and n=3 E12.5 *Zic1/4*^DKO^ embryos, (J-K) n=3 wild-type and n=3 *Zic1/4^DKO^* E16.5 mouse embryos. Statistics: (K) Two-way ANOVA (neuron subtype x genotype). For each neuron subtype in dILA2, Tukey’s multiple comparisons test was performed for each pair of genotypes, the result of which is indicated by asterisks above the plot. *<0.05, **p<0.01, ***p<0.001 Scale bars: 100 μm, except 50 μm in the magnified panel of (G) at the right. Abbreviations: DKO – double knock-out, scRNA – single-cell RNA

Within the single cell data from all dILA families and most dILB families, Zic genes were expressed in a gradient, with refined cell types varying in both how many Zic genes were present and the expression levels of each individual Zic gene (Figure 5B). Zic1 showed the most widespread expression, followed by Zic4, Zic2, and then Zic5. Zic3 was only found in restricted cell types. Focusing on one family, dILA2, the refined types were present along a trajectory in UMAP space that varied smoothly in Zic expression levels from dILA2-M Zic^High^ neurons through dILA2-X Zic^Neg^ neurons (Figure 5C). Ranking dILA2 refined types by their Zic expression revealed that, while each type expressed unique markers, there were also shared markers amongst types that were either Zic^High^ (eg. Pax5), Zic^Med^ (eg. Rorb), or Zic^Low/Neg^ (eg. Mafb) (Figure 5D). Overall, this analysis revealed that a gradient of Zic gene expression sub-divided most dIL families into refined types of dorsal spinal post-mitotic neurons.

Cell fate restriction and diversification are typically established at the progenitor stage, but pdL progenitors have not been reported to display molecular variation. Since Zic expression has been reported previously in the dorsal spinal progenitor domain, we analyzed whether Zic expression varied in this context as well. Indeed, we found that a smooth gradient of Zic expression was present in single cell transcriptomic data from both E14.5 - E16.5 neurons as well as E11.5 - E12.5 progenitors, with Zic1 and Zic4 being the most broadly expressed and Zic2 and Zic5 being the most restricted (Figure 5E).

In embryonic spinal tissue, Zic genes were present in a striking dorsal-ventral spatial gradient in the E11.5-E12.5 progenitor domain, with highest expression in the dorsal-most part of progenitor domain and decreasing expression towards the ventral spinal cord. Zic1 extended the most ventrally, followed by Zic4, Zic2, and then Zic5 which displayed the most dorsally restricted expression; Zic3 had a distinct and more restricted pattern (Figure 5F,G). The dorsal-ventral gradient of Zic1 protein expression was observed in both pdL progenitors and in post- mitotic Lbx1-expressing dILA/B neurons (Figure 5G). Thus, Zic factors were expressed in a molecular-spatial gradient within pdL progenitors and their post-mitotic progeny.

Zic expression could simply correlate with dorsal neuron diversity or it could be an important driver of the cell type repertoire. To test whether Zic factors play an instructive role in dorsal neuron diversification, we examined *Zic3* mutant embryos that have lost a spatially restricted Zic^103,104^ and *Zic1*/*Zic4* double mutant embryos (DKO) that have lost the most widely expressed Zic genes^105^.

Unlike the other Zic genes, *Zic3* was detected in a faint bipolar gradient midway through the gradient of Zic1 (Figure 5E,F). Mutation of *Zic3* through the premature stop codon of the “katun” mutant resulted in a mild increase in Zic^Med^ refined types within several families (Figure S17). This suggests that *Zic3* may act mainly as a transcriptional repressor to modify the repertoire of refined dIL neuron types.

We hypothesized that the loss of both *Zic1* and *Zic4* would have a more profound impact on dorsal neuron development, shifting the domain of Zic-expressing progenitors and neurons dorsally and expanding the domain of Zic^Neg^ progenitors and neurons. To test this prediction, we analyzed the expression of *Satb2* in spinal cords from *Zic1/4*^DKO^ embryos and control littermates. *Satb2* was a marker of a refined neuron type (dILA2-X) with expression that initiated in the transition zone near progenitors, thereby revealing the domain of origin of these newborn neurons around E12.5. As predicted, the domain of *Satb2*-expressing dILA2-X neurons was expanded significantly and shifted dorsally (Figure 5H,I). We next used spatial transcriptomics to assess the cell type composition of the dorsal horn in *Zic1/4*^DKO^ embryos and focused our analysis on dILA2 refined types. The Zic^Neg^ dILA2-X type was significantly expanded and Zic^Low^ types showed modest increases as well. Conversely, nearly all of the Zic^Med^ and Zic^High^ dILA2 types were significantly decreased upon loss of *Zic1/4* (Figure 5J,K and Figure S17-19). Similar results were observed in other dIL families (Figure S18). Thus, the Zic molecular-spatial gradient, anchored by the broadly expressed *Zic1* and *Zic4* genes, was essential for the proper development of refined dIL neuron types.

## Discussion

We sought to uncover the general principles that govern the development of the spinal cord dorsal horn in mice, focusing on how its impressive cellular diversity and layered architecture emerge from a seemingly homogeneous pool of neural progenitors. We found that dorsal spinal progenitors give rise to multiple generations of neurons that varied transcriptionally based on the time and spatial position from which they originated. Specifically, each successive wave of newborn neurons gave rise to an excitatory or inhibitory coarse “family” of related neurons and each family was further divided into refined types based on a dorsal-ventral gradient of Zic transcription factors. Following neurogenesis, the young excitatory neurons settled into an orderly chronotopic map and we found that this organization was essential to the formation of the laminae of the dorsal horn. Altogether, we charted the detailed embryonic origins of mature dorsal neurons, discovered the developmental mechanisms that shape their molecular diversity, and revealed how their anatomical organization is established, thereby laying down the neural substrates for sensorimotor control of behavior.

How is the continuous flow of developmental time converted into the various dILA and dILB families? We propose that the conserved dorsal cell type families reflect the statistical modes of cell types generated over time, rather than discrete types separated by abrupt transitions. This model is supported by the overlapping emergence of dILA and dILB families in the precisely timed single cell data; the superimposed waves of peak birthdates that we observed for each family in neuronal birthdating analysis; and the small amount of “blurring” between temporally adjacent families. In this way, dorsal horn development may be similar to the formation of the mammalian retina, in which the generation of cell types is determined by the stochastic combination of progressive changes in a transcription factor code and the particular timing of progenitor cell cycle exit^106–109^. However, to determine the progenitor mechanisms of temporal cell type diversification in the spinal cord, clonal analysis and/or new sequencing-based methods for tracking cell fate relationships over time will be needed.

Not only does developmental time guide the emergence of dorsal horn families, but we found that it is etched onto the structure of the mature spinal cord. A general theme of laminated structures in the nervous system is the presence of an anatomically organized microcircuit with precise afferent and efferent connections in each layer^110–113^. This is certainly the case for the *adult* spinal dorsal horn, particularly for sensory and descending brain axons that target specific lamina to cooperatively mediate sensorimotor control. Our data suggest that the chronotopic development of the dILB neurons establishes the spatial landscape to receive these inputs at their proper locations. Progressively born excitatory dILB neurons settled into neighboring layers and their loss profoundly disrupted the spatial organization of remaining neural elements including inhibitory neurons and sensory afferents. In contrast, loss of inhibitory dILA neurons or sensory neurons preserved normal laminar organization, while descending axons from the brain only target spinal neurons after the lamina have already been established. An autonomous role for dorsal spinal neurons in self-organizing into their characteristic layers is further supported by the intrinsic ability of cultured dorsal interneurons to form laminae upon transplantation into spinal tissue, with sensory and corticospinal axons invading these miniature dorsal structures in a spatially appropriate manner^114^. Together, these findings suggest that excitatory dorsal neurons form a laminar map which is imparted onto inhibitory dorsal neurons, sensory neurons, and likely also descending pathways from the brain.

Until now, late dorsal progenitors were thought to be a homogeneous group, raising questions about how the impressive diversity of neuron types is generated. We found a dorsal-to-ventral gradient of Zic transcription factors that patterned pdL progenitors, created an array of variants within dIL families, and was required to generate the normal complement of dorsal neurons. Zic genes had previously been described in the dorsal spinal cord, where Zic1 regulates early progenitor proliferation^55^ and Zic2 is known to mark mature spinal neurons that contribute to the post-synaptic dorsal column pathway^19^. It is not known how Zic proteins regulate dorsal neuron diversity, but an interesting hint may come from their similarity to Gli transcription factors^97,115,116^. As Gli factors mediate Hedgehog signaling to pattern the spinal cord ventral horn, Gli and Zic proteins may work together to establish countervailing molecular gradients.

Future work should probe the molecular mechanisms that transform the Zic gradient into gene expression signatures that characterize dorsal horn neurons.

Overall, our work supports a model in which four major principles govern dorsal horn ontogeny (Figure 6). First, six temporal cohorts of dIL neurons are born successively and give rise to the families of dorsal neuronal types. Second, each temporal cohort is patterned along a dorsal-ventral axis by Zic transcription factors, thereby fractionating each family into a range of variant types. Third, individual pdL progenitors can give rise to inhibitory dILA or excitatory dILB neurons as directed by Notch signaling, a process we did not study here but which has been shown previously^45,47^. Fourth, the excitatory neurons establish the anatomical structure of the dorsal horn by settling into an ordered distribution based on their birthdate.

**Figure 6:**
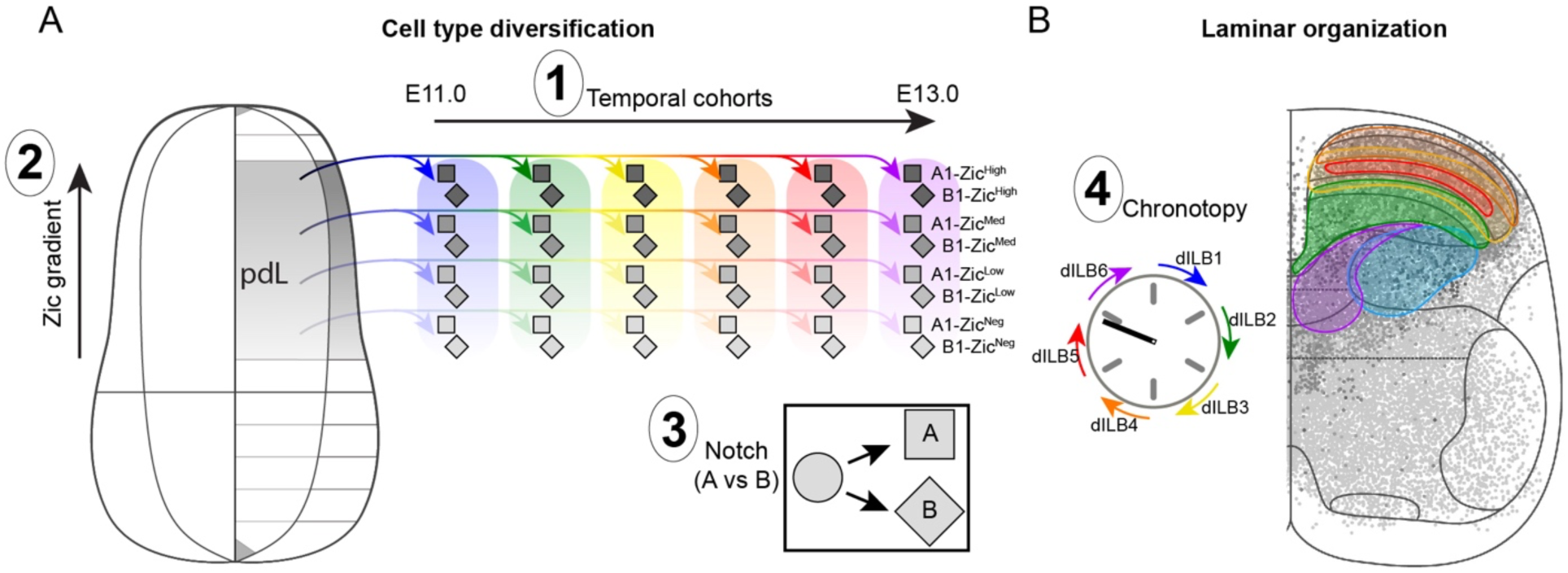
Four principles governing dorsal horn ontogeny. (A) Mechanisms of neuronal diversification: (1) Temporal cohorts of neurons are progressively born from the pdL progenitor zone, producing six cohorts of dILA and dILB neurons each; (2) dorsoventral position within the pdL progenitor zone produces variant neuron types particular to each dIL family; (3) previously known Notch-mediated mechanisms allow dILA and dILB neurons to be simultaneously born from asymmetric terminal progenitor divisions. (B) Mechanisms of laminar formation: (4) Sequentially born excitatory dILB families settle into spatially adjacent laminae.

As this picture comes into focus, the similarity to drosophila optic lobe development is striking: there is a similar number of discrete transcriptional cell types, similar temporal diversification which is transformed into a spatial arrangement for precise circuits, similar dorso-ventral spatial patterning that is most likely coordinated by the same signaling pathways (Shh/Hh, BMP/Dpp, and Wnt/Wg), similar binary cell fate decisions controlled by the Notch pathway, and even a possible similar role for Zic/Opa factors in regulating neuronal diversity^117–121^. These shared features in evolutionarily distant tissues and organisms may be coincidence, or they may reflect convergent forces that produce sensory systems specialized for multi-modal processing. In particular, the high degree of neuronal diversity (generated by the three-fold combination of temporal, spatial, and Notch mechanisms) and the layered anatomical microcircuit may facilitate the parsing, sharpening, blending, integration, and feature extraction from different neural information streams. The next frontier belongs to contextualizing this cellular diversity and structure in terms of behavior – that is, to reconcile these neuronal elements with their contributions to sensation, autonomic control, and movement.

## Resource availability

### Lead Contact

Data and resources will be shared by request by the lead contact.

### Materials availability

No unique materials were generated here. Sources of previously published reagents and mouse lines are noted in the Key Resources table.

### Data and code availability

Raw single-cell RNA-sequencing data (Fastq files), sample multiplexing information, and processed data (Cellranger count outputs and R objects processed with the Seurat package) will be made available on Gene Expression Omnibus (GEO) upon publication. Raw and processed spatial transcriptomics data from the 10X Xenium platform will be made available on GEO upon publication. All original analysis code will be made available for download at https://github.com/ArielLevineLabNINDS/Dorsal_Horn upon publication.

## Acknowledgements

This work was supported by NINDS Intramural funds through 1ZIA NS003153 (R.B.R, A.R., A.J.L.), NICHD R37HD091856 (L.F., J.E.J.), and NIH U54AG076040 (A.Y. and V.M.). We gratefully acknowledge the support, scientific feedback, and general discussion from Dena Goldblatt, Kaya Matson, Lily Li, Tim Petros, and Artur Kania, as well as the generous antibody gifts from Eloisa Herrera (Zic2), Thomas Muller (Lmx1b, Lbx1 and Foxd3), Michael Wegner (Sox5) and Jay Bikoff (Pou6f2).

## Author contributions

Conceptualization (R.B.R and A.J.L). Methodology (R.B.R.). Investigation (R.B.R., L.F., D.N., A.R., and A.J.L.). Formal analysis (R.B.R., A.Y.). Supervision (R.A., R.W., J.E.J., V.M., and A.J.L). Funding acquisition (J.E.J., R.W., V.M., and A.J.L). Visualization (R.B.R). Writing – original draft (R.B.R. and A.J.L.). All authors contributed to the review and editing of the manuscript.

## Supplemental information

Document S1. Figures S1-S19 and Tables S1, S3, S5 and S9.

Table S2. Markers of all spinal cells, related to Figure 1.

Table S4. Markers of neuron families, related to Figure 1.

Table S6. Refined neuron statistics, related to Figure 1.

Table S7. Confusion matrix for refined neuron types, related to Figure 1.

Table S8. Markers of refined dIL neuron types, related to Figure 1.

Table S10. Top PC loadings of dIL families, related to Figure 5.

## STAR Methods

### Experimental model and subject details

#### Animal husbandry and ethical approvals

The following mouse strains were maintained at the National Institutes of Health in Bethesda, MD, and all experiments received assent from the National Institute of Neurological Disorders and Stroke Animal Care and Use Committee (ACUC protocol number 1384). All wild-type mice used in this study were a mix of BALB/cJ and C57BL/6J strains. *Pax3^Cre^*, *Isl2^DTA^,* and *Zic1/Zic4^+/-^* mice were obtained from The Jackson Laboratory (Bar Harbor, ME, USA).

The *Gsx1^+/-^* and *Gsx2^+/-^* mouse strains were maintained at Cincinnati Children’s Hospital Medical Center, and experiments involving these strains were reviewed and approved by the Cincinnati Children’s Hospital Medical Center Institutional Animal Care and Use Committee.

The *Zic3^Ka^* mouse strain was maintained at Australian National University. Mice were maintained according to Australian Standards for Animal Care under protocol A2021/39 approved by The Australian National University Animal Ethics and Experimentation Committee. The katun (*Ka*) allele (MGI: 3043027) of *Zic3* (*Zic3^Ka^*)^103^ was maintained by continuous backcross to C57BL/6J inbred mice; animals from backcross 10 and beyond were used for analysis.

The *Ptf1a^Cre^* mouse line was maintained at the University of Texas Southwestern. All mouse protocols were approved by the Institutional Animal Care and Use Committee at the University of Texas Southwestern Medical Center. All mouse strains were maintained on a mixed background of ICR and C57BL/6.

Only *Zic3^Ka^* embryos were sex-typed (due to *Zic3* being on the X-chromosome), where even numbers of male and female wild-type and null embryos were examined. Embryos of all other experiments were not sex-typed and experiments were assumed to contain an even male:female ratio. Mice housed at all facilities were given ad-libitum access to food and water, maintained on a 12-hour light:dark cycle (6:00-18:00 light period; following daylight savings time).

### Embryonic spinal cord tissue

Appropriate pairs of male and female mice were introduced to the same home cage at approximately 18:00, and checked daily thereafter each morning at approximately 10:00 for vaginal plugs. 12:00 on the day a plug was detected was considered (embryonic day) E0.5, and embryos were dissected at approximately noon on the embryonic day indicated by the experiments described here (below) and in Results. Particular care was taken to ensure EdU birthdating was as accurate as possible; for these experiments, males and females were paired only one night per week, and separated in the morning regardless of whether a plug was detected. Only females which plugged received a subsequent i.p injection of EdU (more details below), ensuring that time of conception was constrained to approximately the dark period (i.e. with an uncertainty of plus or minus 8 hours).

On the indicated embryonic day, female mice were administered CO2 euthanasia, and embryos were promptly dissected and washed in ice-cold 1X PBS (except in cold HBSS for preparation for single-cell suspensions in the 10X Chromium assay). The spinal column was severed at the spinal cord-hindbrain junction, and the column was further dissected from the torso. For experiments prepared at the National Institutes of Health, the vertebral body was further dissected to expose the ventral surface of the spinal cord to fixative; for experiments performed elsewhere, dissection only involved isolation of the column with contained spinal cord. Spinal columns were immersion-fixed in ice-cold 4% paraformaldehyde for either 4-6 hours for immunohistochemical applications, or overnight (20-24 hours) for in-situ hybridization applications (both 10X Xenium and RNA Scope). After fixation, tissue samples were briefly rinsed in 1X PBS (<10 minutes) and then equilibrated to a solution of 30% sucrose plus 1X PBS overnight or until tissue sunk to the bottom of the vessel containing the tissue. In all cases tissues were agitated on an orbital or nutating shaker (depending on the vessel type) during the fixation and cryoprotection intervals in order to facilitate equilibration of tissues to the PFA or sucrose solutions.

### Genotyping

Primer sets are detailed in the Key Resources table. Genotyping was done as described by original publications for each allele (noted in Key Resources table). Expected band sizes are: Ptf1a^Cre^ (Wild-type: 123 bp, Cre: 300 bp), Zic1/4^DKO^ (Wild-type: 465 bp, Mutant: 400 bp), Gsx1^+/-^ (Wild-type: 219 bp, Mutant: 180 bp), Gsx2^Ra^ (otherwise referred to as Gsx2^+/-^, Wild-type: 298 bp, Mutant: 130 bp). Pax3^Cre^ and Isl2^DTA^ embryos were genotyped by Transnetyx (Cordova, TN, USA) using custom primer sets for the alleles (see Jackson Laboratory pages for original primer sets). For purposes of colony maintenance, Pax3^Cre^ mice were genotyped by the presence of a white splotch on their belly fur at the time of weaning (typical of mice bearing one null Pax3 allele). *Zic3^Ka^* embryos were genotyped as sex-typed as follows: Genomic DNA was prepared for genotyping as previously described^104^; 50 ng of genomic DNA was added to each genotyping reaction. The *Zic3^Ka^* strain was genotyped via Allelic Discrimination using the oligonucleotides Ark241_Katun_F and Ark242_Katun_R and probes Wildtype: 5’ 6FAM and G>T mutation: 5’ VIC, with TaqMan® Universal PCR Master Mix (Life Technologies; Cat. No. 4304437). The products were amplified and analysed using a StepOnePlus™ Real-Time PCR System (Applied Biosystems®) and StepOne software (version 2.2.2; Applied Biosystems®). The embryos were sex-typed using MyTaq HS Red (Bioline: BIO-25048) and touchdown PCR with an initial annealing temperature of 60°C, as previously described^104^. The sex was determined using the oligonucleotides Ark1004_Smcx_y_F and Ark1005_Smcx_y_R. Amplicons were separated via agarose gel electrophoresis (2% agarose in 1X tris-borate-EDTA buffer).

## Method Details

### Birthdating of spinal cord tissue

Pregnant female mice (detailed above) were injected with an intra-peritoneal solution containing 50 mg/kg of 5-Ethynyl-2ʹ-deoxyuridine (EdU, Thermo Fisher Scientific) at a concentration of 2 mg/ml in sterile 1x PBS. Injections were performed on separate pregnant females at either 12:00 (for the midpoint of an embryonic day - i.e. E12.5) or 0:00 (for the beginning of an embryonic day - i.e. E12.0). Only one injection at only one particular time point was performed for each individual female. Two separate cohorts of embryos were produced for the present timecourse – an initial cohort which were briefly fixed (4-6 h) suitable for immunohistochemistry (examined in figures 1 and 3), and a second cohort where equal numbers of embryos from each litter were fixed for either 4-6 h or for 20-24 h (more suitable for in-situ hybridization) and which were examined in figure 2. A cohort of embryos with IHC and ISH-suitable fix conditions were necessary to make within-litter comparisons of EdU co-labeling between all dIL families, some of which are best labeled with either IHC or ISH. Histochemistry to detect the presence of EdU in spinal cord nuclei is detailed below.

### Cryosectioning

All tissues were frozen while immersed in OCT compound in a cubic mould. Samples from littermate embryos were always embedded and frozen in the same block of OCT. Moulds were immersed in 100% ethanol chilled with dry ice up to the top of the mould, though direct contact of OCT compound and ethanol was avoided. Blocks were maintained at -80 °C after freezing until cryosectioning, at which points blocks were equilibrated to the cryostat temperature (generally -18 °C to -24 °C) and cut in 25 μm sections (except for 10X Xenium, which were cut at 10 μm). Sections were mounted on microscope slides which were then stored at -80 °C prior to histochemistry.

### Immunohistochemistry (IHC)

All IHC was performed using published primary antibodies (detailed in Key Resources table). A wide variety of secondary antibodies were used here and were Donkey IgG Alexa-fluor conjugates purchased from either Invitrogen or (for Donkey anti- Guinea pig secondary antibodies) Jackson Immunolabs. Working dilutions of antibodies from their manufactured concentrations are detailed in the Key Resources table. Briefly, slides were removed from -80 °C and dried at 37 °C for at least 20 minutes. Sections were then surrounded by a hydrophobic barrier (Vector Laboratories) and dried at room temperature another 5 minutes. Sections were then washed three times successively for 10 minutes each using cold 1x PBS at 4 °C. Working dilutions of primary antibodies were prepared in 1x PBS with 0.1% Triton X- 100, applied inside the hydrophobic barrier, and incubated overnight at 4 °C. Subsequently slides were again washed three times successively for 10 minutes each using cold 1x PBS at 4 °C. Working dilutions of appropriate Alexa-Fluor-conjugated secondary antibodies together with DAPI (for nuclear staining) were prepared in 1x PBS with 0.1% Triton X-100, applied inside the hydrophobic barrier, and incubated for 1 hour at 4 °C. Finally, slides were once again washed three times successively for 10 minutes each using cold 1x PBS at 4 °C. After the final 1x PBS wash was removed from the slide, 2-4 drops of Prolong-Gold (Thermo Fisher Scientific) was applied to the slide and the coverslip was gently laid to spread the medium and avoid trapping bubbles. Covered slides were allowed to dry at least 1 hour prior to imaging and stored at 4 °C prior to and after imaging.

### In-situ hybridization (ISH)

All ISH was performed based on the published protocols for ACD Bio RNA Scope Multiplex Fluoresence (version 2) assay. All reagents and probes were manufactured by ACD Bio and are listed in the Key Resources table. Target retrieval was omitted from the tissue pretreatment protocol for all tissues examined here, due to the step producing excessive damage to embryonic sections. Probe fluorescence signal was not evidently changed in embryonic spinal cord sections receiving versus not receiving target retrieval. Briefly, sections were dried for two hours at room temperature prior to the modified tissue pretreatment protocol. Hybridization, development of fluorescence signal, nuclear staining with DAPI, and cover-slip application was done exactly as described by protocol. Covered slides were allowed to dry at least 1 hour prior to imaging and stored at 4 °C prior to and after imaging. Fluorescence was developed using the newer TSA-Vivid dyes (Vivid 520, Vivid 570 and Vivid 650).

### Pixel-frequency depth histograms

For selected IHC and ISH signals, histograms of pixel- frequency were gathered from E16.5 wild-type, *Ptf1a*-null, *Gsx1/2*-null and sensory neuron-ablated spinal cords. This was done using the Plot Profile function of ImageJ/Fiji on 8-bit grey thresholded images. To measure pixels where two markers colocalize images of both markers were separately thresholded and pixels were multiplied using ImageJ/Fiji’s image calculator, with the histogram being applied to the product image. One to three sections per biological replicate were analyzed and, whenever possible, each hemisection was measured separately. The histogram was collected vertically, with one end of the range at the pial surface of the superficial dorsal horn midway through its mediolateral extent, and the other end of the range at the dorsal-ventral border of the ventricular midline (where both sides become adhered). The histogram width (i.e. how much sampling was done) varied, and generally encompassed one third of the total dorsal horn width except smaller when avoiding sampling autofluorescent vasculature. Histogram bins were compressed to a common number (80) in order to allow comparisons between genotypes, and bin frequencies were normalized to the total counts for that particular bin per replicate. Lamina VI was mostly excluded from this analysis due to the lack of dILB neurons there; accordingly, the dorsal-most 56 bins were plotted in figures, with color saturation corresponding to bin weights.

### EdU histochemistry reaction

In the event of EdU histochemistry, slides being reacted for IHC or ISH were not covered at the end of the protocol; instead, they were washed in 1x PBS with 0.1% Triton X-100 for 30 minutes at 4 °C. Following this, they were incubated at room temperature in a solution of EdU reaction buffer, buffer additive, copper (II) sulfate, and Alexa-Fluor-conjugated azide (Alexa-Fluor-488 used in conjunction with Vivid-570 and Vivid-650 dyes on RNA Scope- reacted slides, or Alexa-Fluor-647 with IHC-reacted slides having omitting Alexa-Fluor-647- conjugated secondary antibodies) at the ratios indicated for the Click-iT EdU Cell Proliferation Kit (Thermo Fisher Scientific). They were subsequently washed three times successively for 10 minutes each using cold 1x PBS at 4 °C, cover-slipped and stored per IHC protocols above.

### Microscopy

All micrographs presented here were obtained either from inverted confocal microscopes (Zeiss LSM-800), or from a Zeiss AxioImager Z2 equipped with Apotome 3 structured illumination and DAPI, GFP, DsRed, Cy5, and Cy7 filter cubes using Zeiss ZEN software. In either case, micrographs were composed of a tile-scan of spinal cord sections using a 20x objective as well as a z-stack of either 6-9 intervals centered approximately mid-way through the tissue section’s thickness. For experiments using Apotome, five images were taken per field of view, each using different positions of the structured illumination grid, and were deconvoluted using ZEN’s Apotome RAW Convert function. Image arrays from both Apotome and confocal experiments were stitched using Zeiss ZEN’s stitching function; all channels at all z- positions were stitched with reference to nuclear DAPI signal from images in the mid-point of the stack. Images were stored in Zeiss’ .czi format, and were subsequently loaded into Fiji (Image J, v2.3.0) with the current Bio-Formats plugin.

### EdU counts

All birthdating data in this study was done on cervical spinal cord sections, between C3-C8. Images with EdU and signal from two other markers (either from IHC or ISH) were counted in ImageJ using the Cell Counter plugin. Cells were considered positive for EdU if the nucleus was mostly filled with signal and excluded if the nucleus was a discontinuous set of puncta. Overlap with either transcription factor IHC signal or with RNA Scope was considered as a positive colocalized cell and was counted separately to either cells only labeled with EdU or cells only labeled with cell-type markers respectively. Anchor points were labeled on each section in order to allow coordinate normalization in data processing. Briefly, a set of ten points were added to each section: two on either dorsal horn’s most dorsal extent; two on either dorsal horn’s lateral-most extent, one at the base of the dorsal columns, two on either ventral horn’s lateral-most extent, two on either ventral horn’s most ventral extent, and one on a point along the midline where ependymal/progenitor cells diverge dorsally but are adhered ventrally; this point is considered to be the boundary between dorsal and ventral horns. Coordinates are then exported using the Measure tool.

### EdU counts - cumulative counts statistics

At least one embryo was assessed per litter; for each embryo, the fraction of co-labeled neurons as a fraction of all neurons of any dIL family at any particular timepoint of EdU pulse were represented as a fraction. The mean of these fractions across all embryos of a litter was counted as a single data point, as we judged that variance across litters was higher than within litters, hence the insufficiency of only one litter for this analysis (regardless of whether that litter had embryos in triplicate).

To produce cumulative counts for each timepoint, the mean co-labeling of each dIL family for each timepoint was produced. This value was then normalized to the fraction of mean co-labeling across all timepoints, so that it reflected the percent of dIL family neurons born at a certain time point. From here, mean co-labeling for all prior timepoints were added to the replicate data of each timepoint such that within timepoint variation would be preserved but that mean cumulative co-labeling at the final timepoint would be 100%. Data were plotted as mean ± standard deviation and sigmoidal fit curves were calculated and plotted in Graphpad Prism using “XY analyses” -> “Interpolate a standard curve” -> “Sigmoidal, 4PL, X is concentration”. Sigmoidal fit could not be done for two timepoints (dILA2 and dILB1) where substantial neurogenesis between E10.5 and E11.0 produced a negligible slow-growth phase to the curve; these curves were instead calculated according to the “Asymmetric Sigmoidal, 5PL, X is concentration” fit. EC50/IC50 values reported by the standard curve interpolation were plotted as vertical lines on the time (x-) axis of standard curve plots in colors corresponding to each of six dILA or dILB families respectively.

### Cell coordinate rotation and normalization

For EdU cell count coordinates (and later also for spatial transcriptomics cell coordinates), cell coordinates were normalized so that statistics and graphics could be produced systematically on cell coordinates across sections and experimental groups. To do this, anchor points (described in **EdU counts** above) were used to transform all coordinates identified from a section. Two midline coordinate pairs (from the dorsal columns base and at the dorsalhorn-ventral-horn junction) were used to rotate all coordinates such that both coordinate pairs had identical x-coordinates. Each hemisection was then stretched to fit an ideal diagram of a spinal cord; y-coordinates from the coordinate pairs at the dorsal surface of the dorsal horn and ventral surface of the ventral horn were normalized to either +2000 or - 2000 arbitrary units of distance. Likewise, the x-coordinates of coordinate pairs at the lateral-most extent of each dorsal horn were normalized to either –2000 or +2000. X and Y coordinates of all intervening spinal cord neuron data were linearly transformed based on their distance from these anchor points. Additionally, neurons were linearly transformed based on their proximity to the y-coordinates of the lateral dorsal horn coordinate pairs or to the y-coordinate of the anchor at the base of the dorsal columns. The normalized value for these were also arbitrary, and changed based on segmental group to best fit an ideal spinal cord diagram; because of this, statistical comparisons between data from different spinal cord segments were not done – only within segment and between EdU pulse time (or genotype). Cell coordinates from all conditions were collated in a single database and analyzed further.

### Contour plots

Due to variability in time of conception, a certain amount of overlap in terms of the patterns of EdU-dIL family co-labeling was expected and observed for adjacent families, and between group differences are difficult to judge on a scatter plot. We therefore prepared and plotted 2-dimensional kernel density estimates for EdU birthdating cell coordinate data. This was done using the kde2d function from the R MASS library. Kernel density estimates were produced at a resolution of 100 for cell coordinates from all litter replicates and all EdU timepoints, which were then collated in a database. Contours were plotted in ggplot2 using geom_contour_filled and drawn to fit the densest 50%, 25%, 15%, 5% and 2.5% of points.

Broader contour levels were drawn with greater transparency to represent lower cell density in these levels.

### Cell preparation for single-cell RNA-sequencing

Female wild-type mice with a pregnancy dated to embryonic day E14.5 or E16.5 were used. Each experimental replicate consisted of a pooled litter of 3-10 spinal cords taken from embryos of one single pregnant female.

Embryos were rapidly dissected and immersed in cold 1X PBS. Spinal cords were sequentially dissected as described above and placed in cold dissection medium (50 ml HBSS- calcium/magnesium-free with phenol red (Gibco) + 750 µl HEPES (1M, Gibco)). When possible, extra care was taken to remove meninges and dorsal root ganglia from the isolated cord. From here, appropriate spinal cord segments were dissected from the cord, and then cut into fine pieces with dissecting spring scissors. Either cervical, thoracic or lumbosacral cords were isolated based on the experimental group. In either case, care was taken not to include transitional zones of the cord (e.g. where cervical tapers into thoracic or where thoracic begins to enlarge to lumbar); due to this, these regions are likely underrepresented in our data.

Pooled spinal cord tissues were then added to a 15 ml conical tube containing dissociation medium (100 units Papain (Worthington), 4.75 ml HBSS+CM (calcium and magnesium; Gibco), and 250 µl of DNAse (Worthington, 1000 units resuspended in 500 µl of HBSS+CM)). The tube was gently inverted several times every 5 minutes while leaving the tube to incubate at 37 °C between inversions. After 20 minutes, the suspension was triturated gently with a series of three fire-polished glass pipettes (Fisher Scientific) with progressively finer bores. Suspensions were then centrifuged for 8 minutes at 850 rpm; after this, the supernatant was removed and the cells were gently resuspended with a solution of 4.5 ml HBSS+CM, 600 µl ovomucoid protease inhibitor solution (Worthington, resuspended in 30 ml HBSS+CM), and 250 µl DNAse (as above). Debris was allowed to settle for 10 minutes, at which point the supernatant with suspended cells was added to a second 15 ml conical tube containing neutralizing solution (10 ml HBSS+CM + 100 mg Trypsin inhibitor (Sigma) + 100 mg bovine serum albumin (Sigma)). This suspension was centrifuged once more for 8 minutes at 850 rpm, and the supernatant was removed. The remaining cells were resuspended in approximately 1 ml of 1X PBS containing 5 µl of molecular-grade BSA (New England Biolabs) and 2 µl of RNAse-inhibitor (Biosearch Technologies). Cell concentration in this suspension was assessed using a standard hemacytometer (a 50:50 sample of cell suspension + Trypan blue). The cell suspension was diluted with the same resuspension medium until the final concentration was appropriate for loading into the 10X Genomics Chromium chip.

### Single-cell RNA-sequencing - data collection

Cell suspensions were introduced to 10X Genomics Chromium chips at concentrations appropriate for retrieval of data from an estimated 7,000 cells. Two Chromium wells were loaded per pooled biological replicate, with an anticipated 14,000 cells retrieved per replicate. cDNA and libraries were constructed based on the 10X Genomics Chromium 3’ gene-expression chemistry (v3.1). Library fragment size and variation were inspected on an Agilent 2100 bioanalyzer, then pooled for sequencing. Sequencing was done using the Novaseq 6000 platform, with the aim of sequencing 40,000 reads per cell. Published datasets were retrieved from GEO or ArrayExpress, per instructions from original publications^50,52,72^.

### Single-cell RNA-sequencing - data processing

Raw sequencing data was processed first with Cellranger mkfastq, and those outputs were then run in Cellranger count to produce count matrices. Reads were mapped onto the *mus musculus* GRCm38-mm10 genome reference. Post-hoc quality control per sample (including effective number of cells retrieved, number of reads and genes detected per cell, etc) are detailed in Table S1. Sample 16B (E16.5 cervical spinal cord replicate 1 reaction 2) was excluded from further analyses at this stage based on an observably poor emulsion post Chromium run, poor read quality and almost negligible cell retrieval.

Further quality control, clustering and annotation were done using Seurat v4 (though Seurat v5 was released in the interim) in R (v4.4.1). To begin in Seurat, data from each sample was imported into Seurat and minimum thresholds on count and gene numbers per cell, as well as maximum thresholds on mitochondrial gene percentages per cell were imposed (Table S1).

From this, the data were log-normalized, variable features were found, and CCA integration (Seurat v4) was run using a list of biological replicates as inputs. At this stage, 195,146 cells were retained.

Principal component (PC) analysis (PCA) was run on the integrated dataset of all biological replicates using 90 PCs, while also regressing gene counts per cell. After clustering all cells at resolution 2, clusters were annotated as major cell-types of the spinal column based on top marker genes per cluster (Table S2). Spinal cord neurons were extracted from this broader dataset of sensory neurons and non-neuronal cells, and this subset was further processed.

Principal component analysis of spinal cord neurons was once again done using 90 PCs, and the dataset was subsequently re-clustered, excluding clusters whose principal marker gene sets were non-neuronal, whose principal marker genes were a combination of neuronal and non- neuronal in addition to substantially higher gene counts (assumed doublets), and/or whose features were of a variety of cell-types in addition to substantially lower gene counts (assumed unhealthy cells and/or data produced from poor emulsion quality).

Clustering of spinal cord neurons was done using 90 PCs at resolution 4. Finally, we split the dataset into three major groups: Group 1 – the well-described cardinal neuron classes as Group 1 (dI1-dI6, v0-v3 and motor neurons); Group 2 – the presumptive dILA neurons; and Group 3 – the presumptive dILB neurons. As the dimensionality of this dataset was evidently quite high, dILA and dILB neurons were clustered together using 150 PCs and annotated by family at resolution 32, while cardinal neuron classes were clustered separately using 180 PCs and annotated by class at resolution 32. Neurons were annotated by cluster and also by major classes (families of dIL neurons or cardinal classes) and these annotations were re-introduced to the dataset comprising all spinal cord neurons. This version of the dataset was presented in main figures, and used as a reference to annotate spatial transcriptomics data as well as to annotate single-nucleus RNA-sequencing data retrieved from Qiu and colleagues (described below)^72^. At this final stage of processing, 94,048 spinal cord neurons were retained.

PHATE trajectory inference was performed using PHATE^71^ (R library “phateR”), with the following parameters: knn = 20, gamma = 0, t = 100, and 90 PCs for dimensionality reduction and visualization. Cluster robustness was performed for all neurons at the level of family, as well as the refined types of dILA and dILB using Silhouette and LISI scores (R libraries “lisi” and “cluster”) and computed with 90 principal components (PCs) each. Dendrograms for different neuronal subsets were generated based on centroid distances. The primary dendrogram, including all refined neuron types, was constructed using 150 PCs (Figure 1F). A second, family-level dendrogram—excluding "Hindbrain"-like, "Immature", and "Other" neuron types—was generated using 90 PCs (Figure S3). Additionally, dendrograms for the dILA and dILA separately (Figure S3) were constructed using 90 PCs.

### Single-cell RNA-sequencing - annotation transfer to published datasets

Data from Qiu and colleagues were retrieved in order to examine how dIL families arise at precisely defined embryonic stages. We first extracted all CNS neurons from timepoints E11.0 to E16.75 and, using iterative clustering and filtering, excluded all clusters which were evidently not spinal neurons – exclusion criteria included presence of Hox genes of group 1-3 but not group 4 and beyond, Lhx6/Lhx8 (presumptive forebrain inhibitory), Tfap2d (presumptive midbrain) and excitatory En1+ cells (presumptive midbrain). After first-pass filtering, data from all timepoints between E11.0 to E16.75 were integrated together using Seurat v5 integration, reclustered using 120 PCs, and cell-type identity was predicted by transferring annotations from single-cell RNA-sequencing data generated here (see Single-cell RNA-sequencing - data processing and Fig. 1) using 120 PCs. As post-integration clustering allowed clear separation of hindbrain and non-hindbrain cardinal classes by absence or presence of Hox group 4 genes and beyond, hindbrain clusters were further filtered using these criteria. From this point, a contingency table of predicted cell types was generated for each individual time point. Data were presented as the fraction of all dIL families and immature neurons present at each specific timepoint (done separately for dILA and dILB neurons).

### Single-cell RNA-sequencing – examination of published data

To examine onset of Zic gene expression in dIL neurons, single-cell RNA-sequencing data from Delile et al. (2019) consisting of only spinal cord neurons at timepoints E9.5, E10.5, E11.5, E12.5, and E13.5, as well as spinal cord neurons from Osseward and colleagues at E12.5 were integrated together (six input datasets. Cell numbers: (N=356 E9.5, N=1724 E10.5, N=3588 E11.5, N=4561 E12.5, N=5542 E13.5)^52^; (N=2914 E12.5).) using Seurat v4 integration, and clustered using 60 PCs at resolution 32. Many annotations from Delile et al. (2019) (e.g. dI5.3, dI5.1) corresponded to immature dIL neurons which bore no markers corresponding to mature neuron cognates in single-cell/nucleus RNA-sequencing data generated here or from adult datasets. Accordingly, we annotated clusters based on markers present in dIL neurons which no longer bore immature markers (i.e. Robo3, Nhlh1/2, Klhl35); from data prior to and including E13.5, only dILB1 (Plscr5/Skor1/Tac1), dILB2 (Maf/Cbln1), dILA1 (Stc1/Plch1) and dILA2 (Rorb/Sall3) could be reliably identified; dI5 was inferred to be clusters bearing Phox2a/Lmx1b, and dI4 was inferred to be early-born (Zfhx3- high+) Lbx1+ inhibitory neurons clustering separately from those expressing Dmrt3 (dI6). To examine ranked expression of *Zic1* among pdL progenitors, neural progenitor-annotated data was retrieved from Delile et al. (2019) from timepoints E11.5 and E12.5 (both timepoints of which are confidently during dIL neurogenesis and beyond dI4/dI5 neurogenesis); to enrich pdL progenitors, ventral progenitors (*Dbx2*, *Nkx6-1*, *Nkx2-2*, and *Olig2*) and dI1-3 progenitors (*Olig3*) were excluded by their respective marker sets. Remaining pdL progenitors expressed *Gsx1*, *Gsx2* and *Ascl1*, as expected.

### Spatial transcriptomics - data collection

All tissues used for spatial transcriptomics were immersion fixed overnight during (20-24 h), identically to those used for ISH (see **Generation of embryonic spinal cord tissue**) and generally using adjacent sections from the same embryos examined in ISH. To determine spatial arrangement of dorsal horn neuron subtypes in normal and mutant spinal cords we used the 10X Genomics Xenium (v1) platform. As much of the mutant tissue analyzed here was fixed-frozen, and as Xenium v1 did not yet have a supported fixed frozen tissue protocol, we followed the guide for fresh-frozen tissues with modifications detailed below. Cryosectioning was done as described in CG000579 revision C (10X Genomics) and fixation and permeabilization was done as described in CG000581 revision C (10X Genomics), except that the 30-minute incubation in fixation solution was omitted. Tissue chemistry was done as described in CG000582 revision F (10X Genomics). Probe-sets used were the v1 mouse brain panel (248 genes) plus 100 custom probes (Table S8) marking either classes or individual clusters from the single-cell RNA-sequencing dataset collected here. A reference dataset was produced using four wildtype E16.5 spinal cords spanning upper cervical to sacral segments, to most accurately capture the variation of the dorsal horn across segments and animals, and these comprised the first of two slides. The second slide sought to capture a variety of mutant tissues. Operation of the Xenium Analyzer instrument was done as described in CG000584 revision E (10X Genomics). Sections were grouped by animal or genotype during Xenium analyzer setup, and these data are also noted in the table. Areas which contained overlapping sections were omitted from this step and were not further analyzed.

### Spatial transcriptomics - data processing

Raw 10X Xenium datasets obtained from the Xenium analyzer computer were imported into Seurat, cells with 0 transcripts were removed, all cells in a group were clustered using 60 PCs at 0.5 resolution, and Seurat objects were written as .RDS files (see code **Xenium_2024_1_FilterClusterExport** on Github). Subsequently, coordinates were assigned bounding each section, clusters and gene markers were displayed on a UMAP reduction, and non-neuronal and sensory neuron clusters were excluded from further processing (see code **Xenium_2024_2_DefineROI_ExtractNeurons** on Github). From filtered spinal neurons, annotations were predicted using Robust Cell-Type Decomposition (RCTD)^50,78^ from the R package spacexr and our single-cell RNA-sequencing annotations (described above) as reference. Based on predicted cell-types, family-level metadata was assigned to each cell.

Further, a section identifier, and an estimated spinal segment were added as metadata. Additional exclusion of non-neuronal clusters was done here, particularly for clusters which had widely indistinct or sparse annotation transfer; on inspection, these were often clusters with most cells bearing both neuronal and non-neuronal markers resulting from poor cell segmentation or minor cell overlay. Cell-type proportions were also examined here separately per spinal cord zone (C1-C4 – Cervical, C5-C8 – Brachial, Thoracic, L1-L3 – Upper lumbar, L4-L6 – Lower lumbar or Sacral); to ensure proportions were not affected by exclusion of particular parts of a section only complete sections or complete hemisections were included; from these, cell proportions are represented as a fraction of all neurons from the total sections or hemisections examined of a single embryo. Finally, cell coordinates and appropriate metadata were exported as .csv files for localization analysis (see code **Xenium_2024_3_RCTD** on Github for above processing steps). Subsequently, anatomical landmarks to be used as anchors for coordinate normalization were identified for each individual section retained as this stage (see **EdU counts - coordinate rotation and normalization** for details). Coordinates from all sections of a given genotype and/or spinal segment were combined and overlaid on each other to best visualize neuron-type location across sections. These coordinates were subsequently exported as scatter plots for figures (see code **Xenium_2024_4_Coordinate_processing_overlay** on Github for above processing steps). To illustrate the relative dorso-ventral frequency of neurons from each dIL family (Figure 3C), the normalized y-coordinate of all dIL neurons was assigned to one of 200 bins spanning the full extent of both dorsal and ventral horns. Histograms were smoothed (using Ggplot2 stat_smooth) and plotted as frequency (x-axis) and dorsoventral bin order (y-axis).

Data are made available as raw Xenium outputs, Seurat objects containing all section ROIs filtered to remove non-neurons, normalized collated cell coordinates from all sections within the indicated segmental zone, and (for selected analyses) mirrored normalized collated cell coordinates from both wild-type (coordinates x<0) and null (coordinates x>0) embryos with wild-type cells downsampled to approximately match the number of null cells (numbers of each predicted cell-type were downsampled based on the ratio of total null to total wild-type neurons).

## Spatial transcriptomics – statistics

To understand changes in cell-type frequency between genotypes, contigency tables of predicted cell-types from E16.5 wild-type, *Zic1/4*-null and *Zic3*-null spinal cords were generated separately for each biological replicate and then normalized to the number sections collected per replicate. Selected cell-type frequencies were compared across all three groups in a two-way ANOVA (cell-type x genotype), and the post-hoc Tukey’s multiple comparisons test was done between all genotypes for each cell-type.

**Supplemental Tables:**

