## Supplemental materials for "Ontogeny of the spinal cord dorsal horn"

**Supplemental material:**

Figure S1: Generation of embryonic spinal cord single-cell RNA-sequencing data

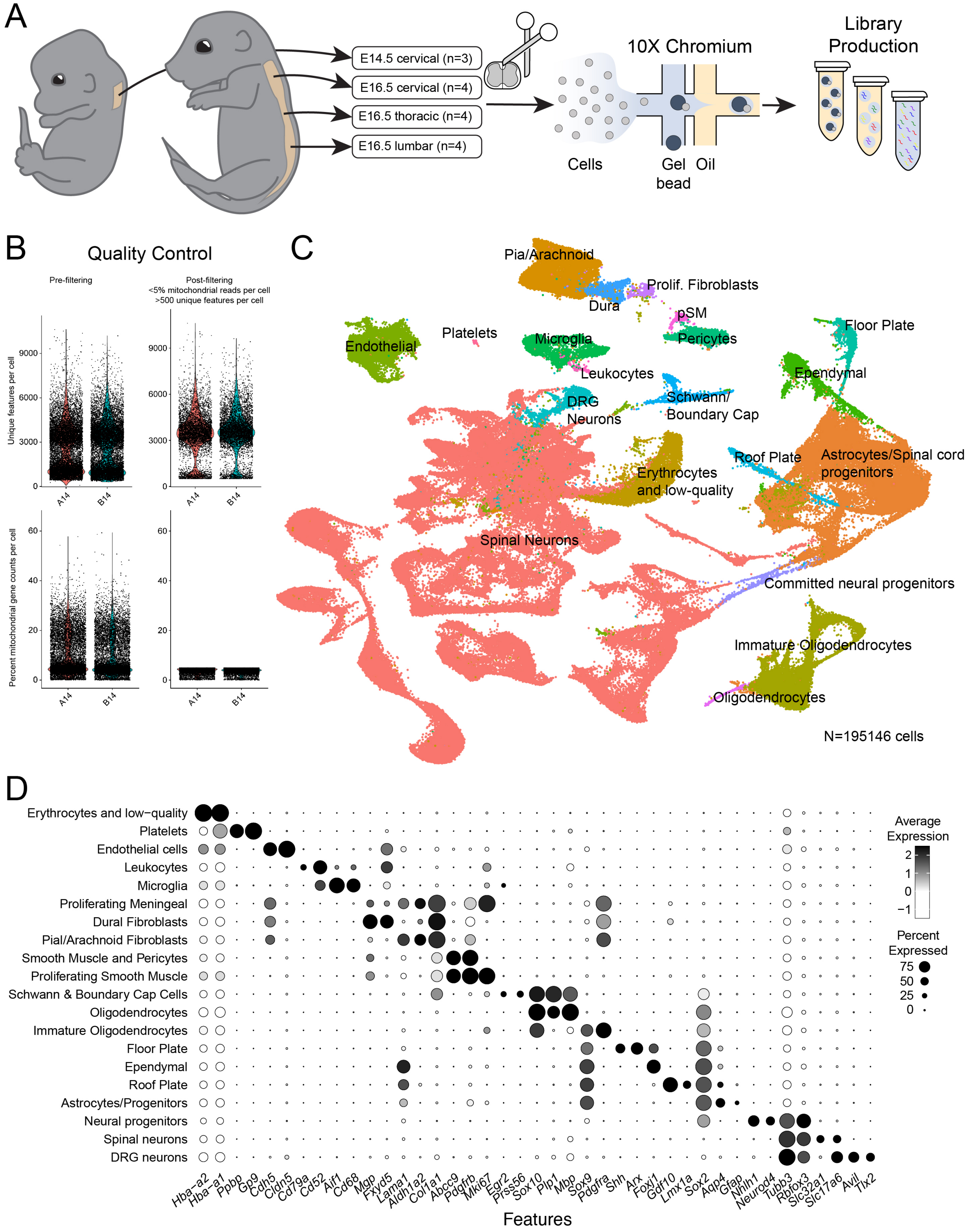

**Figure S1: Generation of embryonic spinal cord single-cell RNA-sequencing data, related to Figure 1.**

(A) Spinal cords were extracted from E14.5 and E16.5 spinal cords, dissociated, run through the 10X Chromium 3’ gene expression pipeline, and libraries were produced and sequenced. (B) After importation into Seurat, poor quality cells with low unique features or a high percentage of mitochondrial gene reads were removed from the dataset. (C) UMAP plot of the high-quality cells, following integration and clustering. (D) Dot plot showing all major annotated spinal cord cell types, showing marker gene sets used to classify clusters into archetypal spinal cord cell types. Dot size represents the fraction of a cell type expressing a gene, and dot shading represents the average normalized expression of a gene per cell type.
Numbers: Pooled embryonic spinal cord replicates (A): E14.5 cervical spinal cord (n=3), E16.5 cervical spinal cord (n=4), E16.5 thoracic spinal cord (n=4), E16.5 lumbosacral spinal cord (n=4). Total cells after quality control: N=195146.
Abbreviations: DEG – differentially-expressed gene, UMAP – uniform manifold approximation and projection.

Figure S2: Zic1 expression defines the boundary of dI4/dILA and dI5/dILB

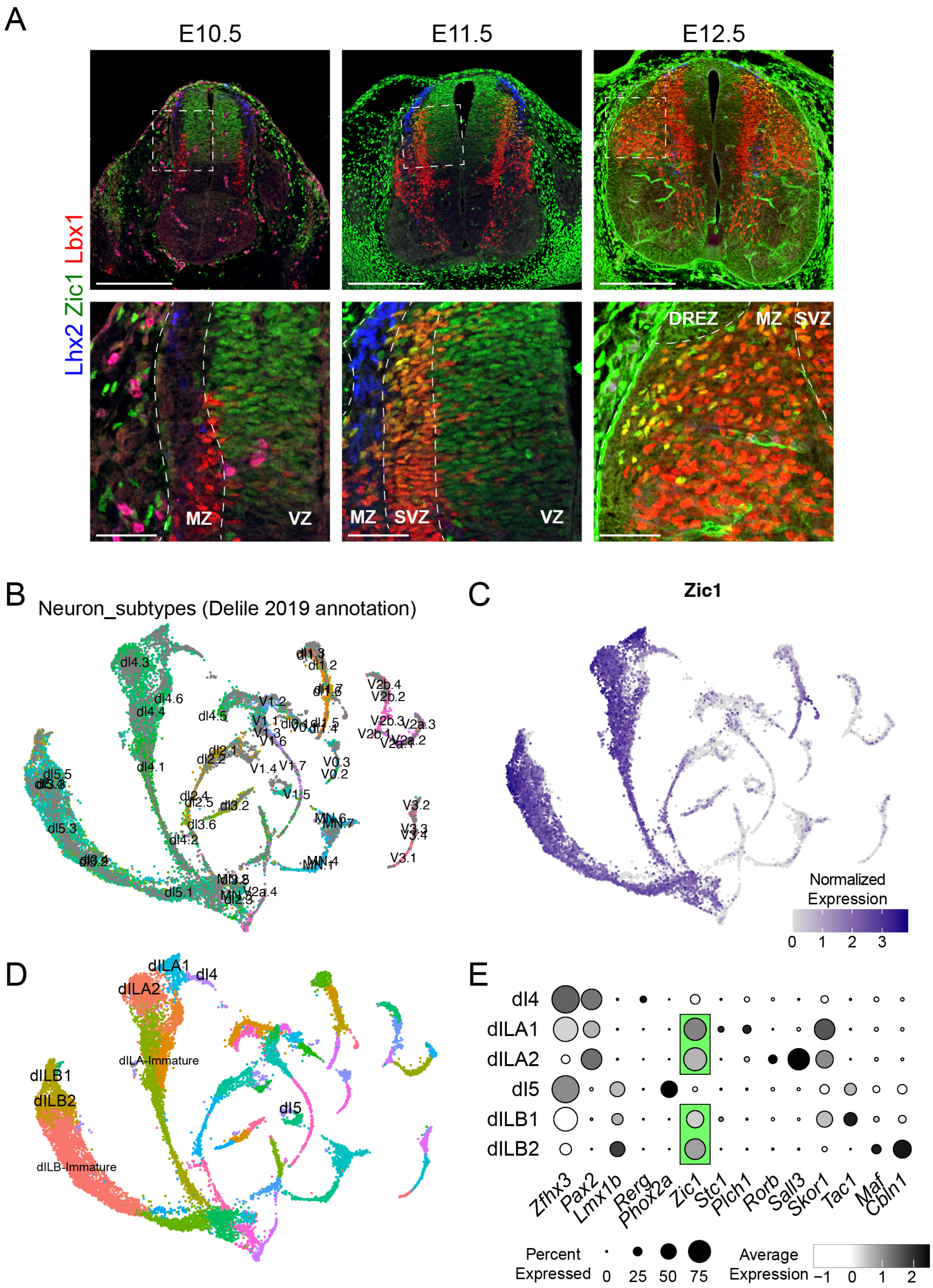

**Figure S2: *Zic1* expression defines the boundary of dI4/dILA and dI5/dILB, related to Figure 1.**

(A) Expression of Zic1 in embryonic day (E) 10.5, E11.5 and E12.5 spinal cords. Zic1 is expressed in dorsal progenitors throughout neurogenesis, appears in newborn dIL neurons (Lbx1+) at E11.5, and is expressed in varying degrees throughout dIL neurons at E12.5. Notably it is absent from dI4-6 neurons marked by Lbx1 at E10.5, as well as dI1 neurons marked by Lhx2.
(B-D) UMAP plots of single-cell RNA-sequencing data from E9.5 – E13.5 spinal cords from Delile et al, 2019 and Osseward et al, 2021, encompassing the timeframe of (A), showing *Zic1* expression in dIL neurons but not dI4/5 neurons. (B) Cluster annotations from Delile et al., 2019. (C) Normalized expression of *Zic1*. (D) Re-annotation of the Delile data based on our analysis. (E) Dot plot showing that the earliest-born dorsal horn neurons (dI4/dI5) by TTF expression do not express *Zic1*, while those bearing dILA1/2 and dILB1/2 markers do. Dot size represents the fraction of a cell type expressing a gene, and dot shading represents the average normalized expression of a gene per cell type.
Numbers: Representative images from n=3 spinal cords from E10.5, E11.5 and E12.5. Single-cell RNA-sequencing cells: Delile et al., 2019 (N=356 E9.5, N=1724 E10.5, N=3588 E11.5, N=4561 E12.5, N=5542 E13.5); Osseward et al., 2021 (N=2914 E12.5).
Scale bars: (A) 250 mm (top row), 50 mm (bottom row).
Abbreviations: DREZ – dorsal root entry zone, MZ – marginal zone, SVZ – subventricular zone, TTF – temporal transcription factor, UMAP – uniform manifold approximation and projection, VZ – ventricular zone.

Figure S3: Transcriptomic analysis of post-neurogenesis embryonic spinal cord neurons

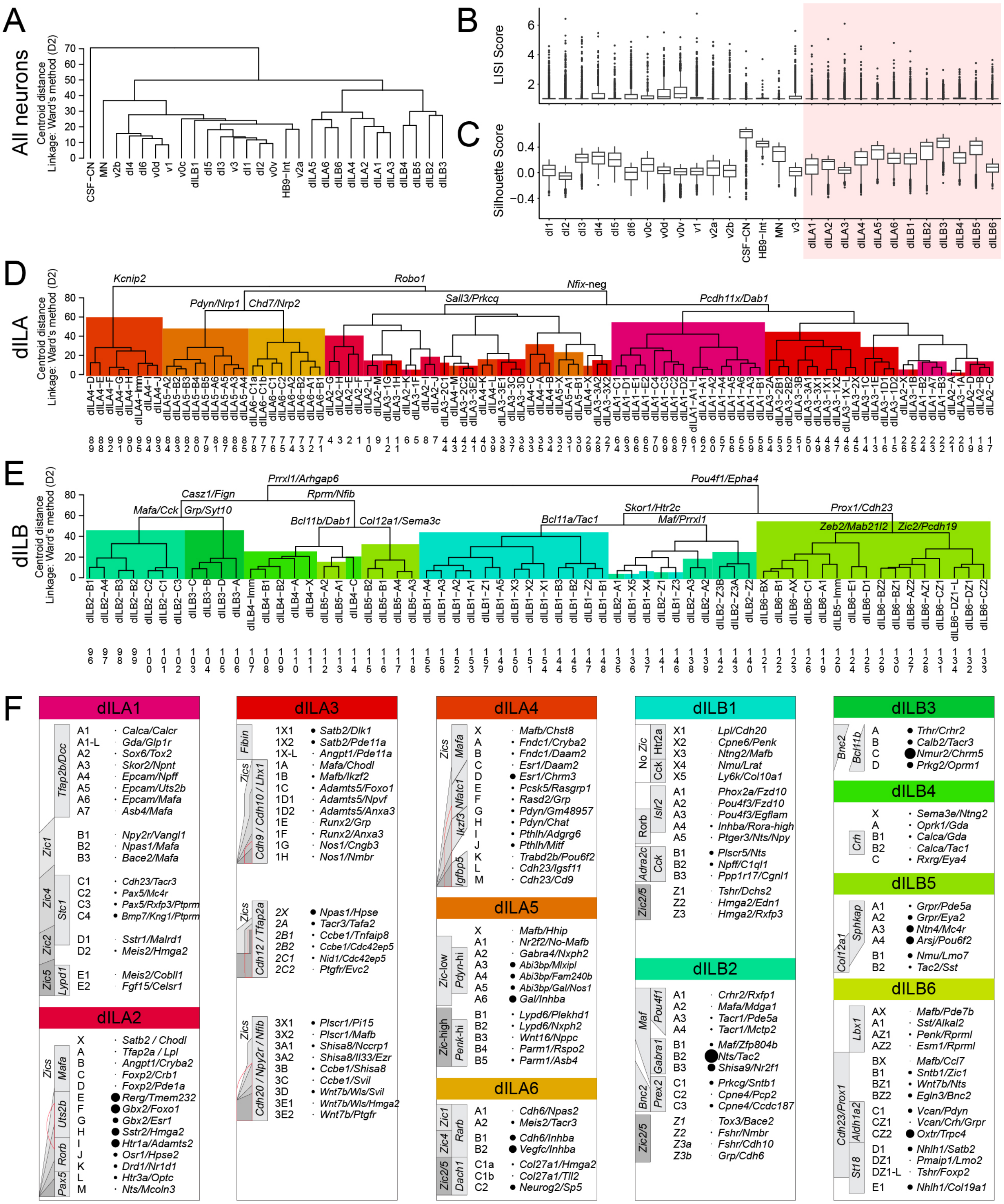

**Figure S3: Transcriptomic analysis of post-neurogenesis embryonic spinal cord neurons, related to Figure 1.**

A) Hierarchical relationships of all major spinal cord neuron classes by cluster centroid distance in PC space. Dendrograms were produced by Ward's method (D2), with use of the top 90 PCs. (B,C) Cluster robustness metrics for major spinal cord neuron classes, using LISI (B) and silhouette (C) scores. dIL neurons are indicated by red background. (D,E) Hierarchical relationships of dILA (D) neuron subtypes and dILB (E) neuron subtypes within each respective class by cluster centroid distance in PC space. Dendrograms were produced by Ward's method (D2), with use of 90 PCs in both cases. Colors represent major dILA and dILB families. High-order branches are annotated with top DEGs. Numbers refer to terminal dendrogram branches in Figure 1F which corresponding to subtype names indicated on terminal dendrogram branches here. F) Annotated tables for each of twelve dIL families, showing subtypes and top marker genes. The "-L" suffix represents neuron subtypes which are virtually only found samples taken from the lumbosacral cord; no outstanding neuron subtypes are exclusive to cervical or thoracic segments given the current number of dIL neurons sampled. Dot size next to subtype name corresponds to the relative frequency of dIL neuron subtype among subtypes of all six families (excluding clusters annotated as transitional or immature (those overrepresented in E14.5 data or with substantial *Robo3*/*Nhlh1* expression).
Numbers: Single-cell RNA-sequencing data analyzed here was derived from 94,048 neurons (see Figure 1C and methods for details).
Abbreviations: DEG – differentially-expressed gene, PC – principal component

Figure S4: Robustness of spinal neuron clusters using a random forest classifier

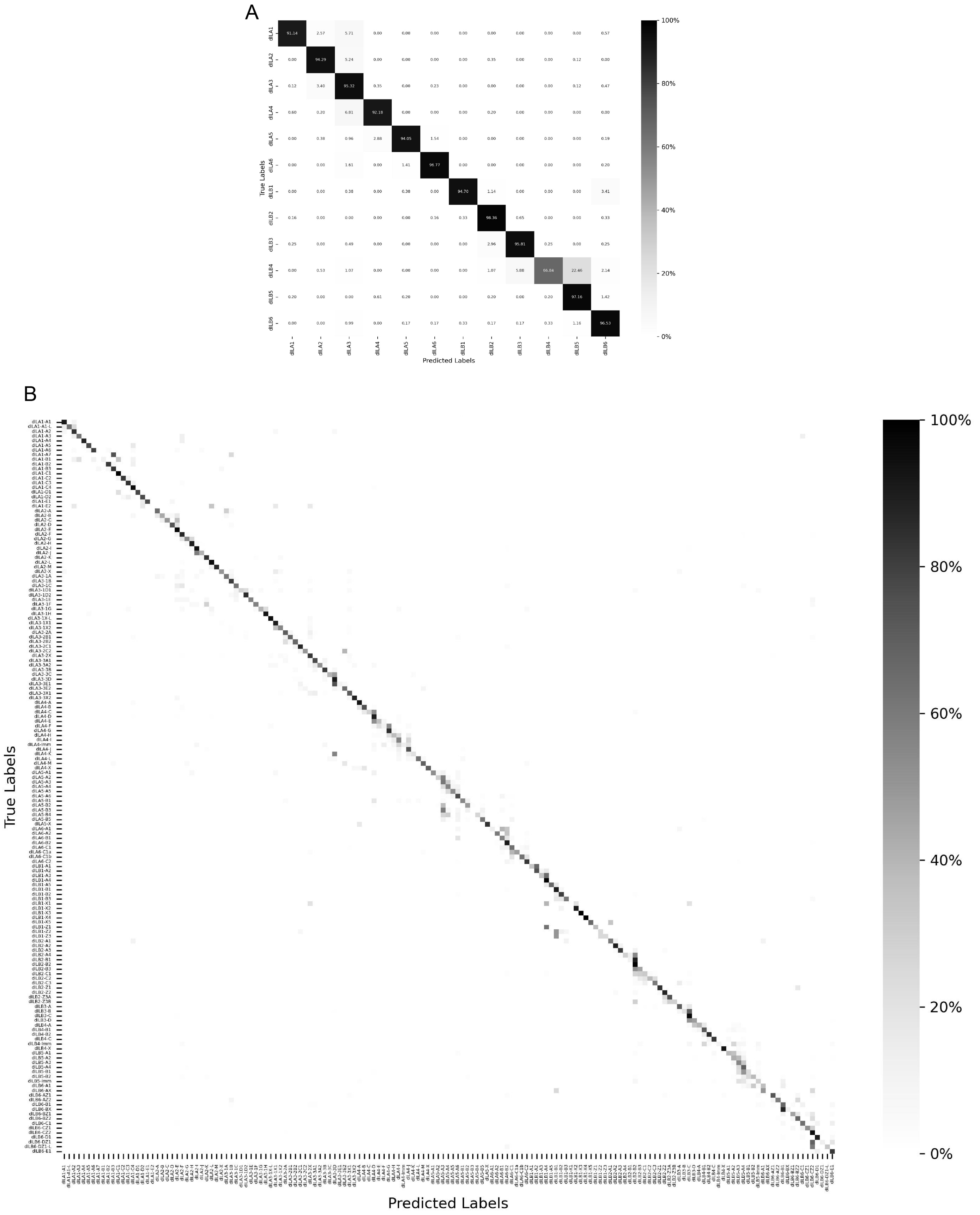

**Figure S4: Robustness of spinal neuron clusters using a random forest classifier, related to Figure 1.**

Confusion matrix heatmaps demonstrating the accuracy of a trained random forest classifier in predicting either family-level (A) or family subtype-level (B) annotations for individual dIL neurons using single-cell RNA-sequencing data generated and annotated here.

Figure S5: UMAP plot of dIL family subtypes

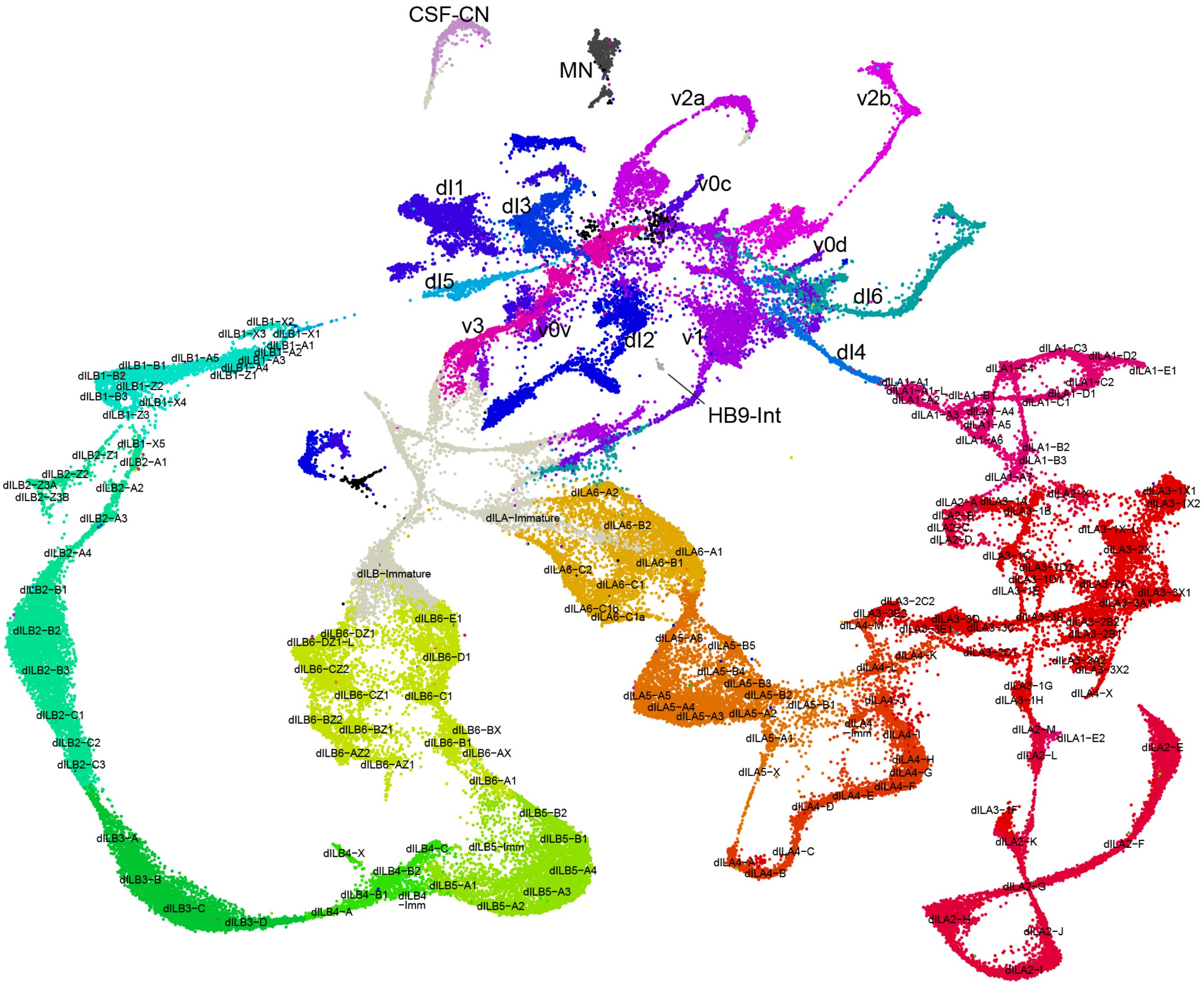

**Figure S5: UMAP plot of dIL family subtypes, related to Figure 1.**

UMAP plot of all spinal cord neurons, with the approximate UMAP centroid of each dIL neuron type annotated. Cardinal-class neurons are annotated in larger text.
Abbreviations: UMAP – uniform manifold approximation and projection.

Figure S6: EdU birthdating of dILA families

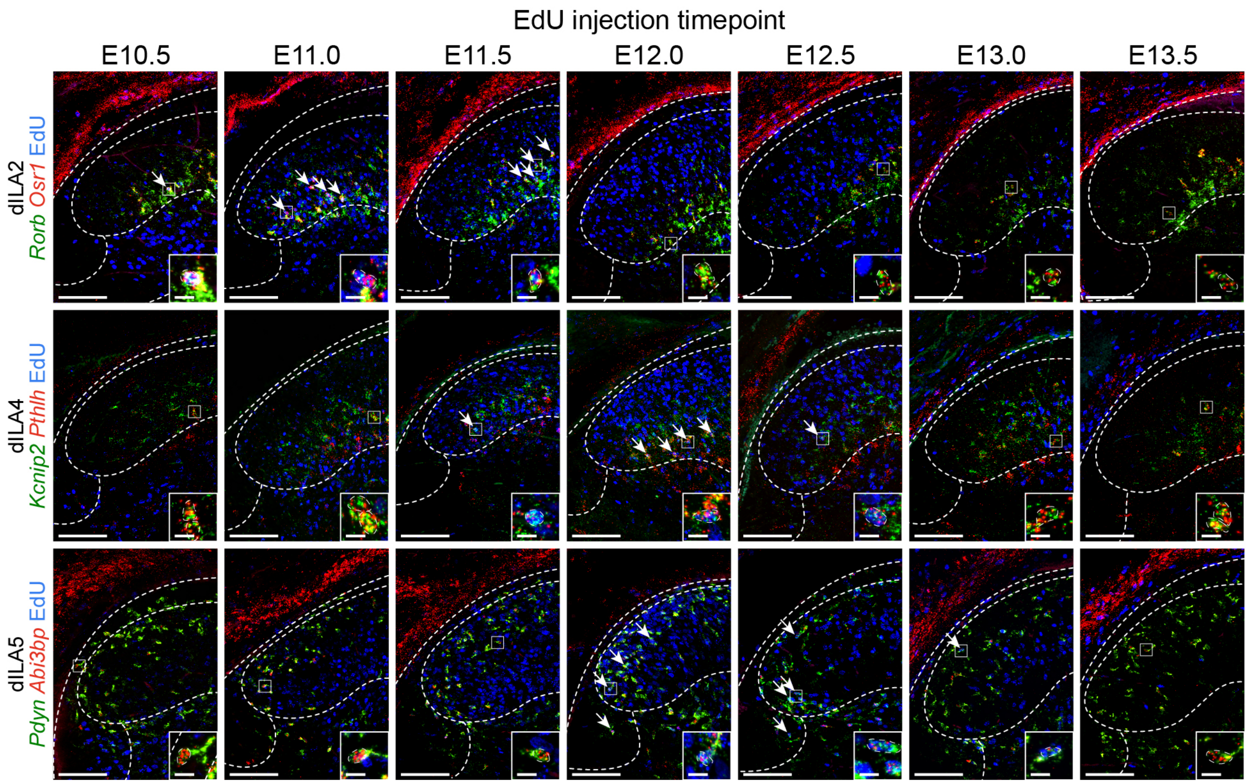

**Figure S6: EdU birthdating of dILA families, related to Figure 2.**

Representative images of the lateral dorsal horn of the spinal cord from E16.5 embryos having received EdU pulses at either E10.5, E11.0, E11.5, E12.0, E12.5, E13.0 or E13.5 respectively. A subset of dILA families are identified by coincident expression of two genes (red and green) revealed by RNA Scope in-situ hybridization which exclude all other dIL neuron families; dILA2 – *Rorb* and *Osr1*, dILA4 – *Kcnip2* and *Pthlh*, dILA5 – *Pdyn* and *Abi3bp*. Arrows indicate dILA family neurons expressing the corresponding marker gene set and which are labeled with nuclear EdU (blue). Dashed lines indicate the border of the superficial dorsal horn with the deep dorsal horn and the lateral spinal nucleus, the lateral spinal nucleus with the lateral funiculus, and the pia. Insets indicate examples of triple-positive neurons surrounded by dashed lines. Example images from this figure used in Figure 2E are reused here for contextual purposes.
Numbers: Individual litters of embryos from a mother given a single EdU injection were considered a single biological replicate. E10.5 (n=5 litters), E11.0 (n=4 litters), E11.5 (n=5 litters), E12.0 (n=5 litters), E12.5 (n=5 litters), E13.0 (n=5 litters), E13.5 (n=4 litters).
Scale bars: 250 $\mu$m, 25 $\mu$m for inset panels.
Abbrevations: EdU - 5-ethynyl-2'-deoxyuridine.

Figure S7: EdU birthdating of dILB families

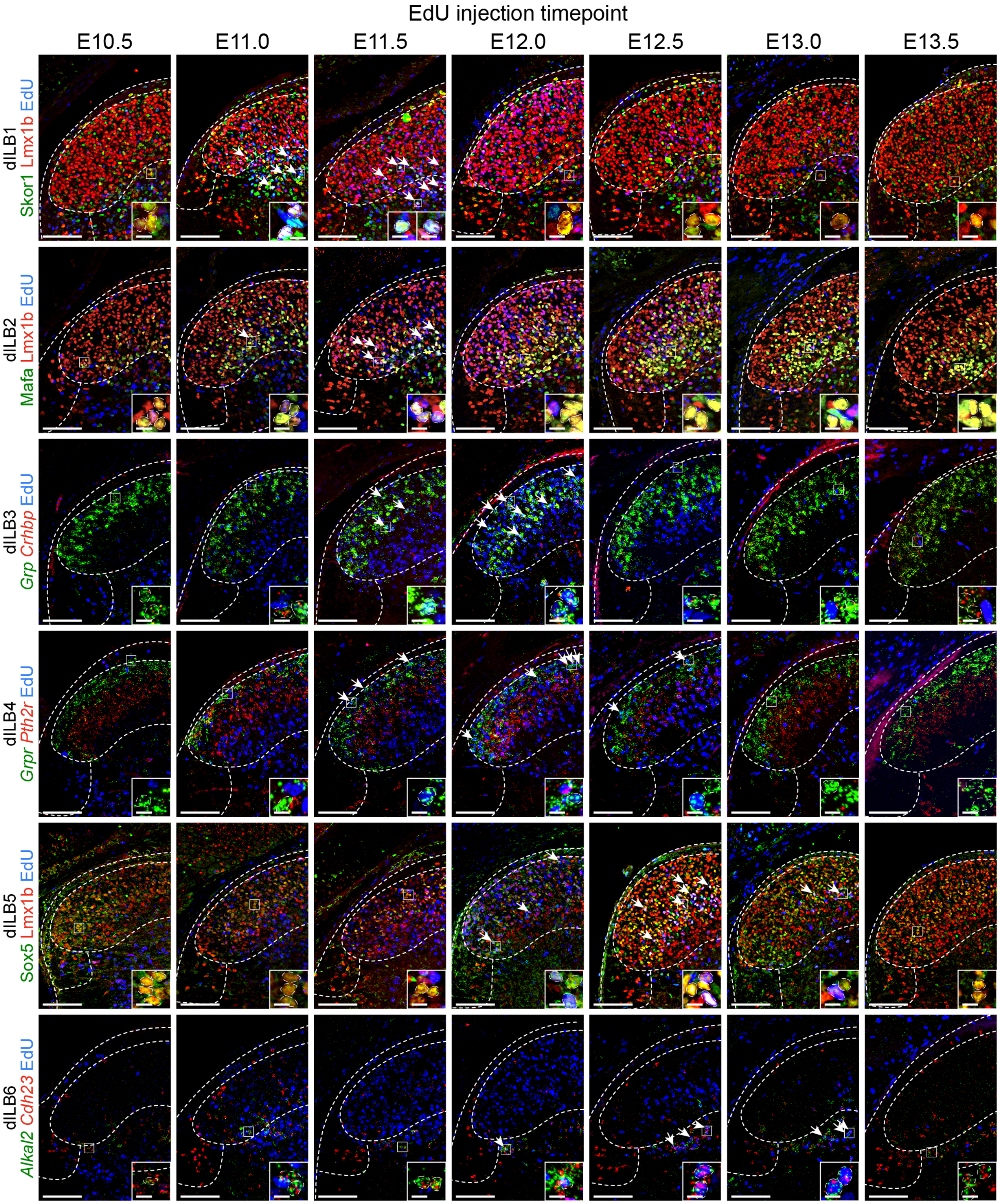

**Figure S7: EdU birthdating of dILB families, related to Figure 2.**

Representative images of the lateral dorsal horn of the spinal cord from E16.5 embryos having received EdU pulses at either E10.5, E11.0, E11.5, E12.0, E12.5, E13.0 or E13.5 respectively. A subset of dILB families are identified by coincident expression of two genes (red or green) revealed by RNA Scope in-situ hybridization or nuclear protein expression; dILB1 – Lmx1b and Skor1, dILB2 – Lmx1b and Mafa, dILB3 – *Grp* and *Crhbp*, dILB4 – *Grpr* excluding *Pth2r*, dILB5 – Lmx1b and Sox5, dILB6 – *Cdh23* and *Alkal2*. Arrows indicate dILB family neurons expressing the corresponding marker gene set and which are labeled with nuclear EdU (blue). Dashed lines indicate the border of the superficial dorsal horn with the deep dorsal horn and the lateral spinal nucleus, the lateral spinal nucleus with the lateral funiculus, and the pia. Insets indicate examples of triple-positive neurons surrounded by dashed lines. Example images from this figure used in Figure 2E are reused here for contextual purposes.
Numbers: Individual litters of embryos from a mother given a single EdU injection were considered a single biological replicate. E10.5 (n=5 litters), E11.0 (n=4 litters), E11.5 (n=5 litters), E12.0 (n=5 litters), E12.5 (n=5 litters), E13.0 (n=5 litters), E13.5 (n=5 litters).
Scale bars: 250 μm, 25 μm for inset panels.
Abbrevations: EdU - 5-ethynyl-2'-deoxyuridine.

Figure S8: EdU birthdating of dILB neurons in a cohort unpaired with in-situ hybridization

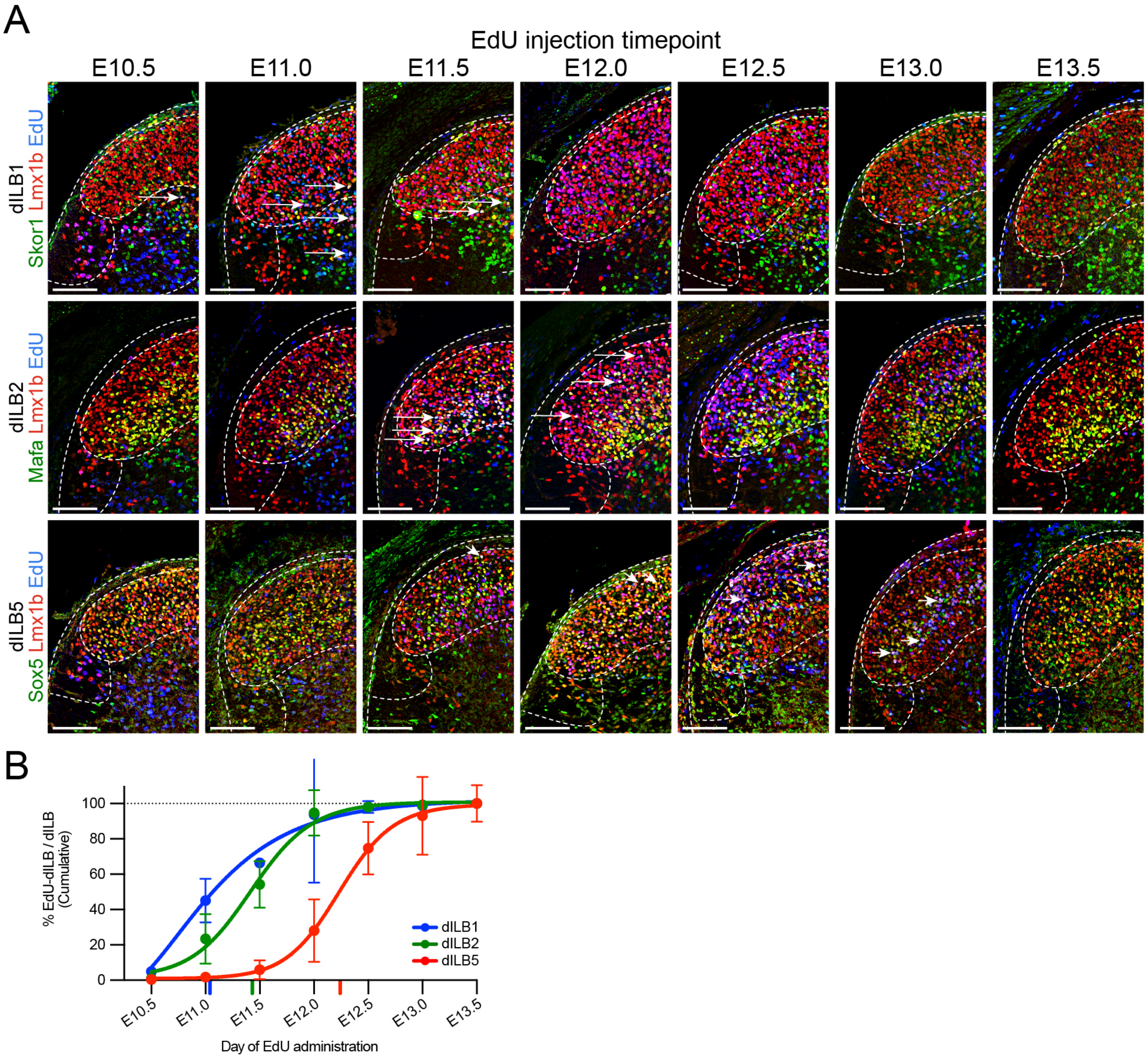

**Figure S8: EdU birthdating of dILB neurons in a cohort unpaired with in-situ hybridization, related to Figure 2.**

Embryos from birthdated litters where separate embryos were not appropriately fixed for in-situ hybridization were only examined for dILB families identifiable using immunohistochemistry (dILB1, dILB2 and dILB5). This experiment was supplanted by those in Figures 2, S7 and S8, and thus not all litters were collected in triplicate prior to conducting subsequent experiments. Despite this, these experiments provide independent support the same birth order for dILB1, dILB2 and dILB5.
(A) Representative images of the lateral dorsal horn of the spinal cord from E16.5 embryos having received EdU pulses at either E10.5, E11.0, E11.5, E12.0, E12.5, E13.0 or E13.5 respectively. A subset of dILB families are identified by coincident expression of two nuclear proteins (red and green) which exclude all other dIL neuron families; dILB1 – Lmx1b and Skor1, dILB2 – Lmx1b and Mafa, dILB5 – Lmx1b and Sox5. Arrows indicate dILB family neurons expressing the corresponding marker gene set and which are labeled with nuclear EdU (blue). Dashed lines indicate the border of the superficial dorsal horn with the deep dorsal horn and the lateral spinal nucleus, the lateral spinal nucleus with the lateral funiculus, and the pia. Insets indicate examples of triple-positive neurons surrounded by dashed lines.
(B) Plot showing the cumulative labeling of dILB1, dILB2 or dILB5 neurons by EdU over subsequent embryonic days. Points represent cumulative EdU labeled for each dILB family; individual data points are mean cumulative EdU labeling for all embryos of a single litter. Lines represent best-fit sigmoidal relationships of means of each dILB family.
Numbers: Images and cell counts shown here are derived from the same cohort of embryos analyzed in Figures 1A, 1B, 3A and 3B; however, here data from all embryos in a single litter given a single EdU injection were averaged and considered a single biological replicate. E10.5 (n=2-3 litters), E11.0 (n=3 litters), E11.5 (n=4 litters), E12.0 (n=3 litters), E12.5 (n=4 litters), E13.0 (n=3 litters), E13.5 (n=4 litters).
Scale bars: 250 µm.
Abbrevations: EdU - 5-ethynyl-2'-deoxyuridine.

Figure S9: Spatial transcriptomics reveal the laminar position of dIL family subtypes

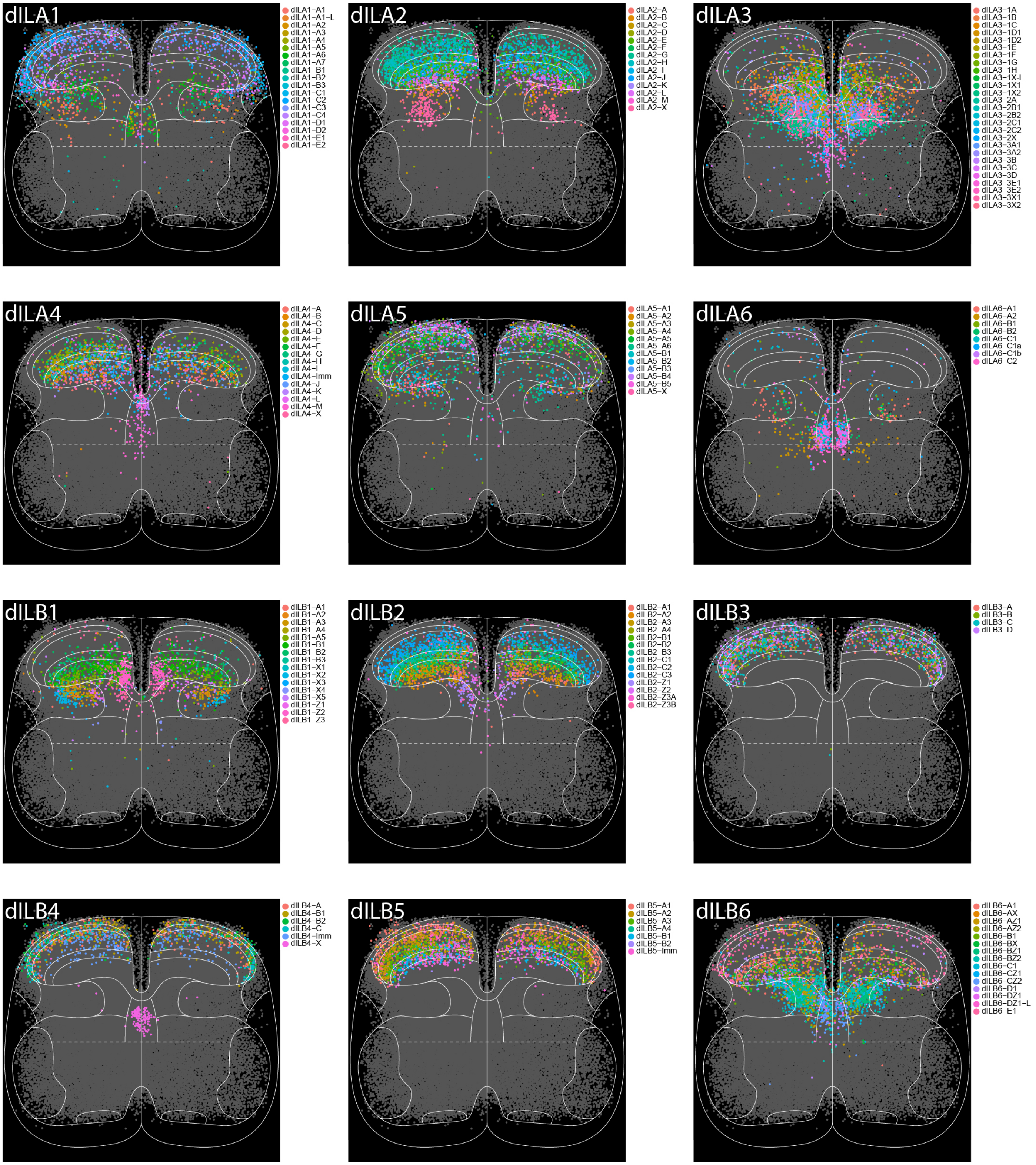

**Figure S9: Spatial transcriptomics reveal the laminar position of dIL family subtypes, related to Figure 3.**

Scatter plots depict the coordinates of all wild-type brachial (C5-C8) E16.5 neurons examined in the Xenium spatial transcriptomics assay. Cell-type identity was predicted using RCTD, with single-cell RNA-sequencing data from Figure 1 as a reference. Cell coordinates were normalized to anatomic landmarks from each profiled spinal cord section; subsequently, the plane of cell coordinates was rotated and stretched to fit an ideal diagram of a spinal cord (white solid lines in each panel, with approximate laminar boundaries in dashed white lines). This process was identical for all cells regardless of annotation. Cells assigned to each of 6 dILA and 6 dILB families are plotted separately, as indicated in the top left of each panel. Cells not belonging to the indicated dIL family are plotted in gray; otherwise, they are colored based on their family subtype identity in the corresponding legend to each panel.
Numbers: Cell coordinates were derived from six whole sections and seven hemisections (N=9.5 sections) among n=3 replicate wild-type E16.5 spinal cords. From these sections 54,166 cells were identified as neurons and plotted, of which 27,714 were annotated as dIL neurons.
Abbreviations: RCTD – Robust cell-type decomposition (R library).

Figure S10: dILB neurons laminate normally in absence of dILA neurons – images

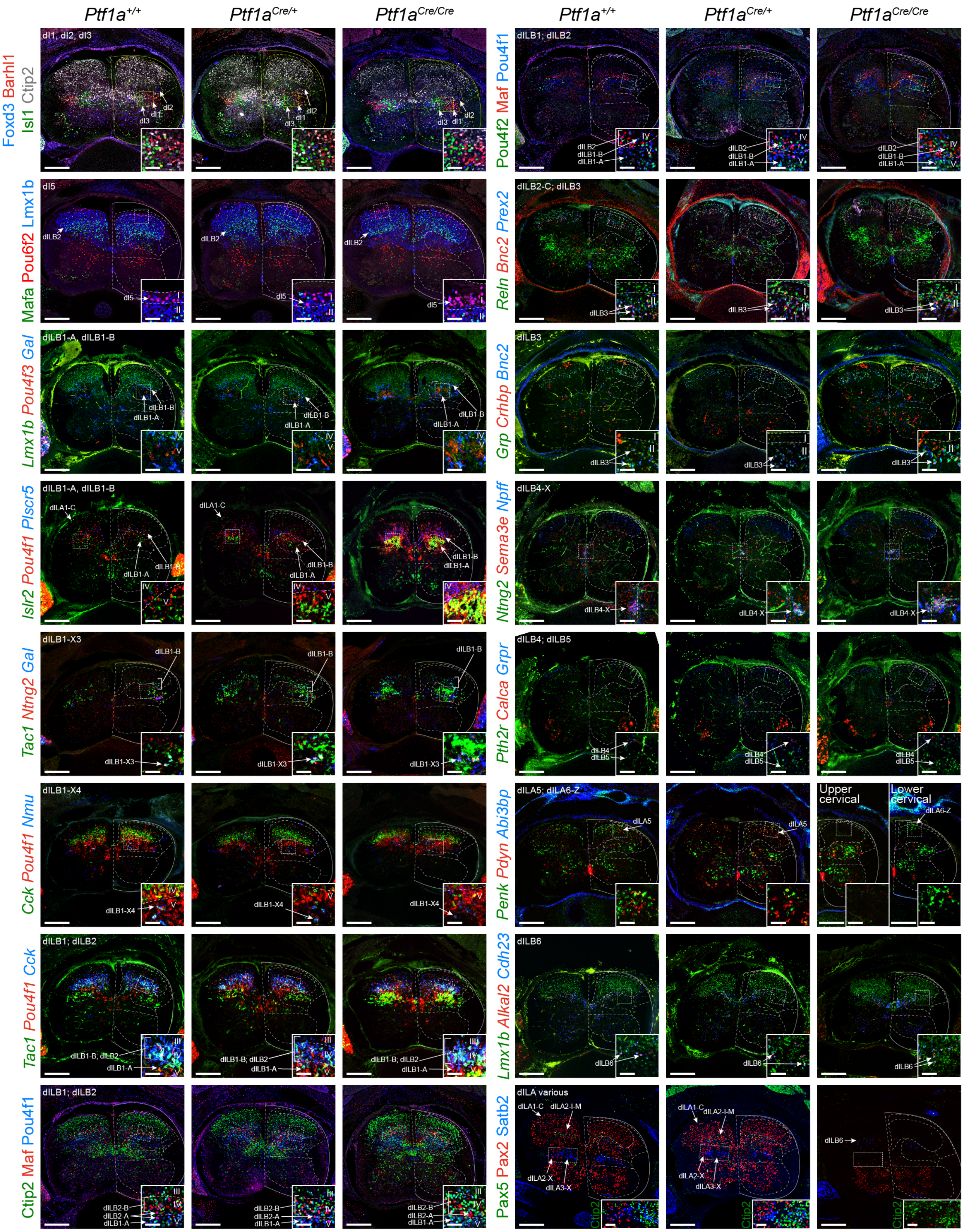

**Figure S10: dILB neurons laminate normally in absence of dILA neurons – images, related to Figure 4.**

Images of immunohistochemistry and RNA Scope in-situ hybridization of various cardinal excitatory, dILB dorsal horn neuron types and selected dILA populations (indicated in the top left of each row) are examined in E16.5 spinal cords from embryos wild-type, heterozygous-null, or null for *Ptf1a*. dILA neurons are largely absent from null spinal cords while excitatory neurons are anatomically similar to controls (though present in increased numbers). White arrows indicate examples of dIL family subtypes. Insets at the bottom right of each panel are magnified fields of view from boxes marked by dashed white lines in the larger panel. Other solid and dashed white lines indicate approximate anatomical boundaries of the spinal cord. Images of *Plscr5* (row 4, left) are reused here from Figure 4, for contextual purposes.
Numbers: Representatives images from n=4 *Ptf1a*$^{+/+}$; n=4 *Ptf1a*$^{Cre/+}$; n=4 *Ptf1a*$^{Cre/Cre}$ E16.5 spinal cords.
Scale bars: 250 µm, 50 µm for inset panels.

Figure S11: Transdifferentiation of dIL neurons in *Ptf1a*- and *Gsx1/2*-null embryos

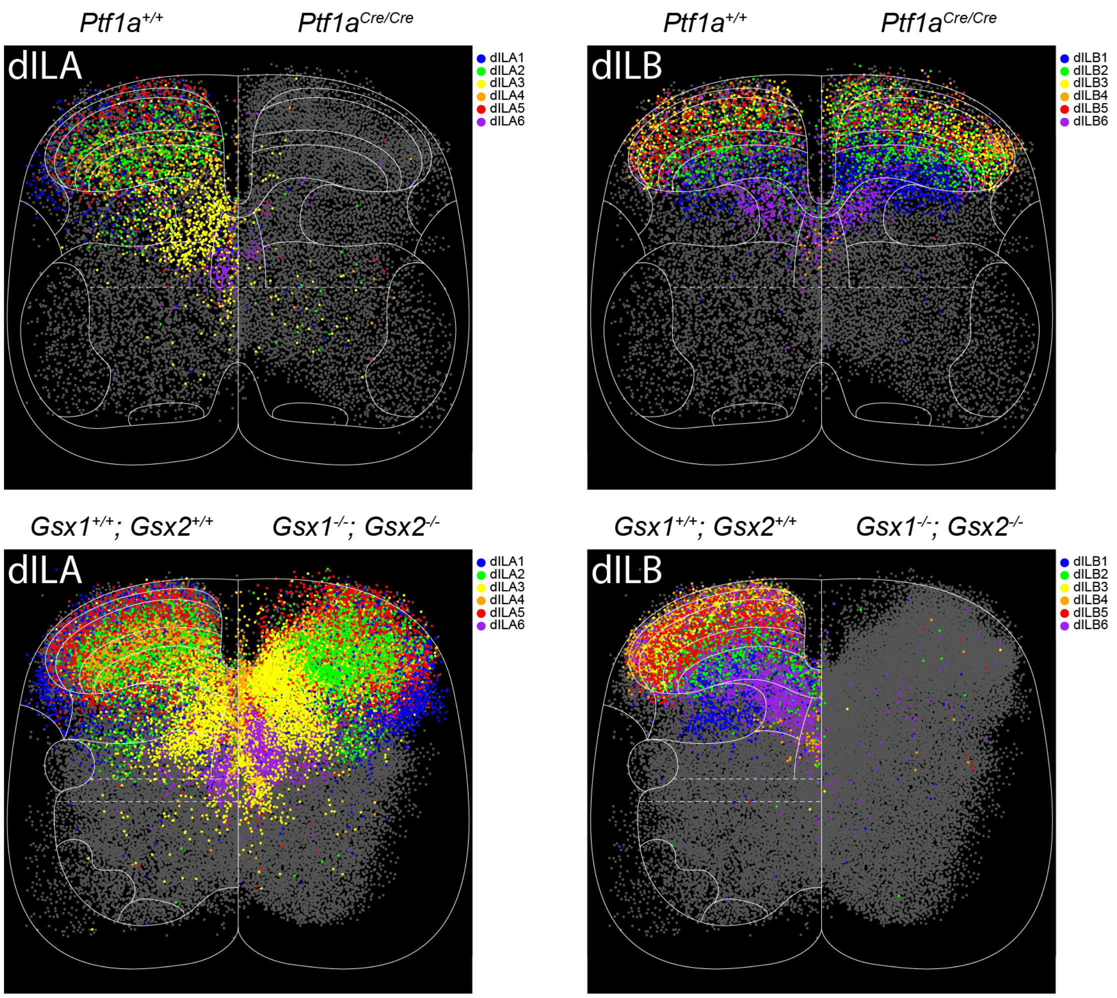

**Figure S11: Transdifferentiation of dIL neurons in *Ptf1a*- and *Gsx1/2*-null embryos, related to Figure 4.**

Scatter plots depict the coordinates of wild-type, *Ptf1a*-null or *Gsx1/2*-null E16.5 neurons examined in the Xenium spatial transcriptomics assay, colored by either dILA family (column 1) or dILB family (column 2). Here, segmentally-matched wild-type cell coordinates have been mirrored and overlaid on the left side of each diagram and segmentally corresponding cell coordinates from either *Ptf1a*-null tissue (row 1) or *Gsx1/2*-null tissue (row 2) have been mirrored and overlaid on the right side of each diagram in order to facilitate anatomical comparisons between genotype. *Ptf1a*-null cells have been compared to brachial wild-type cell coordinates displayed in Figure S9 while *Gsx1/2*-null tissue analyzed from upper thoracic segments have been compared to wild-type thoracic cell coordinates not previously displayed. Wild-type neurons have been evenly downsampled from each neuron subtype annotation to match the total number of null cells displayed per diagram. Notably, dILA neurons are absent from Ptf1a-null tissue and dILB neurons are absent from *Gsx1/2*-null tissue. Cells not belonging to the indicated dIL family are plotted in gray. Solid and dashed white lines indicate approximate anatomical boundaries of the spinal cord.
Numbers: Spatial transcriptomics from E16.5 wild-type embryo brachial segments was previously described (Figure S9). For E16.5 wild-type thoracic segments, cell coordinates were derived from five whole sections and one hemisection (N=5.5 sections), among n=3 replicate spinal cords. From these sections 27,758 cells were identified as neurons and plotted, of which 13,724 were annotated as dIL neurons. For E16.5 *Ptf1a*-null brachial segments, cell coordinates were derived from two whole sections (N=2 sections) among n=1 spinal cord. From these sections, 8,267 cells were identified as neurons, and of which 2,767 were dIL neurons. E16.5 brachial data was downsampled to 8,106 neurons of which 4,151 were dIL neurons to match. For E16.5 *Gsx1/2*-null thoracic segments, cell coordinates were derived from four whole sections and three hemisections (N=5.5 sections) among n=3 replicate spinal cords. From these sections, 24,737 cells were identified as neurons, and of which 12,217 were dIL neurons. E16.5 thoracic data was downsampled to 24,583 neurons with, also, 12,217 dIL neurons, to match.

Figure S12: dILB neurons laminate normally in absence of dILA neurons – Xenium

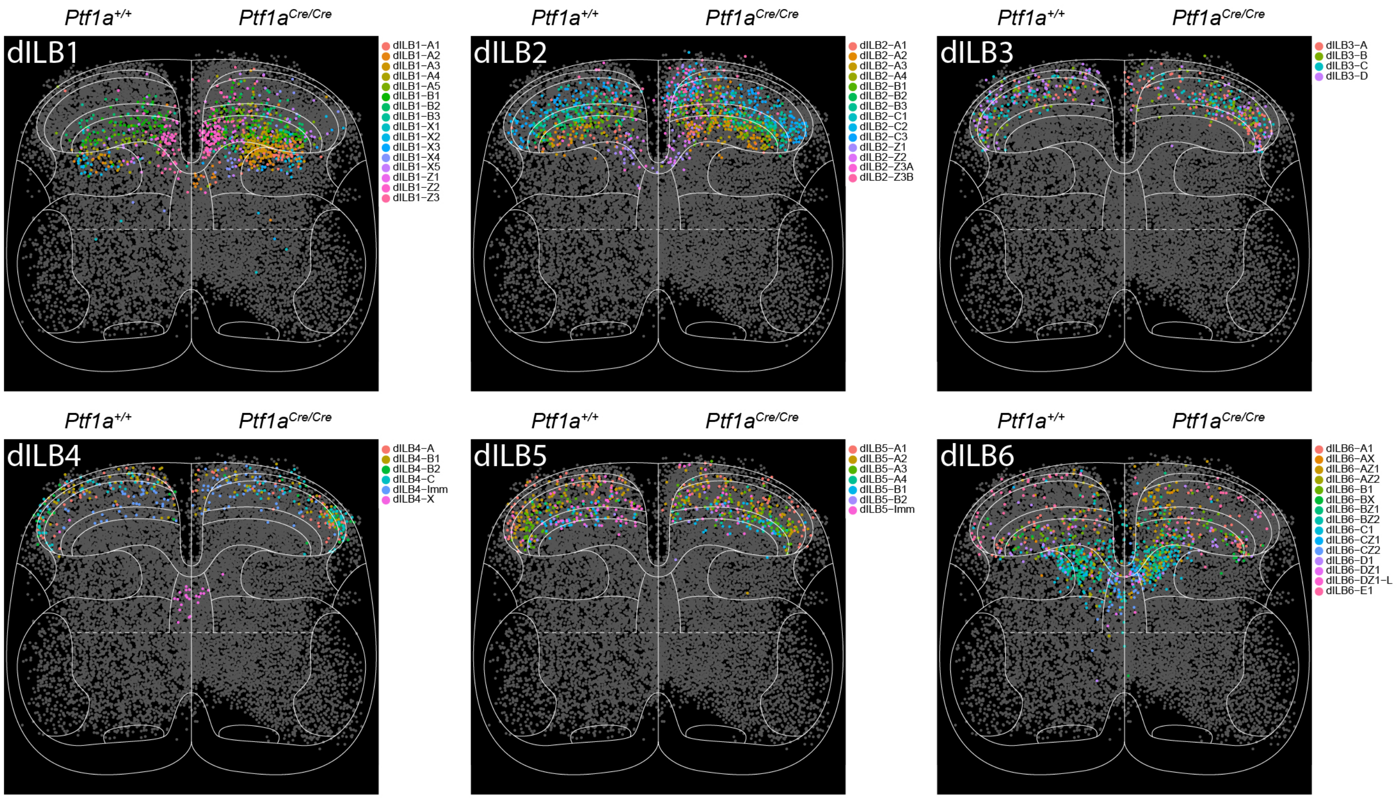

**Figure S12: dILB neurons laminate normally in absence of dILA neurons – Xenium, related to Figure 4.**

Scatter plots depict the coordinates of wild-type and *Ptf1a*-null E16.5 neurons examined in the Xenium spatial transcriptomics assay, sorted into each of six dILB families and colored by subtype annotation. As described previously, wild-type cells are displayed on the left and null cells on the right of each diagram, and wild-type cells were downsampled to match the number of null cells. All cells presented here are also presented in Figure 4 and S11 with family annotations. Cells not belonging to the indicated dIL family are plotted in gray. Solid and dashed white lines indicate approximate anatomical boundaries of the spinal cord.

### Figure S13: dILA neurons lose laminar patterning in absence of dILB neurons – images

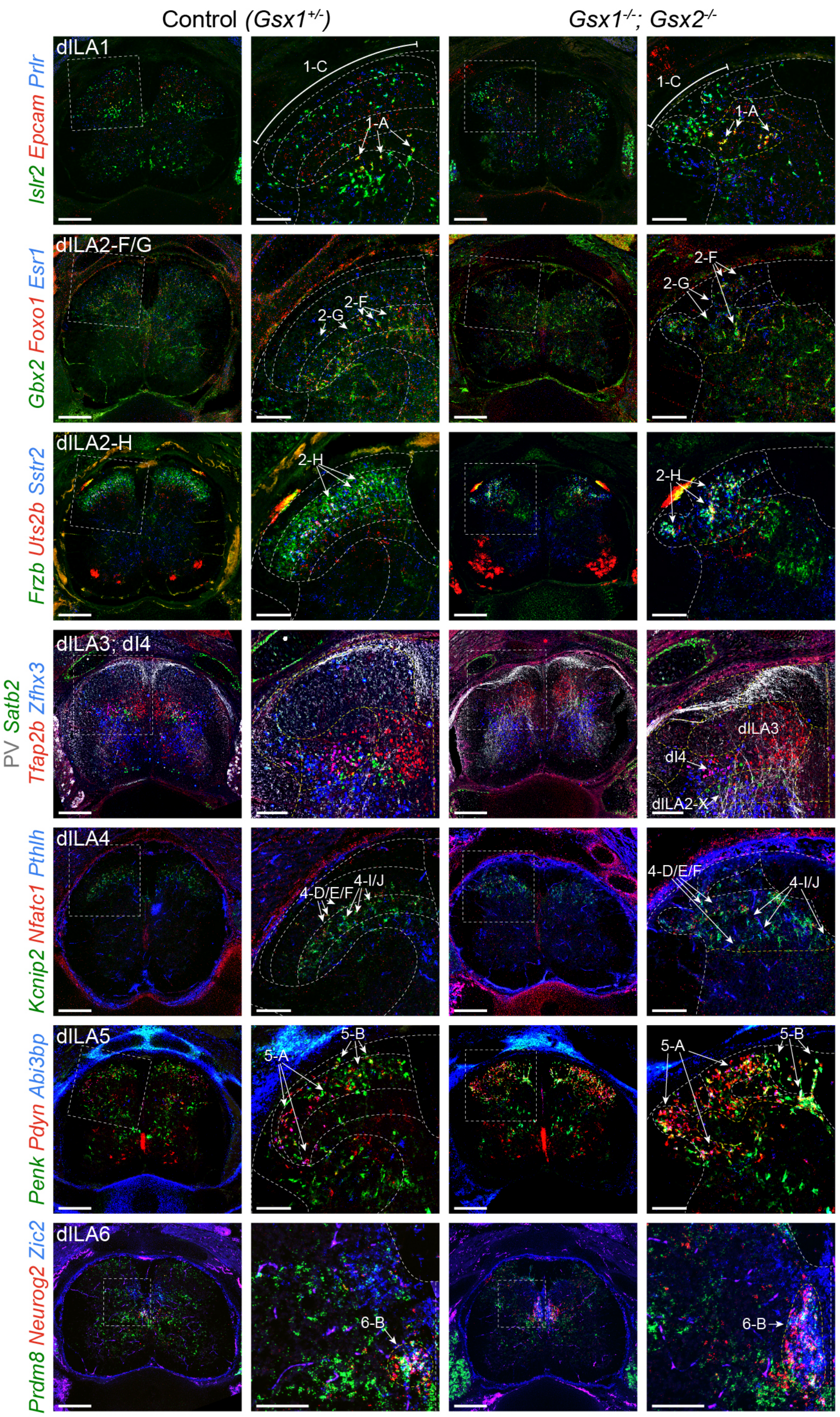

**Figure S13: dILA neurons lose laminar patterning in absence of dILB neurons – images, related to Figure 4.**

Images of immunohistochemistry and RNA Scope in-situ hybridization of dILA family subtypes (indicated in the top left of each row) are examined in E16.5 spinal cords from embryos heterozygous-null for *Gsx1* (as control), or null for both *Gsx1* and *Gsx2*. dILA neurons normally present in laminae are no longer present in laminae and are largely disorganized. Unlaminated dILA neurons in the deep dorsal horn are slightly displaced, though in anatomically similar locations. dILB neurons are largely absent from null spinal cords. White arrows indicate examples of dIL family subtypes. For each genotype, the left column of images depicts an entire spinal cord section while the right column depicts a magnified image from the dashed white line box in the left column. Other solid and dashed white lines indicate approximate anatomical boundaries of the spinal cord.
Numbers: Representative images from n=4 *Gsx1*$^{+/-}$; *Gsx2*$^{+/+}$, n=4 *Gsx1*$^{-/-}$; *Gsx2*$^{-/-}$.
Scale bars: 250 μm for left column, 100 μm for magnified panels (right column) for each genotype.
Images of PV (row 4) are reused here from Figure 4, for contextual purposes.

### Figure S14: dILA neurons lose laminar patterning in absence of dILB neurons – Xenium

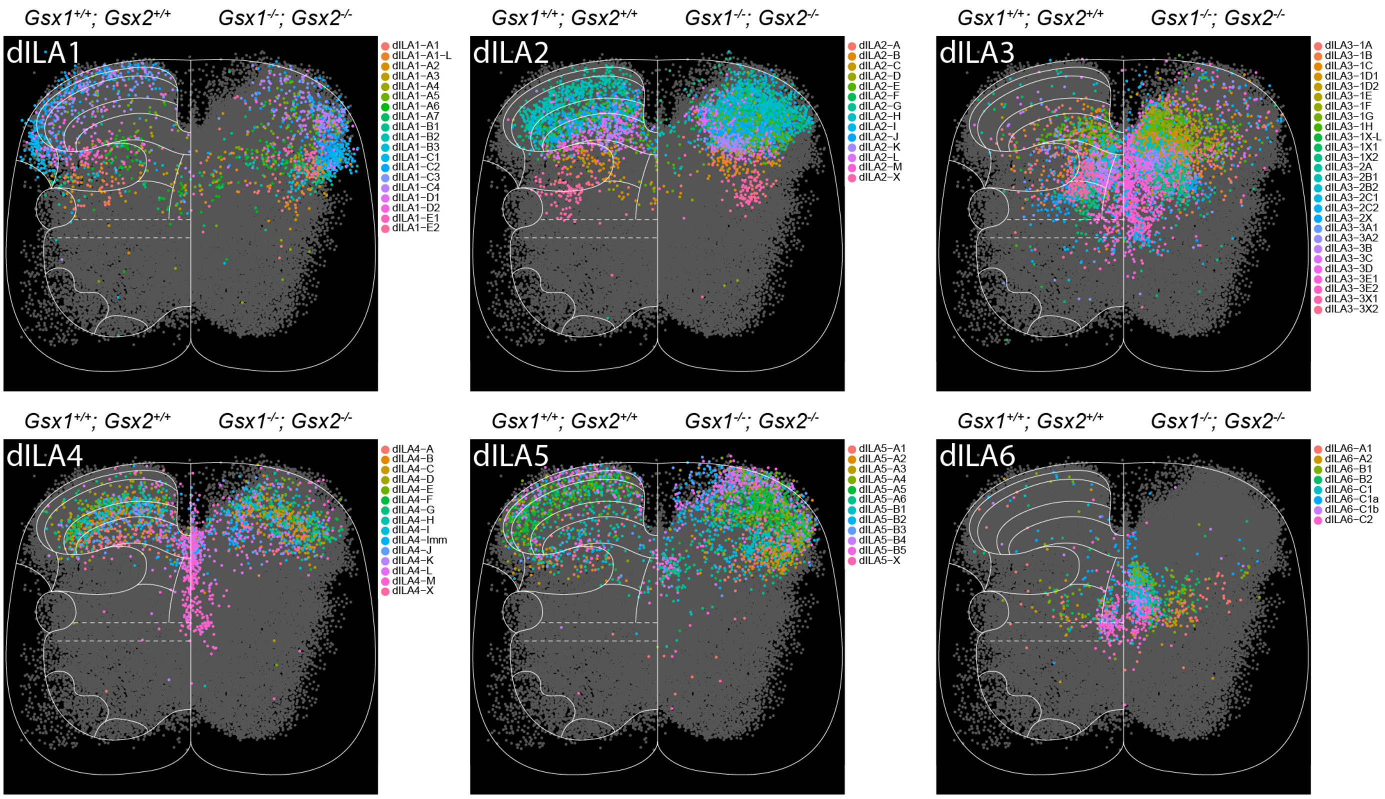

**Figure S14: dILA neurons lose laminar patterning in absence of dILB neurons – Xenium, related to Figure 4.**

Scatter plots depict the coordinates of wild-type and *Gsx1/2*-null E16.5 neurons examined in the Xenium spatial transcriptomics assay, sorted into each of six dILA families, and colored by subtype annotation. As described previously, wild-type cells are displayed on the left and null cells on the right of each diagram, and wild-type cells were downsampled to match the number of null cells. All cells presented here are also presented with family-level annotations in Figure S11 and with selected refined annotations in Figure 4. Cells not belonging to the indicated dIL family are plotted in gray. Solid and dashed white lines indicate approximate anatomical boundaries of the spinal cord.

Figure S15: Sensory neuron afferents are dispensable for dorsal horn structure

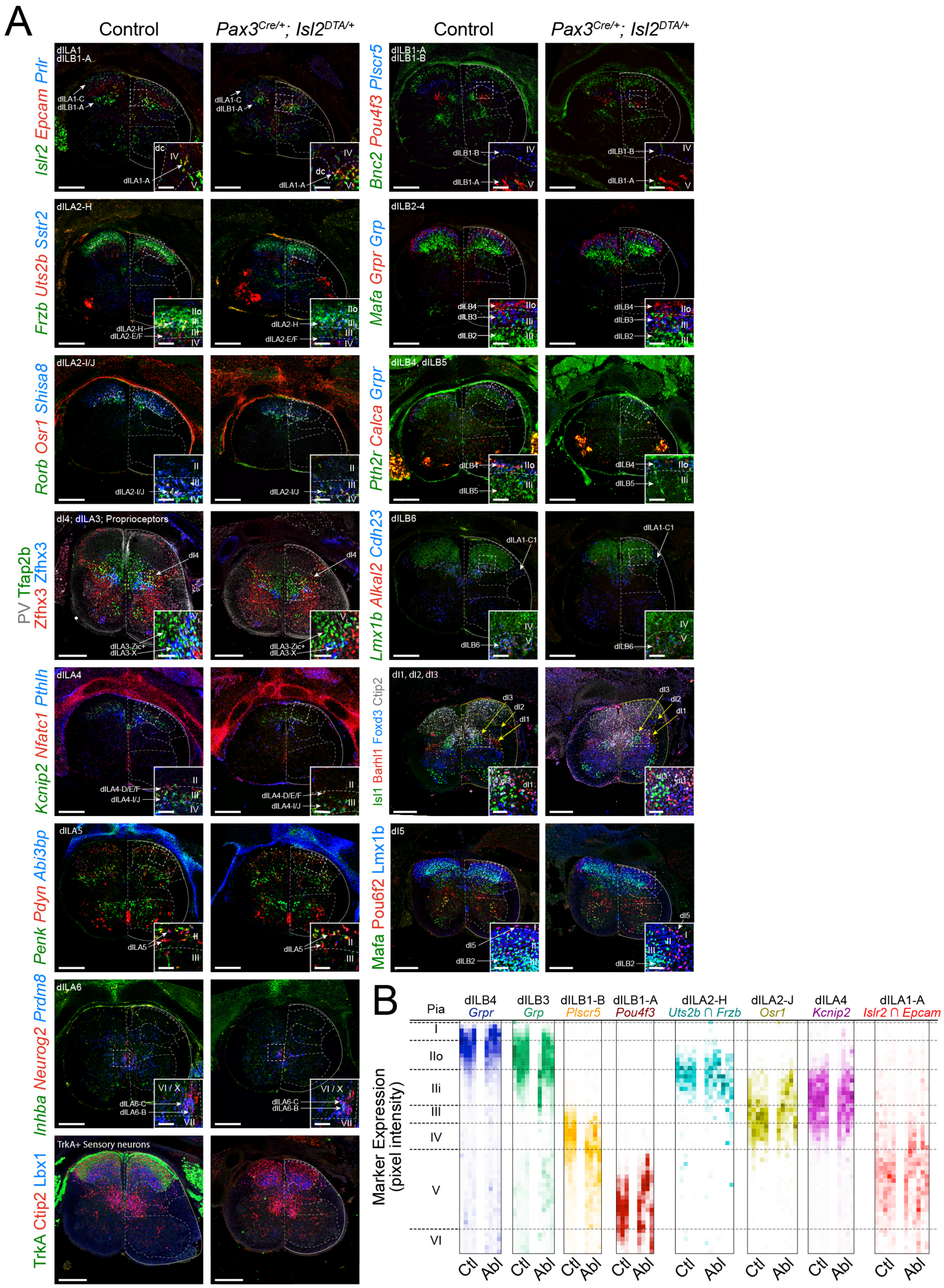

**Figure S15: Sensory neuron afferents are dispensable for dorsal horn structure, related to Figure 4.**

(A) Immunohistochemistry and RNA Scope in-situ hybridization of dILA family subtypes, dILB family subtypes, and sensory afferents (indicated in the top left of each row) are examined in E16.5 spinal cords from control littermates, (either *Pax3*$^{Cre/+}$ or *Isl2*$^{DTA/+}$) or embryos which are both *Pax3*$^{Cre/+}$ and *Isl2*$^{DTA/+}$, resulting in pan-sensory neuron ablation. DILA and dILB neurons laminate in sensory neuron-ablated spinal cord similarly to control spinal cords. Notable phenotypes are atrophy of the dorsal columns (due to a lack of sensory neuron axons), and mild atrophy of dorsal horn interneurons non-specifically when sensory neurons are absent. White arrows indicate examples of dIL family subtypes. Insets at the bottom right of each panel are magnified fields of view from boxes marked by dashed white lines in the larger panel. Other solid and dashed white lines indicate approximate anatomical boundaries of the spinal cord.
(B) Spatial distribution of individual gene or protein markers or marker sets (as indicated at the top of each box), showing the normalized proportion of total signal per dorso-ventral bin (color intensity) where the pia is set as the dorsal boundary and approximate laminar positions are shown. Biological replicates are grouped together as individual columns for either Ctl (control – either *Pax3*$^{Cre/+}$ or *Isl2*$^{DTA/+}$) or Abl (Ablated - *Pax3*$^{Cre/+}$; *Isl2*$^{DTA/+}$).
Numbers: Representative images and data from n=3-5 control (either *Pax3*$^{Cre/+}$ or *Isl2*$^{DTA/+}$), n=4-7 *Pax3*$^{Cre/+}$;*Isl2*$^{DTA/+}$ E16.5 spinal cords.
Scale bars: 250 µm, 50 µm for inset panels.

Figure S16: Expression of Zic family genes in the dorsal horn

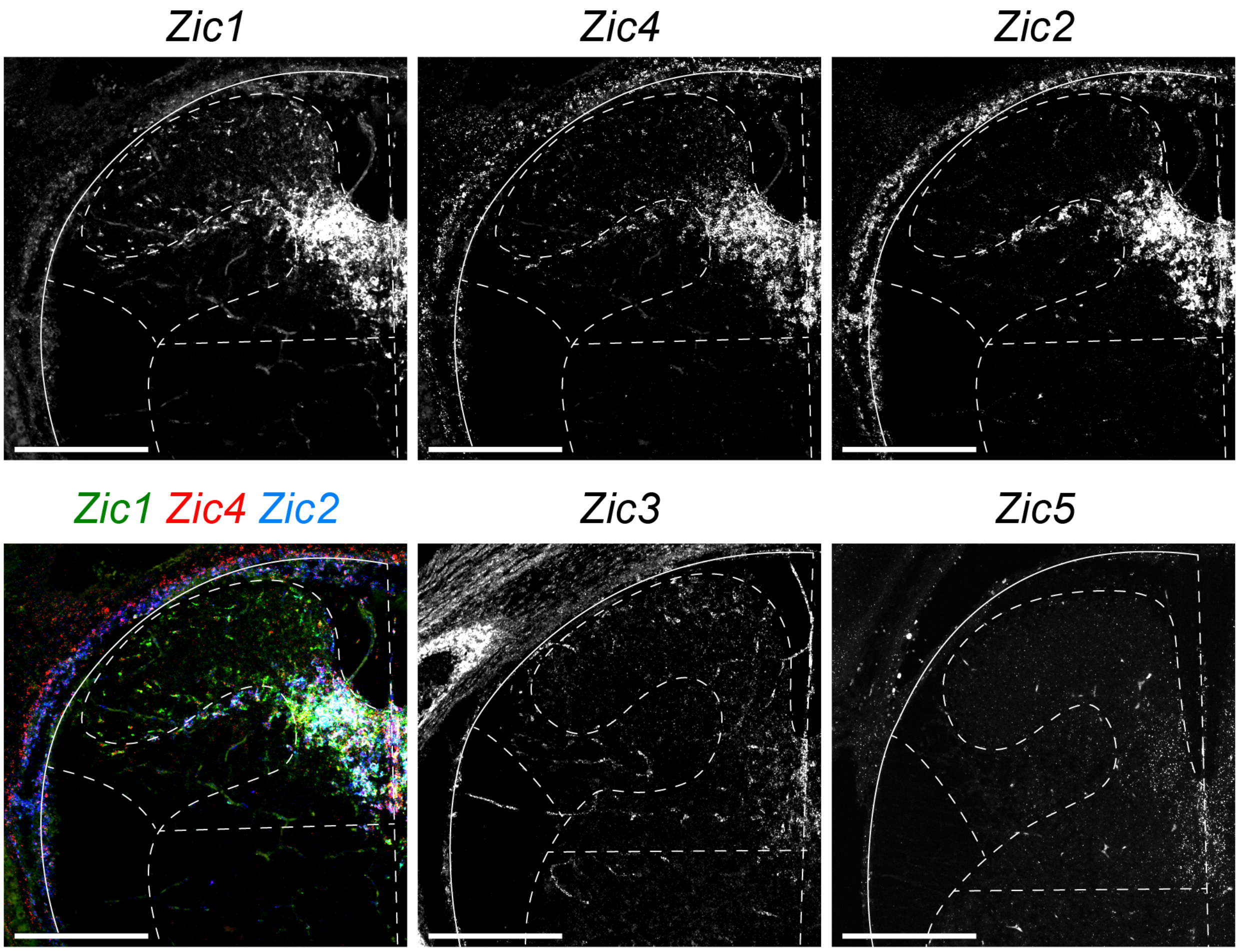


**Figure S16: Expression of Zic family genes in the dorsal horn, related to Figure 5.**

Zic1-5 genes are expression in stereotypical patterns restricted to dorsal horn neurons, with notable expression also along vasculature and meninges. The breadth and intensity of Zic gene expression corresponds well to that observed within single-cell RNA-sequencing of dIL neurons.

Numbers: Representative images from n=3 E16.5 wild-type mouse spinal cords

Scale bars: 250 µm

Figure S17: *Zic3* mildly represses formation of mid Zic-gradient subtypes of dIL neurons

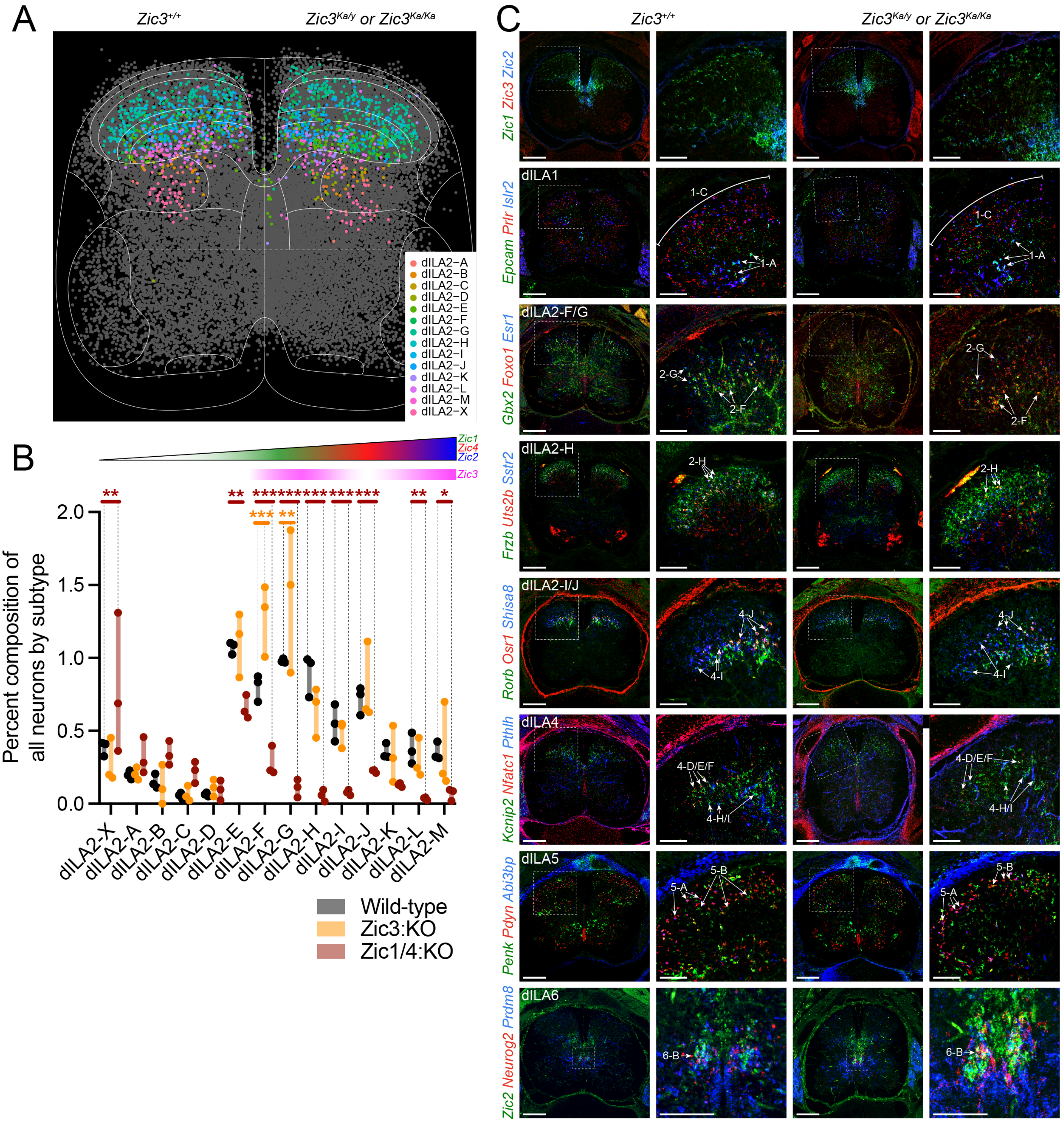

**Figure S17: *Zic3* mildly represses formation of mid Zic-gradient subtypes of dIL neurons, related to Figure 5.**

(A) The scatter plot depicts the coordinates of brachial wild-type and *Zic3*-null E16.5 neurons examined in the Xenium spatial transcriptomics assay and colored by subtype annotation of the dILA2 family (as an example). As described previously, wild-type cells are displayed on the left and null cells on the right of each diagram, and wild-type cells were downsampled to match the number of null cells. Cells not belonging to the dILA2 family are plotted in gray. Solid and dashed white lines indicate approximate anatomical boundaries of the spinal cord. (B) The fraction of dILA2 family subtypes of all neurons in *Zic3*-null spinal cords is depicted here compared to wild-type and *Zic1/4*-null data, gathered from predicted neuron subtype identities in Xenium spatial transcriptomics analyses. Wild-type and *Zic1/4*-null data are depicted in Figure 5K, and are displayed here again for reference. Vertical black dashed lines connect data points to their corresponding statistical significance of comparisons at the top of the plot. (C) RNA Scope in-situ hybridization was performed on E16.5 wild-type matched littermate and *Zic3*-null spinal cords, showing qualitatively normal dILA neuron family location and numbers. For each genotype, a column of images showing a full spinal cord section and a column from a magnified region containing example dIL family subtypes are shown, with the field of view of the second column derived from the region within the box drawn with dashed white lines. White arrows indicate examples of dIL family subtypes. Other solid and dashed white lines indicate approximate anatomical boundaries of the spinal cord.
Numbers: Spinal cord sections from the same E16.5 embryos were used for both spatial transcriptomics and in-situ hybridization: n=3 wild-type, n=3 *Zic3*-null and n=3 *Zic1/4*$^{DKO}$ with an additional five wild-type and *Zic3*-null embryos examined for in-situ hybridization alone (n=8 wild-type, n=8 *Zic3*-null). For spatial transcriptomics, numbers of E16.5 wild-type brachial neurons are noted in Figure S9, downsampled to 13,356 neurons of which 6,216 were classified as dIL. Additionally, cell coordinates were derived from two whole sections and one hemisection (N=2.5 sections) among n=3 replicate Zic3-null E16.5 spinal cords. From these sections 13,197 cells were identified as neurons and plotted, of which 6,753 were annotated as dIL neurons.
Statistics: (B) Two-way ANOVA (neuron subtype x genotype). For each neuron subtype in dILA2, Tukey's multiple comparisons test was performed for each pair of genotypes, the result of which is indicated by asterisks above the plot. **p<0.01, ***p<0.001
Scale bars: 250 µm, or 100 µm when magnified from boxes.
Abbreviations: ANOVA – Analysis of variance

Figure S18: Absence of *Zic1* and Zic4 systematically alters dIL subtype composition across families

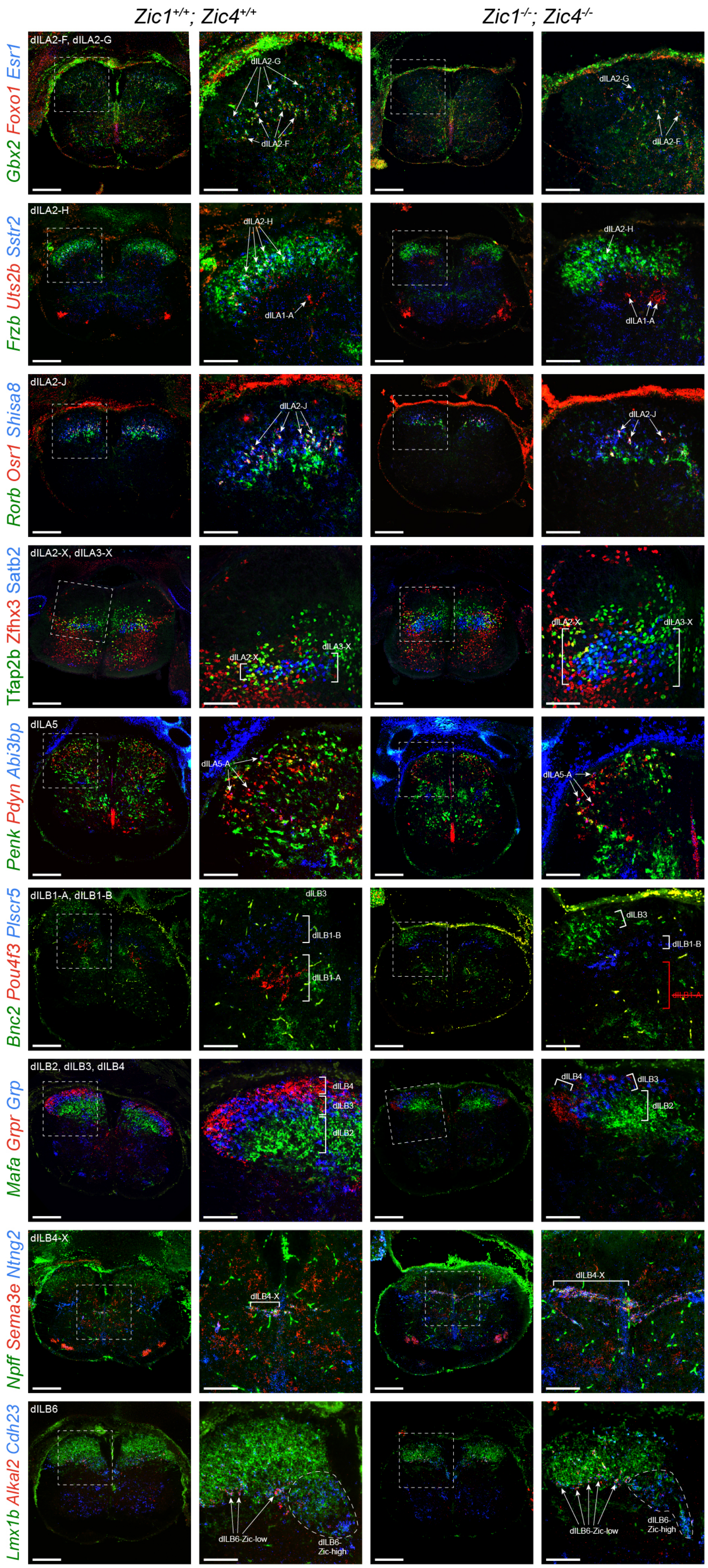

**Figure S18: Absence of *Zic1* and *Zic4* systematically alters dIL subtype composition across families, related to Figure 5.**

Images of immunohistochemistry and RNA Scope in-situ hybridization of selected dILA and dILB dorsal horn neuron types (indicated in the top left of each row) are examined in E16.5 spinal cords from wild-type (columns 1 and 2) or *Zic1/4*$^{DKO}$ embryos (columns 3 and 4). For each genotype, a low magnification images (columns 1 and 3) and a high magnification image derived from boxes marked by dashed white lines in low magnification images (columns 2 and 4) are shown. White arrows or brackets indicate examples of dIL family subtypes. Other solid and dashed white lines indicate approximate anatomical boundaries of the spinal cord. Red text or brackets indicates depletion of neuron types in DKO spinal cords.
Numbers: Images obtained from representative n=3 wild-type and n=3 *Zic1/4*$^{DKO}$ embryos.
Scale bars: 250 µm for columns 1 and 3; 100 µm for columns 2 and 4.
Abbreviations: DKO – double knock-out

Figure S19: Laminar position of dIL family subtypes in *Zic1/4*$^{DKO}$ spinal cords

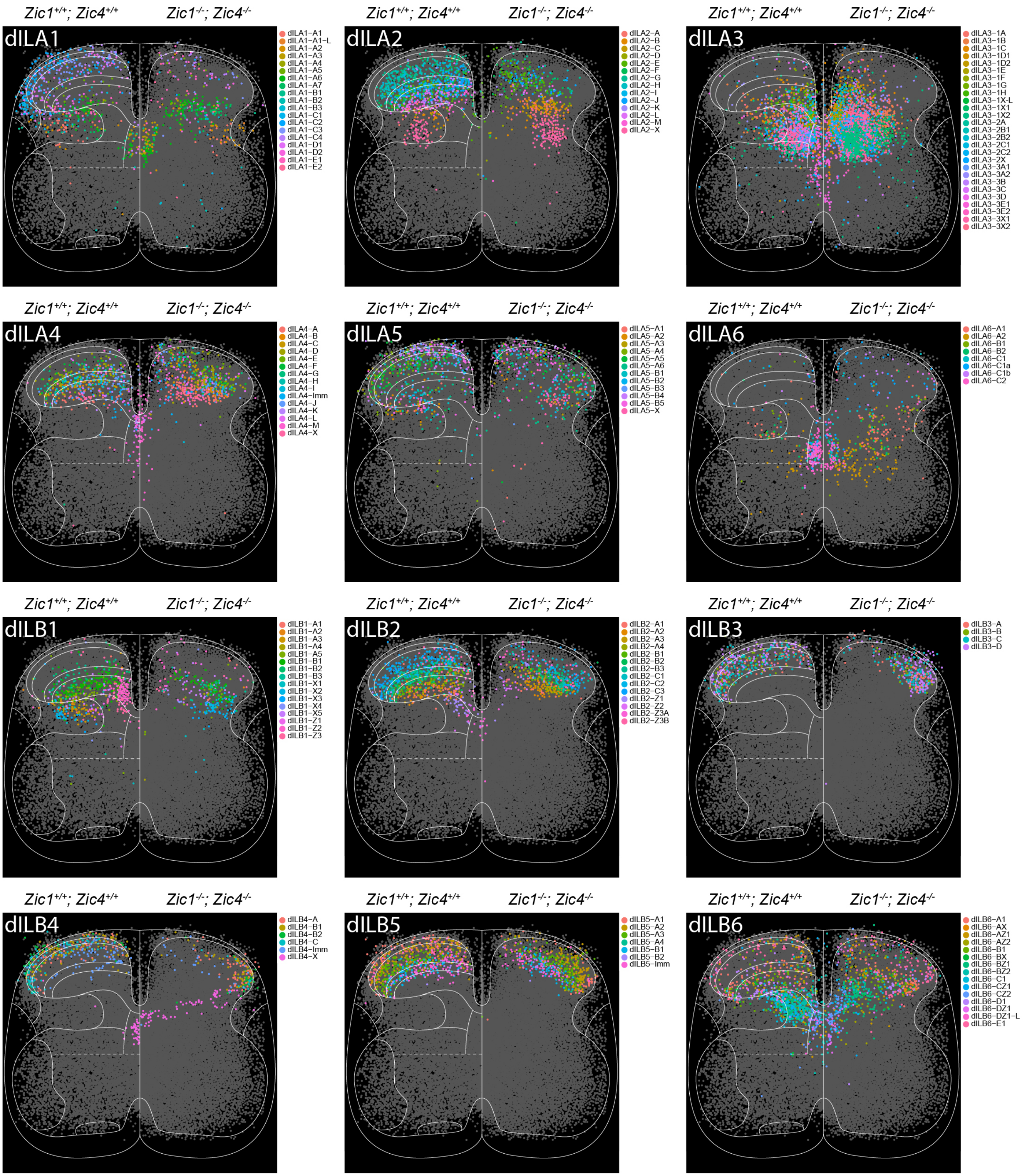

**Figure S19: Laminar position of dIL family subtypes in *Zic1/4*$^{DKO}$ spinal cords, related to Figure 5.**

Scatter plots depict the coordinates of brachial wild-type and *Zic1/4*$^{DKO}$ E16.5 neurons examined in the Xenium spatial transcriptomics assay, sorted into each of six dILA and six dILB families, and colored by subtype annotation. As described previously, wild-type cells are displayed on the left and null cells on the right of each diagram, and wild-type cells were downsampled to match the number of null cells. Cells not belonging to the indicated dIL family are plotted in gray. Solid and dashed white lines indicate approximate anatomical boundaries of the spinal cord. Numbers: Spatial transcriptomics from E16.5 wild-type embryo brachial segments was previously described (Fig. S9). For E16.5 *Zic1/4*$^{DKO}$ brachial segments, cell coordinates were derived from three whole sections and one hemisection (N=3.5 sections) among n=3 replicate spinal cords. From these sections, 17,505 cells were identified as neurons, and of which 7,376 were dIL neurons. E16.5 brachial data was downsampled to 17,347 neurons of which 8,852 were dIL neurons to match.

### Table S1: Single-cell RNA-sequencing sample statistics, related to Figure 1

| Sample | Sample index | Cells retrieved | Mean reads per cell | Mean genes per cell | nFeature_RNA > | %percent.mt < | Neurons / sample | Age | Segment | Notes |
|---|---|---|---|---|---|---|---|---|---|---|
| 14A | SI-GA-G1 | 12,634 | 41,083 | 2,678 | 500 | 5 | 2731 | E14.5 | Cervical | None |
| 14B | SI-GA-G2 | 12,363 | 41,183 | 2,918 | 500 | 5 | 3336 | E14.5 | Cervical | None |
| 14C | SI-GA-G3 | 9,576 | 46,151 | 2,359 | 500 | 5 | 1751 | E14.5 | Cervical | None |
| 14D | SI-GA-G4 | 7,945 | 48,082 | 2,448 | 500 | 5 | 2493 | E14.5 | Cervical | None |
| 14E | SI-GA-G5 | 13,317 | 34,404 | 1,594 | 500 | 5 | 2376 | E14.5 | Cervical | None |
| 14F | SI-GA-G6 | 7,507 | 51,770 | 3,713 | 500 | 5 | 2563 | E14.5 | Cervical | None |
| 16A | SI-GA-F7 | 13,350 | 45,507 | 3,059 | 500 | 5 | 3124 | E16.5 | Cervical | None |
| 16B | SI-GA-F8 | Excluded | Excluded | Excluded | Excluded | Excluded | Excluded | E16.5 | Cervical | Poor emulsion; excluded |
| 16C | SI-GA-G7 | 7,759 | 49,781 | 2,796 | 500 | 5 | 1045 | E16.5 | Cervical | None |
| 16D | SI-GA-G8 | 8,245 | 50,412 | 2,836 | 500 | 5 | 1037 | E16.5 | Cervical | None |
| 16E | SI-GA-G9 | 12,730 | 43,743 | 2,463 | 500 | 5 | 3093 | E16.5 | Cervical | None |
| 16F | SI-GA-G10 | 8,399 | 49,602 | 3,203 | 500 | 5 | 2752 | E16.5 | Cervical | None |
| 16G | SI-GA-H1 | 15,775 | 43,041 | 2,973 | 500 | 5 | 2769 | E16.5 | Cervical | None |
| 16H | SI-GA-H2 | 14,174 | 41,088 | 2,806 | 500 | 5 | 2664 | E16.5 | Cervical | None |
| 16LA | SI-GA-H3 | 11,834 | 48,483 | 3,718 | 200 | 8 | 4023 | E16.5 | Lumbosacral | None |
| 16LB | SI-GA-H4 | 35,257 | 15,364 | 1,266 | 200 | 8 | 2944 | E16.5 | Lumbosacral | None |
| 16LC | SI-GA-H5 | 7,802 | 66,608 | 4,403 | 200 | 8 | 3822 | E16.5 | Lumbosacral | None |
| 16LD | SI-GA-H6 | 8,509 | 36,892 | 3,267 | 200 | 8 | 3893 | E16.5 | Lumbosacral | None |
| 16LE | SI-GA-H7 | 7,383 | 106,417 | 5,059 | **3000** | 8 | 2615 | E16.5 | Lumbosacral | Minor emulsion failure; filters set high |
| 16LF | SI-GA-H8 | 8,258 | 65,626 | 4,334 | 200 | 8 | 3428 | E16.5 | Lumbosacral | None |
| 16LG | SI-TT-A1 | 8,858 | 57,587 | 5,112 | 200 | 8 | 3846 | E16.5 | Lumbosacral | None |
| 16LH | SI-TT-A2 | 9,583 | 63,766 | 4,999 | 200 | 8 | 3673 | E16.5 | Lumbosacral | None |
| 16TA | SI-GA-E12 | 993 | 178,101 | 730 | 500 | 5 | 32 | E16.5 | Thoracic | None |
| 16TB | SI-GA-F10 | 10,666 | 51,835 | 4,624 | **2500** | 5 | 4550 | E16.5 | Thoracic | Minor emulsion failure; filters set high |
| 16TC | SI-GA-F11 | 16,619 | 31,028 | 3,876 | 500 | 5 | 7114 | E16.5 | Thoracic | None |
| 16TD | SI-GA-F12 | 15,928 | 32,305 | 3,790 | 500 | 5 | 6994 | E16.5 | Thoracic | None |
| 16TE | SI-TT-A3 | 29,398 | 28,327 | 1,623 | 500 | 5 | 4218 | E16.5 | Thoracic | None |
| 16TF | SI-TT-A4 | 6,598 | 111,576 | 6,382 | 500 | 5 | 4659 | E16.5 | Thoracic | None |
| 16TG | SI-TT-A5 | 7,840 | 94,765 | 5,769 | 200 | 8 | 3168 | E16.5 | Thoracic | None |
| 16TH | SI-TT-A6 | 8,122 | 76,001 | 5,651 | 200 | 8 | 3335 | E16.5 | Thoracic | None |

### Table S3: Neuron family statistics

| neuron types | frequency | percent composition | Mean genes per cell | Mean silhouette score | Mean LISI score |
|---|---|---|---|---|---|
| CSF-CN | 811 | 0.862325621 | 4884.347719 | 0.564226197 | 1.068983446 |
| dI1 | 3098 | 3.294062606 | 5593.61685 | 0.02699042 | 1.11943434 |
| dI2 | 3783 | 4.022414086 | 5764.700767 | -0.055296513 | 1.103688505 |
| dI3 | 1604 | 1.705512079 | 5677.580424 | 0.208083592 | 1.119244123 |
| dI4 | 819 | 0.870831916 | 5523.943834 | 0.22369405 | 1.316595595 |
| dI5 | 709 | 0.753870364 | 6353.574048 | 0.19264621 | 1.127166402 |
| dI6 | 2617 | 2.78262164 | 5539.52159 | -0.013385167 | 1.246705571 |
| dILA1 | 3668 | 3.900136101 | 4889.60578 | 0.086811215 | 1.034098062 |
| dILA2 | 8516 | 9.054950663 | 4559.639854 | 0.11291544 | 1.050834646 |
| dILA3 | 8695 | 9.245279006 | 5023.275216 | 0.0328348 | 1.053900842 |
| dILA4 | 4998 | 5.314307588 | 4691.958183 | 0.203464403 | 1.051976747 |
| dILA5 | 5052 | 5.371725077 | 4250.411322 | 0.30324228 | 1.041783033 |
| dILA6 | 4632 | 4.925144607 | 4335.858592 | 0.215672939 | 1.089141818 |
| dILB1 | 2667 | 2.835785982 | 5618.602925 | 0.200086619 | 1.06214202 |
| dILB2 | 6304 | 6.702960191 | 4299.443687 | 0.327612298 | 1.020879822 |
| dILB3 | 4044 | 4.29993195 | 4275.0591 | 0.444091959 | 1.029691992 |
| dILB4 | 1733 | 1.84267608 | 4914.615118 | 0.219520476 | 1.1210689 |
| dILB5 | 4851 | 5.158004423 | 4677.446506 | 0.363075049 | 1.04413541 |
| dILB6 | 6284 | 6.681694454 | 4478.111871 | 0.069308565 | 1.062557291 |
| HB9-Int | 78 | 0.082936373 | 5981.525641 | 0.427726905 | 1.108151671 |
| hindbrain | 290 | 0.308353181 | 3927.024138 | 0.111834411 | Not calculated |
| Immature | 5037 | 5.355775774 | 4094.726424 | 0.020971014 | Not calculated |
| MN | 794 | 0.844249745 | 6991.423174 | 0.273319801 | 1.018233824 |
| other | 108 | 0.114834978 | 6290.324074 | 0.228359563 | Not calculated |
| v0c | 233 | 0.247745832 | 6480.321888 | 0.119235925 | 1.279123907 |
| v0d | 1410 | 1.499234433 | 5945.839716 | 0.010501009 | 1.388347551 |
| v0v | 1171 | 1.245108881 | 5848.221178 | -0.001023449 | 1.535806909 |
| v1 | 3785 | 4.02454066 | 5796.785205 | 0.011365893 | 1.163589215 |
| v2a | 2011 | 2.13826982 | 5434.530085 | 0.060594438 | 1.081866375 |
| v2b | 2169 | 2.306269139 | 5348.930844 | 0.019703151 | 1.08873846 |
| v3 | 2077 | 2.208446751 | 5778.061146 | 0.006636804 | 1.199208707 |

#### Table S5: Centroid-centroid distance between Russ et al. (2021) and Roome et al. annotations, related to Figure 1

| | CSF-cN | MN | Npy | Rorb | Ml | Cdh3 | Pdyn | Megf11 | Maf | Cpne4 | Prkcg | Reln | Rreb1 | Sox5 | ME/Lmx1b | ME | VE | Chat | Vl |
|---|---|---|---|---|---|---|---|---|---|---|---|---|---|---|---|---|---|---|---|
| CSF-CN | 8.54960379 | 52.5809004 | 49.0890971 | 49.9099976 | 47.9810616 | 49.7906775 | 49.9441726 | 49.1898147 | 49.6647114 | 51.5704406 | 49.4072513 | 51.7448462 | 49.0895174 | 49.5589128 | 47.8428771 | 47.4912426 | 48.5379813 | 49.6881691 | 48.4940819 |
| MN | 45.6336675 | 0 | 29.083821 | 31.0961545 | 26.5862455 | 31.2063677 | 32.2539551 | 27.5117056 | 30.4893623 | 33.5496639 | 30.3856428 | 33.8213392 | 28.1150185 | 29.9260339 | 26.0733221 | 23.6063927 | 23.5469024 | 25.5438632 | 24.1396341 |
| dILA1 | 41.4278342 | 28.1518935 | 3.03756802 | 16.6036074 | 11.6965996 | 18.9396917 | 18.8140188 | 16.3470503 | 19.4929894 | 23.4741737 | 19.2561145 | 23.5763896 | 16.6199657 | 18.7494493 | 15.339512 | 14.5522368 | 14.2541599 | 16.6624515 | 14.5889035 |
| dILA2 | 42.8673465 | 31.4292082 | 17.069349 | 2.22983405 | 15.9406834 | 16.3928109 | 20.2651738 | 20.9738123 | 20.9584552 | 24.2443113 | 20.8772348 | 24.9578517 | 20.2944981 | 21.1821831 | 19.879464 | 19.7579267 | 20.1002213 | 20.8377523 | 20.2487574 |
| dILA3 | 41.0010805 | 27.2487874 | 14.3823935 | 16.0288325 | 5.6890549 | 15.5805554 | 18.6576728 | 16.2418981 | 18.9309054 | 23.5543941 | 18.8918378 | 24.0777774 | 15.9608446 | 18.6061017 | 13.8779885 | 12.4081075 | 12.1708268 | 15.0881691 | 12.1143217 |
| dILA4 | 43.4270087 | 32.670671 | 21.0761392 | 16.6831867 | 18.9618715 | 3.65751389 | 20.1305363 | 23.6214957 | 22.5959814 | 26.3355931 | 22.9877163 | 27.4852563 | 22.7171356 | 23.393549 | 21.8660161 | 21.8761961 | 22.7003962 | 21.6832822 | 22.4654949 |
| dILA5 | 42.9815725 | 32.8653077 | 18.8128243 | 19.7135077 | 17.7834826 | 18.9240998 | 2.35754052 | 23.0024697 | 23.1321124 | 26.0460336 | 22.828247 | 25.8049896 | 20.4891794 | 21.8234554 | 20.2145343 | 20.6806401 | 22.230639 | 20.7189203 | 22.5654456 |
| dILA6 | 40.8575553 | 32.555468 | 20.9045313 | 22.8174862 | 14.9850966 | 22.3134626 | 19.5684024 | 23.0590764 | 22.6760426 | 26.0678447 | 21.9919795 | 26.416491 | 20.2866062 | 21.8938187 | 16.9750667 | 17.6262615 | 21.1706092 | 21.918081 | 21.3039067 |
| dILB1 | 42.3304096 | 27.8503936 | 18.5530342 | 21.5465752 | 16.6628092 | 22.8976937 | 23.6986743 | 2.34983414 | 18.7527987 | 23.9541791 | 19.9043915 | 24.7257858 | 16.6331449 | 19.4580231 | 14.2415595 | 15.3764243 | 13.9699977 | 19.8726 | 16.3073982 |
| dILB2 | 43.9060451 | 33.1727388 | 23.0856271 | 23.2520794 | 21.6710158 | 24.1059758 | 25.3517808 | 21.1791207 | 6.19161868 | 7.15464416 | 6.66832826 | 20.5248389 | 20.2461056 | 19.729063 | 20.4722375 | 21.744041 | 22.6383438 | 24.3746754 | 23.7609459 |
| dILB3 | 46.7725827 | 36.8772577 | 27.0925551 | 27.4174727 | 26.8500941 | 29.3621919 | 28.4874781 | 27.1697667 | 24.0527426 | 20.0516973 | 22.5064483 | 4.48024751 | 22.4541439 | 22.2575189 | 25.5466143 | 26.7929781 | 27.7516952 | 29.6477029 | 29.0519126 |
| dILB4 | 43.2390029 | 30.3662369 | 19.9232165 | 21.7882655 | 18.5767295 | 23.3000096 | 22.2527684 | 18.4490102 | 19.7031556 | 22.0616829 | 19.0308362 | 17.9112992 | 5.55431904 | 11.9134629 | 16.9363697 | 17.8854592 | 18.6065928 | 21.9096938 | 20.5249007 |
| dILB5 | 45.2262139 | 34.2887648 | 24.4709358 | 25.0430995 | 23.621684 | 26.3186448 | 25.7665531 | 23.6584326 | 21.9305946 | 23.7605892 | 20.7184871 | 22.2478756 | 17.1638005 | 7.25047143 | 22.3402134 | 23.3193416 | 24.2457105 | 26.2577336 | 25.7388231 |
| dILB6 | 40.4916931 | 29.2773639 | 18.2218195 | 20.5234967 | 14.32653 | 20.7778323 | 20.1555211 | 15.5711024 | 17.5326779 | 21.8453234 | 16.9713189 | 22.4701789 | 14.4532121 | 17.0941907 | 5.53962418 | 13.5753769 | 16.2884234 | 19.1076745 | 18.2335338 |
| dI1 | 41.3683879 | 25.2202284 | 17.6408715 | 20.8730154 | 13.3448383 | 21.8861411 | 22.8581881 | 14.7491419 | 19.8275715 | 24.1028237 | 19.3199232 | 24.2273689 | 15.8259057 | 18.5211658 | 12.1763547 | 8.93288252 | 5.619466 | 16.3504193 | 10.8321896 |
| dI2 | 40.8270286 | 23.8874358 | 17.1178247 | 19.900647 | 11.5750405 | 20.3567478 | 21.6152401 | 14.0585278 | 18.7253908 | 22.9115313 | 18.2093587 | 23.6969689 | 14.1806433 | 17.6736562 | 11.1279262 | 6.65450624 | 5.04561491 | 15.4138643 | 8.02154616 |
| dI3 | 41.3567615 | 22.4587432 | 17.5506941 | 21.0550357 | 13.2133432 | 22.0235029 | 23.075783 | 13.5014897 | 19.699662 | 24.1424178 | 19.5193595 | 24.6768688 | 16.388057 | 18.9168352 | 12.41921 | 8.5286062 | 3.93806516 | 15.8791 | 8.51346167 |
| dI4 | 41.364224 | 25.0127455 | 14.4457223 | 18.8225094 | 10.1127405 | 19.9024957 | 21.8846504 | 15.504688 | 20.1977712 | 24.5345501 | 20.2253492 | 25.1312675 | 17.0142917 | 19.349999 | 14.3089547 | 11.0162338 | 7.19281492 | 13.3341636 | 4.89350445 |
| dI5 | 42.0223606 | 21.9428681 | 18.0507349 | 22.0033635 | 15.5737376 | 22.472631 | 23.6783375 | 13.0692666 | 19.7310424 | 23.6609687 | 19.8968223 | 24.2270961 | 15.2916283 | 18.4569489 | 12.8401798 | 10.9503783 | 8.66366776 | 16.611516 | 12.2780805 |
| dI6 | 41.3419646 | 24.8064699 | 17.2511452 | 20.2596013 | 11.797873 | 20.3689391 | 21.488667 | 17.2393958 | 20.4214289 | 24.5674294 | 20.3864027 | 25.4604542 | 18.0395305 | 20.2298897 | 15.0059919 | 11.7342916 | 10.5974566 | 6.43586969 | 7.89471402 |
| v0d | 40.9225096 | 23.9825438 | 17.3568584 | 20.5201042 | 10.7431926 | 20.5252061 | 22.1007619 | 16.1772148 | 20.176944 | 24.6636571 | 20.3394326 | 25.5734546 | 17.8055322 | 19.9994169 | 14.0693718 | 9.65166097 | 7.60933101 | 12.1094188 | 2.80294167 |
| v0v | 40.9977822 | 22.9172206 | 17.9575462 | 20.6857638 | 11.4880778 | 20.9591261 | 22.2906587 | 14.8031106 | 19.3591246 | 23.9973066 | 19.3304994 | 24.7116783 | 16.1524291 | 18.5726627 | 12.4208298 | 6.68369809 | 4.54182698 | 14.9032256 | 7.02559577 |
| v0c | 42.8492789 | 17.2930549 | 20.8264336 | 24.4815515 | 18.4649136 | 24.7085655 | 25.965537 | 19.0557532 | 23.6763828 | 27.4552963 | 23.5206497 | 27.7074123 | 19.7765503 | 22.2044209 | 17.3497484 | 13.2442541 | 12.78016 | 18.1283168 | 14.3132827 |
| v1 | 41.4495143 | 24.6437634 | 17.4431941 | 20.623394 | 11.6798014 | 20.9235829 | 22.7600166 | 16.4401612 | 20.7782021 | 25.1836195 | 20.814922 | 26.1541412 | 18.0085129 | 20.3635709 | 14.7018496 | 10.0296641 | 7.11404339 | 15.1498682 | 3.24019325 |
| v2a | 42.4290214 | 26.0009364 | 20.7998076 | 24.0061984 | 17.1877034 | 24.2370078 | 25.1333495 | 19.2436798 | 22.4845085 | 26.5896956 | 22.2474517 | 26.9052849 | 19.8199615 | 21.897907 | 16.7880065 | 11.1623001 | 12.4330551 | 19.2974954 | 14.9216036 |
| v2b | 36.072427 | 25.4448519 | 17.6722473 | 20.616754 | 12.4721576 | 20.8310576 | 22.4912051 | 16.9082173 | 20.8260616 | 25.1036658 | 20.6317984 | 25.7440999 | 18.367253 | 20.372484 | 15.0397816 | 11.3504358 | 10.2588866 | 12.4752545 | 7.5745812 |
| v3 | 40.4249155 | 23.5586361 | 18.5803565 | 21.5570879 | 12.7844508 | 21.9085529 | 22.8988005 | 15.8837827 | 20.0807913 | 24.7506824 | 19.9490066 | 25.0859743 | 16.9831207 | 19.2218854 | 13.3258481 | 7.19756389 | 7.13657987 | 16.5420739 | 10.6530484 |
| HB9-Int | 41.0942085 | 19.0725445 | 19.1981772 | 23.4572315 | 17.1425745 | 24.5152073 | 25.0056821 | 17.7012354 | 21.9220702 | 25.9452947 | 21.1423882 | 26.1689017 | 18.4900977 | 21.07249 | 16.4578536 | 10.963924 | 10.3436505 | 18.4686042 | 12.9102759 |

#### Table S9: Xenium gene panel, related to Figure 3

| Genes | Ensembl_ID | Num_Probes | Codewords | Annotation | Panel |
|---|---|---|---|---|---|
| 2600014E21[illegible] | ENSMUSG00 | 8 | NA | NA | XED3D3 |
| 4933411E02[illegible] | ENSMUSG00 | 8 | NA | NA | XED3D3 |
| Abi3bp | ENSMUSG00 | 8 | NA | NA | XED3D3 |
| Alkal2 | ENSMUSG00 | 8 | NA | NA | XED3D3 |
| Barhl1 | ENSMUSG00 | 8 | NA | NA | XED3D3 |
| Bhlhe23 | ENSMUSG00 | 8 | NA | NA | XED3D3 |
| Bnc2 | ENSMUSG00 | 8 | NA | NA | XED3D3 |
| C1ql1 | ENSMUSG00 | 6 | NA | NA | XED3D3 |
| C1ql3 | ENSMUSG00 | 8 | NA | NA | XED3D3 |
| Calcr | ENSMUSG00 | 8 | NA | NA | XED3D3 |
| Cdh23 | ENSMUSG00 | 8 | NA | NA | XED3D3 |
| Dmbx1 | ENSMUSG00 | 8 | NA | NA | XED3D3 |
| Dmrt3 | ENSMUSG00 | 8 | NA | NA | XED3D3 |
| Drd1 | ENSMUSG00 | 8 | NA | NA | XED3D3 |
| En1 | ENSMUSG00 | 7 | NA | NA | XED3D3 |
| Esr1 | ENSMUSG00 | 8 | NA | NA | XED3D3 |
| Etv1 | ENSMUSG00 | 8 | NA | NA | XED3D3 |
| Evx1 | ENSMUSG00 | 8 | NA | NA | XED3D3 |
| Foxa2 | ENSMUSG00 | 8 | NA | NA | XED3D3 |
| Foxb1 | ENSMUSG00 | 8 | NA | NA | XED3D3 |
| Foxd3 | ENSMUSG00 | 1 | NA | NA | XED3D3 |
| Foxo1 | ENSMUSG00 | 8 | NA | NA | XED3D3 |
| Gal | ENSMUSG00 | 6 | NA | NA | XED3D3 |
| Gata3 | ENSMUSG00 | 8 | NA | NA | XED3D3 |
| Gbx1 | ENSMUSG00 | 8 | NA | NA | XED3D3 |
| Gbx2 | ENSMUSG00 | 8 | NA | NA | XED3D3 |
| Gm6213 | ENSMUSG00 | 8 | NA | NA | XED3D3 |
| Grp | ENSMUSG00 | 6 | NA | NA | XED3D3 |
| Grpr | ENSMUSG00 | 8 | NA | NA | XED3D3 |
| Il33 | ENSMUSG00 | 8 | NA | NA | XED3D3 |
| Irx2 | ENSMUSG00 | 8 | NA | NA | XED3D3 |
| Irx4 | ENSMUSG00 | 8 | NA | NA | XED3D3 |
| Isl1 | ENSMUSG00 | 8 | NA | NA | XED3D3 |
| Islr2 | ENSMUSG00 | 8 | NA | NA | XED3D3 |
| Krt71 | ENSMUSG00 | 8 | NA | NA | XED3D3 |
| Lbx1 | ENSMUSG00 | 7 | NA | NA | XED3D3 |
| Lhx1 | ENSMUSG00 | 8 | NA | NA | XED3D3 |
| Lhx2 | ENSMUSG00 | 8 | NA | NA | XED3D3 |
| Lhx9 | ENSMUSG00 | 8 | NA | NA | XED3D3 |
| Lmx1b | ENSMUSG00 | 8 | NA | NA | XED3D3 |
| Lpl | ENSMUSG00 | 8 | NA | NA | XED3D3 |
| Maf | ENSMUSG00 | 8 | NA | NA | XED3D3 |
| Mafa | ENSMUSG00 | 3 | NA | NA | XED3D3 |
| Mafb | ENSMUSG00 | 8 | NA | NA | XED3D3 |
| Mcoln3 | ENSMUSG00 | 8 | NA | NA | XED3D3 |
| Mnx1 | ENSMUSG00 | 8 | NA | NA | XED3D3 |
| Neurod2 | ENSMUSG00 | 5 | NA | NA | XED3D3 |
| Neurog2 | ENSMUSG00 | 8 | NA | NA | XED3D3 |
| Nfib | ENSMUSG00 | 8 | NA | NA | XED3D3 |
| Nkx2-2 | ENSMUSG00 | 7 | NA | NA | XED3D3 |
| Nms | ENSMUSG00 | 8 | NA | NA | XED3D3 |
| Nmu | ENSMUSG00 | 6 | NA | NA | XED3D3 |
| Nos1 | ENSMUSG00 | 8 | NA | NA | XED3D3 |
| Npas1 | ENSMUSG00 | 4 | NA | NA | XED3D3 |
| Nr4a2 | ENSMUSG00 | 8 | NA | NA | XED3D3 |
| Nr5a2 | ENSMUSG00 | 8 | NA | NA | XED3D3 |
| Ntng2 | ENSMUSG00 | 8 | NA | NA | XED3D3 |
| Nxph2 | ENSMUSG00 | 8 | NA | NA | XED3D3 |
| Olig3 | ENSMUSG00 | 8 | NA | NA | XED3D3 |
| Onecut1 | ENSMUSG00 | 8 | NA | NA | XED3D3 |
| Osr1 | ENSMUSG00 | 8 | NA | NA | XED3D3 |
| Otp | ENSMUSG00 | 8 | NA | NA | XED3D3 |
| Pax2 | ENSMUSG00 | 8 | NA | NA | XED3D3 |
| Pax5 | ENSMUSG00 | 8 | NA | NA | XED3D3 |
| Phox2a | ENSMUSG00 | 4 | NA | NA | XED3D3 |
| Pitx2 | ENSMUSG00 | 8 | NA | NA | XED3D3 |
| Pkd2l1 | ENSMUSG00 | 8 | NA | NA | XED3D3 |
| Plscr5 | ENSMUSG00 | 8 | NA | NA | XED3D3 |
| Pou4f1 | ENSMUSG00 | 8 | NA | NA | XED3D3 |
| Pou4f2 | ENSMUSG00 | 8 | NA | NA | XED3D3 |
| Pou4f3 | ENSMUSG00 | 8 | NA | NA | XED3D3 |
| Pou6f2 | ENSMUSG00 | 6 | NA | NA | XED3D3 |
| Pth2r | ENSMUSG00 | 8 | NA | NA | XED3D3 |
| Rasgrp1 | ENSMUSG00 | 8 | NA | NA | XED3D3 |
| Rxrg | ENSMUSG00 | 8 | NA | NA | XED3D3 |
| Shox2 | ENSMUSG00 | 8 | NA | NA | XED3D3 |
| Sim1 | ENSMUSG00 | 8 | NA | NA | XED3D3 |
| Skor2 | ENSMUSG00 | 8 | NA | NA | XED3D3 |
| Slc18a3 | ENSMUSG00 | 7 | NA | NA | XED3D3 |
| Slc32a1 | ENSMUSG00 | 7 | NA | NA | XED3D3 |
| Sox1 | ENSMUSG00 | 8 | NA | NA | XED3D3 |
| Sox6 | ENSMUSG00 | 8 | NA | NA | XED3D3 |
| Sp8 | ENSMUSG00 | 8 | NA | NA | XED3D3 |
| Sp9 | ENSMUSG00 | 8 | NA | NA | XED3D3 |
| Syt10 | ENSMUSG00 | 8 | NA | NA | XED3D3 |
| Tac1 | ENSMUSG00 | 8 | NA | NA | XED3D3 |
| Tac2 | ENSMUSG00 | 5 | NA | NA | XED3D3 |
| Tacr3 | ENSMUSG00 | 8 | NA | NA | XED3D3 |
| Tfap2a | ENSMUSG00 | 8 | NA | NA | XED3D3 |
| Tfap2b | ENSMUSG00 | 8 | NA | NA | XED3D3 |
| Tnfaip8 | ENSMUSG00 | 8 | NA | NA | XED3D3 |
| Tox2 | ENSMUSG00 | 8 | NA | NA | XED3D3 |
| Vsx2 | ENSMUSG00 | 8 | NA | NA | XED3D3 |
| Wt1 | ENSMUSG00 | 8 | NA | NA | XED3D3 |
| Zfhx3 | ENSMUSG00 | 8 | NA | NA | XED3D3 |
| Zic1 | ENSMUSG00 | 8 | NA | NA | XED3D3 |
| Zic2 | ENSMUSG00 | 8 | NA | NA | XED3D3 |
| Zic3 | ENSMUSG00 | 8 | NA | NA | XED3D3 |
| Zic4 | ENSMUSG00 | 8 | NA | NA | XED3D3 |
| Zic5 | ENSMUSG00 | 8 | NA | NA | XED3D3 |
